# Comparative analysis of early divergent land plants and construction of DNA tools for hyper-expression in *Marchantia* chloroplasts

**DOI:** 10.1101/2020.11.27.401802

**Authors:** Eftychios Frangedakis, Fernando Guzman-Chavez, Marius Rebmann, Kasey Markel, Ying Yu, Artemis Perraki, Sze Wai Tse, Yang Liu, Jenna Rever, Susanna Sauret-Gueto, Bernard Goffinet, Harald Schneider, Jim Haseloff

## Abstract

Chloroplast genes are present at high ploidy in plants, and capable of driving very high levels of gene expression if mRNA production and stability are properly regulated. *Marchantia polymorpha* is a simple model plant that allows rapid transformation studies, however post-transcriptional regulation in plastids is poorly characterized in this liverwort. We have mapped patterns of transcription in *Marchantia* chloroplasts. Furthermore, we have obtained and compared sequences from 51 early-divergent plant species, and identified putative sites for pentatricopeptide repeat protein binding that are thought to play important roles in mRNA stabilisation. Candidate binding sites were tested for their ability to confer high levels of reporter gene expression in *Marchantia* chloroplasts, and levels of protein production and effects on growth were measured in homoplasmic transformed plants. We have produced novel DNA tools for protein hyper-expression in a facile plant system that is a test-bed for chloroplast engineering.

## INTRODUCTION

Chloroplasts are the semi-autonomous organelles responsible for the capture of light energy through the conversion of CO_2_ to organic molecules in plants. Chloroplast (plastid) genomes are small and highly conserved, present at high copy number per cell, and not subject to gene silencing. Foreign proteins have been produced in chloroplasts at high levels, sometimes reaching a major proportion of the total soluble proteins in transformed plants (Cosa et al. 2001; Oey et al. 2009; Kanamoto et al. 2006). However, previous attempts to harness this capacity for routine hyper-expression (>1% soluble protein) have been irregular and sporadic. The primary reasons for this lack of application are the relatively small number of species with established methods for chloroplast transformation, the slow pace and inefficiency of plastid transformation, and the inconsistent levels of gene expression between experiments.

Past attempts to build more efficient vectors for chloroplast gene expression have focused on increasing the efficiency of transcription, translation initiation and codon usage. However, recent work has led to a breakthrough in understanding the important roles of post-transcriptional processing and mRNA stability in conferring high levels of gene expression in chloroplasts (Legen et al. 2018; Rojas et al. 2019). Plastid RNA transcripts are subject to a series of complex processing steps that are primarily mediated by nucleus-encoded factors, including pentatricopeptide repeat (PPR) containing proteins. The PPR proteins are a large family of RNA-binding proteins that have undergone a substantial expansion in plants (Barkan and Small 2014) and are required for stabilisation of mRNAs by protection from exonuclease activity in the plastid (Prikryl et al. 2011; Legen et al. 2018). The sequence-specific RNA-binding properties and defined target sites for these proteins make them excellent candidates as artificial regulators of RNA degradation, in addition to being used as highly effective tools for enhancing gene expression in chloroplasts (Legen et al. 2018; Rojas et al. 2019).

*Marchanti*a *polymorpha* is one of the few land plant species for which chloroplast transformation is well established (Boehm et al. 2016; Chiyoda, Yamato, and Kohchi 2014). *Marchantia* has a series of characteristics that make it an ideal platform for chloroplast engineering (Boehm et al. 2017). It grows rapidly through both asexual and sexual life cycles, has a remarkable regenerative capacity in the absence of phytohormones, the dominant phase of the life cycle is haploid and transplastomic plants can be isolated within 8 weeks (Sauret-Güeto et al. 2020). In addition, post-transcriptional regulation of chloroplasts mRNAs in *Marchantia* is relatively simple compared to vascular plants. For example, the *Marchantia* nuclear genome encodes 75 PPR proteins (Bowman et al. 2017) directed to chloroplast and mitochondria, while the *Arabidopsis* and rice genomes encode over 450 and 600 PPR proteins, respectively (Gutmann et al. 2020). Additionally, no evidence of PPR protein-mediated base editing has been found in *Marchantia* chloroplast transcripts (Ichinose and Sugita 2016). *Marchantia* shows great promise as a simple and facile test-bed for chloroplast engineering, but little is known of the cis-regulatory elements required to fully exploit the capacity of plastids for high and sustained levels of gene expression.

In order to exploit *Marchantia* as a testbed for chloroplast engineering, we conducted a transcriptional analysis of the *Marchantia* chloroplast, and examined an expanded range of Bryophyte plastid genomes for conserved sequences in the 5’ UTRs of highly expressed mRNAs. This study provides the first description of chloroplast transcription patterns in a liverwort, and comparisons within this under-studied group of early divergent plants. It has also produced a variety of new DNA tools that enable the generation of plants capable of hyper-expression of proteins in this facile model system.

## RESULTS

### *Marchantia* chloroplast transcriptome analysis

We previously generated a high-quality plastid genome assembly for the *M. polymorpha* Cam1/2 isolates using next generation sequencing data (Genbank accession: MH635409.1) (Sauret-Güeto et al. 2020) (Supplementary Fig. 1). We conducted this assembly to resolve a taxonomic misidentification of the source of the reference plastid genome (Genbank accession NC_001319.1), which likely originated from the related species *Marchantia paleacea* (Kijak, Łodyga, and Odrzykoski 2018). The plastid genome of *M. polymorpha* Cam1/2 is 120,314 bp and contains 123 annotated genes, which are mainly involved in photosynthesis, electron transport, transcription, and translation. A small number of genes with more specific functions are also present, such as the *chl*L gene involved in chlorophyll biosynthesis (Ueda et al. 2014).

Recent experiments have demonstrated the crucial importance of both promoter identity and adjacent 5’ untranslated regions for initiating and stabilising high levels of transcription in chloroplasts (Rojas et al. 2019; Q. Yu, Barkan, and Maliga 2019). In order to better understand which sequences might be useful for engineering high levels of gene expression, we employed differential RNA sequencing (dRNA-seq) (Sharma et al. 2010), which allowed identification of primary transcripts in extracted chloroplast RNAs. This technique was initially developed for prokaryotic organisms but has also successfully been applied to barley chloroplasts (Sharma et al. 2010; Zhelyazkova, Sharma, et al. 2012). RNAs isolated from *Marchantia* chloroplasts were treated with Terminator™ 5’ phosphate dependent exonuclease (TEX) in order to selectively degrade RNAs with 5’ monophosphate termini, while primary transcripts with 5’ triphosphate termini are resistant to degradation (Fig. 1a). Treated and untreated RNA populations were sequenced to locate transcription start sites (TSS), and putative promoter and 5’ UTR regions. The main goals of these experiments were (i) to identify highly transcribed regions of the *Marchantia* chloroplast genome, (ii) to locate transcription start sites of mRNAs that accumulate to high levels, and (iii) screen for conserved sequences that might indicate important features that could be incorporated into synthetic promoter and mRNA elements to promote high levels of protein expression.

**Fig. 1:**
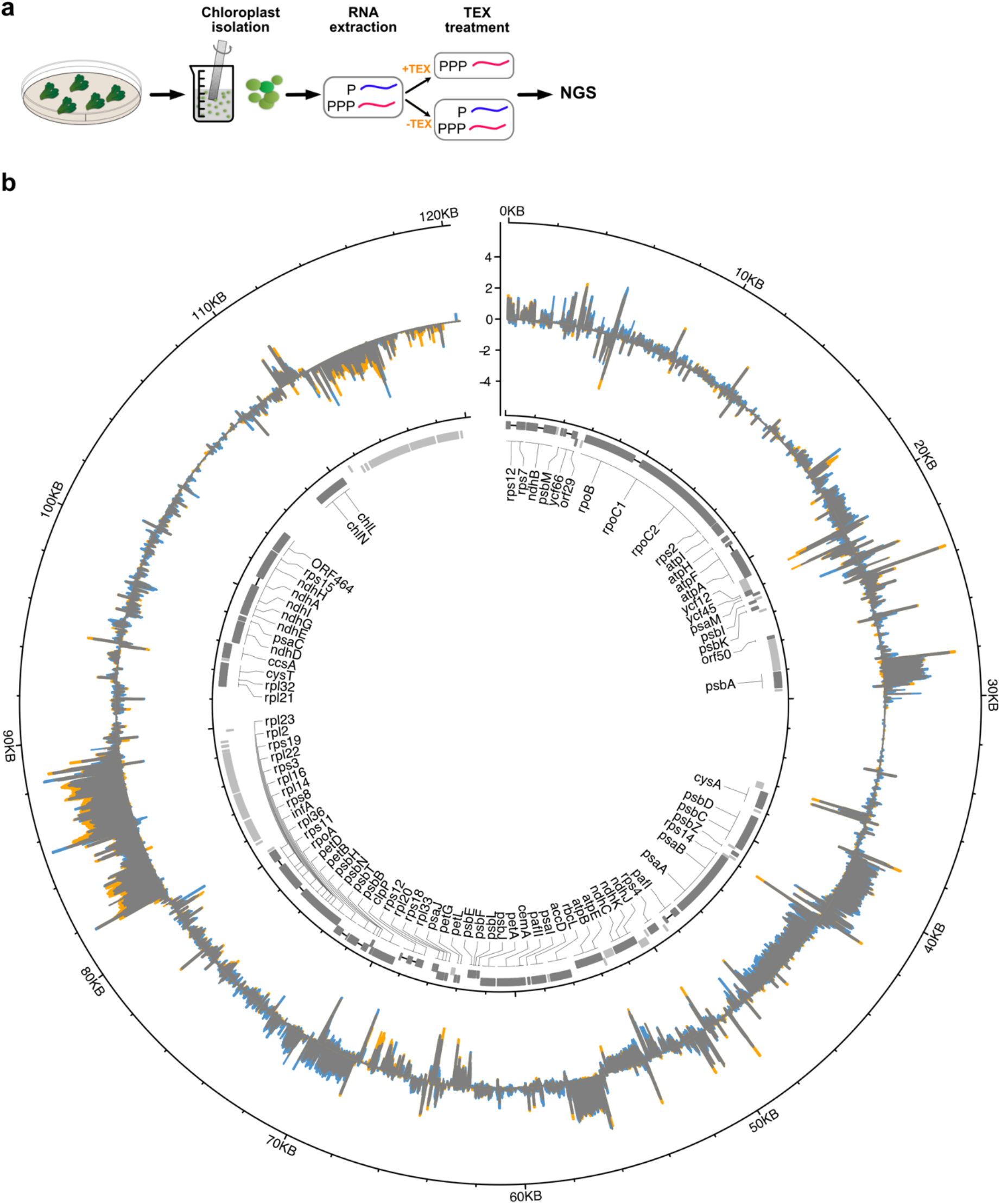
**a)** Outline of dRNAseq pipeline: The plant tissue is collected and homogenised. Intact chloroplasts are isolated from homogenised plant tissue, RNA is extracted and then subjected to treatment with the terminator exonuclease (TEX) enzyme. TEX degrades RNAs with a 5’ monophosphate (processed transcripts) but not with a 5’ triphosphate (primary transcripts). Consequently, the comparison of next generation sequencing libraries generated from TEX treated (TEX+) and non-treated (TEX-) samples can be used to identify the protected primary transcripts and their TSSs. The identification of TSS allows also the more accurate mapping of promoter regions. **b)** dRNAseq in Marchantia. Outer circle: Mapped reads of TEX treatment (TEX+ libraries) and non-enriched (TEX-libraries) mapped on *M. polymorpha* accession Cam1/2 accession plastid genome (MH635409). Forward strand coverage faces outwards, reverse strand coverage faces inwards. Y-axis: log10 coverage per million mapped reads. Blue: excess TEX-coverage (TEX-enrichment), Orange: excess TEX+ coverage (TEX+ enrichment), Grey: TEX-= TEX+. Inner circle depicts the gene organization of the Marchantia plastid genome. Protein coding genes are shown in dark grey, boxes show coding sequences and lines introns. Non-coding genes are shown as light grey boxes. Boxes for genes encoded clockwise face outwards, those encoded counterclockwise strand genes face inwards. Gene names are shown for protein coding genes in the centre.

Short sequence reads (75 bp) were obtained from TEX treated and untreated RNA samples and mapped onto the plastid genome of *M. polymorpha* accession Cam1/2 (MH635409) (Fig. 1b and Supplementary Fig.2 and Supplementary Table 1). The levels of transcript abundance could be observed. These were mapped onto different regions of the plastid genome, with evident polarity that reflected the directions of transcription across transcribed genes and operons.

We manually assigned a total of 186 potential TSSs to locations on the *Marchantia* chloroplast genome (Fig. 2a and Supplementary table 2). The identified TSSs could be grouped into four categories based on their genomic location: i) gene TSSs (gTSSs), found within a region upstream of annotated genes, ii) internal TSSs (iTSSs) found within annotated genes and giving rise to sense transcripts, iii) antisense TSSs (aTSSs) located on the opposite strand within annotated genes and giving rise to antisense transcripts, which could indicate the synthesis of non-coding RNAs; and iv) orphan TSSs (oTSSs). In total, we mapped 108 gTSSs, 40 iTSSs, 21 aTSSs and 17 oTSSs (Fig. 2a).

**Fig. 2:**
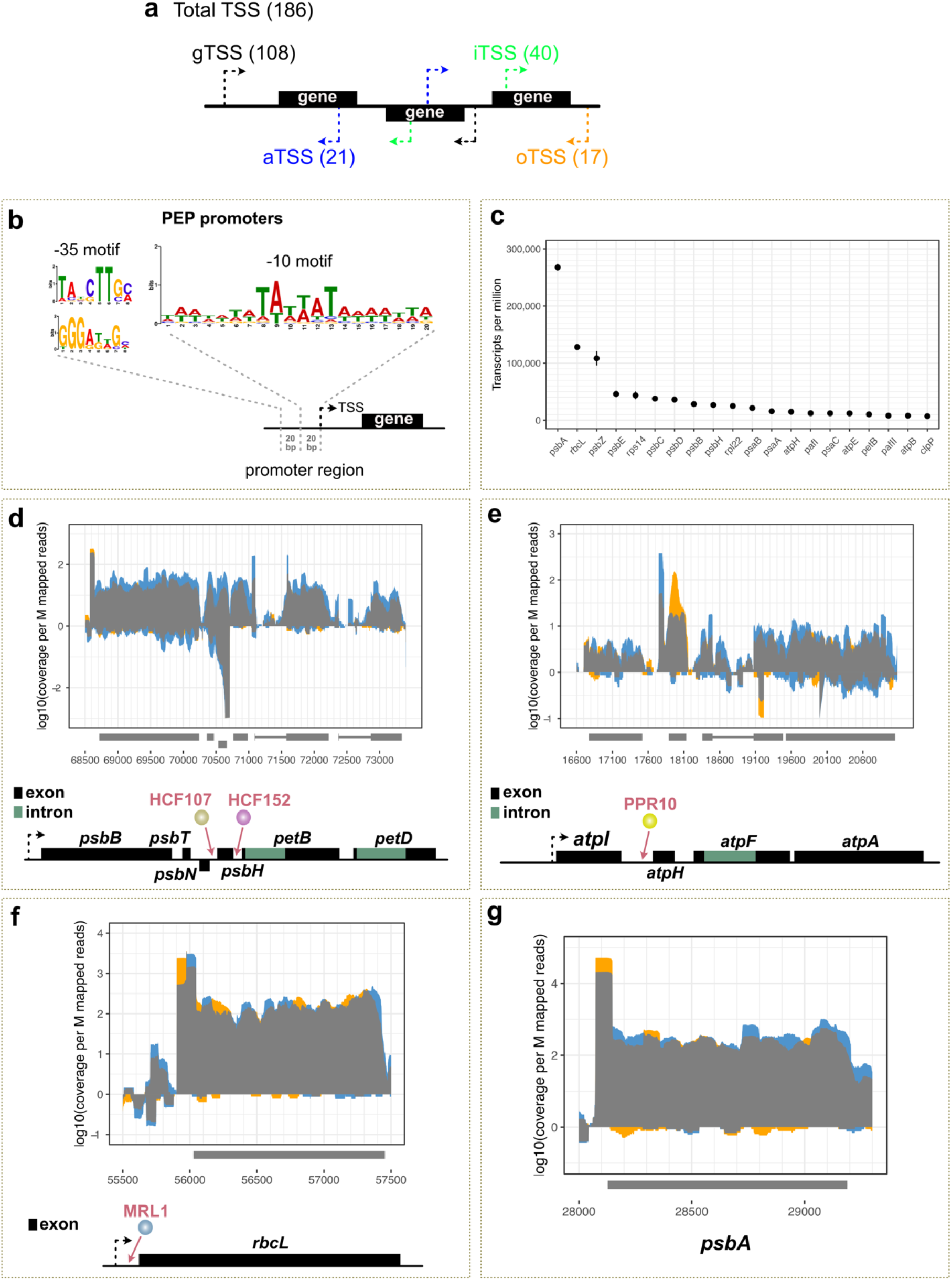
**a)** Graphical summary of different species of TSSs identified in Marchantia plastid genome using dRNAseq. A total of 186 potential TSSs, with the most abundant species associated with tRNAs. The identified TSSs could be further grouped into four categories based on their genomic location: i) gene TSSs (gTSSs), found within a region upstream of annotated genes, ii) internal TSSs (iTSSs) found within annotated genes and giving rise to sense transcripts, iii) antisense TSSs (aTSSs) located on the opposite strand within annotated genes and giving rise to antisense transcripts; and iv) orphan TSSs (oTSSs). In total 108 gTSSs, 40 iTSSs, 21 aTSSs and 17 oTSSs were mapped. **b)** MEME (Bailey et al. 2015) analysis discovered a −10 PEP consensus element upstream of 140 TSSs (e-value 5.3e-028). Two −35 PEP consensus motifs were predicted in 25 out of the 140 sequences. Top: 16 sequences (e-value: 2.5e+001) Bottom: 9 sequences (e-value: 8.4e-002). **c)** Top 20 genes, excluding tRNAs and rRNAs, with the highest expression levels (TPM) in Marchantia chloroplast. **d-g)** Primary transcript enriched (TEX+ libraries) and non-enriched (TEX-libraries) mapped on the genomic location of **d)** Mp-*psbB* operon and **e)** large Mp-*atp* operon **f)** Mp-*rbcL* and **g)** Mp-*psbA*. X-axis: genomic position. Y-axis: coverage per million of mapped reads. Blue: excess TEX-coverage (TEX-enrichment), Orange: excess TEX+ coverage (TEX+ enrichment) and Grey is TEX-= TEX+. Operon maps are depicted below the graphs **d)** The *psbB* operon comprises five genes: *psbB, psbT, psbH, petB* and *petD*. Each of the *petB* and *petD* genes contains an intron. The *psbN* gene, which is encoded in the intercistronic region between *psbH* and *psbT*, is transcribed in the opposite direction. In Marchantia we identified a TSS 144 bp upstream the *psbB* gene, 47 bp upstream the *psbN* gene, 36 bp upstream the *psbH* gene and 43 bp upstream the *petB* gene. In *Arabidopsis* the HCF152 PPR protein binds to a sequence located in the 5’ untranslated region of the *petB* chloroplast gene stabilising RNA transcripts against 5’→3’ exonuclease degradation (Meierhoff et al. 2003). The HCF107 protein binds upstream *psbH* to stabilize the *psbH* transcript and activates *psbH* translation (Felder et al. 2001). **e)** The large *atp* operon is composed of four genes: *atpI, atpH, atpF* and *atpA*. In Marchantia we identified a TSS 73 bp upstream the *atpI* gene and an internal TSS (*atpI*) 143 bp upstream the *atpH* gene. In maize the PPR10 protein binds to a sequence located in the 5’ untranslated region of the *atpH* chloroplast gene and has been found to play a role in controlling translation by defining and stabilising the termini, protecting them from exonucleases (Pfalz et al. 2009). **f)** We identified a TSS 124 bp upstream the *rbcL* gene. In *Arabidopsis* the MRL1 PPR protein binds to a sequence located in the 5’ untranslated region of the *rbcL* chloroplast gene, acting as a barrier to 5 ‘->3 ‘ degradation (Pfalz et al. 2009; Johnson et al. 2010). **g)** We identified a TSS 54 bp upstream the *psbA* gene.

The most abundant gTSSs corresponded to tRNA genes. The *Marchantia* plastid genome encodes 31 unique transfer RNAs (tRNA), five of which are present in two copies in the inverted repeat (IR) regions. Given that the genome contains only 123 genes, the number of identified TSSs exceeded expectations, especially considering that some are likely encoded in co-transcribed operons. The experimental approach can be confounded by post-transcription processing or degradation, or low abundance of primary transcripts.

### Characterisation of active promoters and transcripts

Plastid transcription is mediated by two distinct RNA polymerases: the eukaryotic nuclear encoded RNA polymerase (NEP) and the prokaryote-like plastid encoded RNA polymerase (PEP), which is retained from the cyanobacterial endosymbiont (Yagi and Shiina 2014). PEP recognises bacterial type promoters that contain conserved domains at positions −35 and −10 (TATA) (Ortelt and Link 2014), whereas NEP recognises promoters that have a core sequence “YRTA” (where Y is cytosine or thymine and R is, Guanine or Adenine) motif in close proximity to the transcription start site (Ortelt and Link 2014; Hess and Börner 1999). However, many genes can be transcribed by both. In general, PEP promoters appear to be much stronger than NEP promoters, and highly expressed genes in the plastid genome (e.g., most photosynthesis genes) are usually transcribed from PEP promoters (Ortelt and Link 2014). For this reason, PEP promoters have been predominantly used to drive the expression of plastid transgenes.

A limited number of promoters have been employed for transgene expression in chloroplasts, and mainly in systems such as tobacco and *Chlamydomonas* (Adem, Beyene, and Feyissa 2017; S. Jin and Daniell 2015). These promoters are derived from highly expressed plastid genes, such as the large subunit of ribulose-1,5-bisphosphate carboxylase/oxygenase (RuBiSco) (*rbc*L), the photosystem II protein D1 (*psb*A) gene and the plastid rRNA operon, *rrn*. Only two studies have focused on promoter regions of plastid genes in *Marchantia*: (Shimmura et al. 2017) analysed the promoter region of the *psb*D gene and (Lyubetsky, Rubanov, and Seliverstov 2010) predicted the promoter regions of *psa*A, *psb*A, *psb*B, *psb*E and *rbcL* genes based on sequence comparison of several plant species.

Studies in *Marchantia* have employed heterologous tobacco *psb*A and *prrn* promoters to drive expression of transgenes (Boehm et al. 2016; Chiyoda, Yamato, and Kohchi 2014). The identification of *Marchantia* plastid gene TSSs has allowed precise characterization of the initiation sites for transcription, and the mapping of the 5’ termini of transcripts in a wide range of genes. These newly identified elements crucially expand the repertoire of available promoter parts to be considered when designing transgenes for *Marchantia* chloroplast engineering.

The 50-nucleotide regions upstream of the identified TSSs were screened for potential promoter motifs using the Multiple Expectation maximization for Motif Elicitation (MEME) tool (Bailey et al. 2015). We found a −10 TAttaT motif located three to nine nucleotides upstream of the transcription start point for 140 predicted TSSs, similar to that found in barley (Zhelyazkova, Sharma, et al. 2012)(Supplementary Table 3). Examination of the −35 region showed a lower degree of sequence conservation than the −10 box. Two −35 motifs were mapped in only 25 out of those 140 TSSs (Fig. 2b).

To distinguish candidate DNA parts for high level gene expression, we used data from untreated dRNAseq samples and identified the 20 protein-encoding genes with the highest RNA accumulation in the *Marchantia* chloroplast. (Fig. 2c). As was predicted based on other plant models (S. Jin and Daniell 2015), the *psb*A and *rbc*L genes have the highest mRNA transcript levels in *Marchantia* chloroplasts. The dRNAseq profiles of the promoter regions of these genes were examined in more detail. The genetic maps and transcript profiles of these regions are shown in Fig. 2d-g. After TEX treatment, we observed an approximately 5-fold enrichment of reads mapped at the 5’ end of the primary transcript for *rbc*L and approximately 2.5-fold enrichment for *psb*A. The identified TSSs were located 124 bp and 54 bp upstream of the predicted start codons for *rbc*L and *psb*A, respectively. In addition, all four regions show similar gene arrangements compared to plastid genomes of other land plants. The homologous regions in vascular plants include binding sites for conserved PPR proteins that have been found to bind to chloroplast mRNAs, to confer increased RNA stability by protection against ribonucleases, and promote high levels of gene expression. The approximate locations of four potential PPR binding sites are indicated in these regions of the *Marchantia* plastid genome.

### Operons

Many chloroplast genes, often functionally-related, are organised in co-transcribed operons. Examples include the *psbB* operon and the two ATP synthase (*atp*) operons (the large *atp*I/H/F/A and the small *atp*B/E operon). Operons are usually transcribed as a unit and the transcripts processed to yield smaller monocistronic mRNAs. Operon processing is mediated by various factors that recognise particular operon non-coding sequences. These sequences harbour gene expression elements, such as PPR binding motifs, that are potentially useful for plastid engineering applications. As for promoters, the available information about operon structure and regulation in *Marchantia* is very limited.

The *psb*B operon comprises five genes encoding the photosystem II subunits CP47 (*psb*B), T (*psb*T), and H (*psb*H) as well as cytochrome b6 (*pet*B) and subunit IV (*pet*D) of the cytochrome b6f complex. In *Arabidopsis* it is initially transcribed as a large precursor mRNA, which is extensively processed (Meierhoff et al. 2003). Each of the *pet*B and *pet*D genes contains an intron, which is spliced during post-transcriptional modification. The *psb*B operon is regulated by more than one promoter (Fig. 2d). In particular, the small subunit of photosystem II (*psb*N), which is encoded in the intercistronic region between *psb*H and *psb*T, is transcribed in the opposite direction by an additional promoter. In *Marchantia* we identified a TSS 144 bp upstream of the *psb*B gene, 47 bp upstream of the *psb*N gene, 36 bp upstream the *psb*H gene and 43 bp upstream the *pet*B gene.

The large *atp* operon is composed of four genes: *atp*I, *atp*H, *atp*F and *atp*A. Plastid operons often have multiple promoters that enable a subset of genes to be transcribed within the operon (Kuroda and Maliga 2002). For example, this operon is transcribed by two PEP promoters in *Arabidopsis*, one upstream and one within the operon, and harbours four potential sites for RNA-binding proteins (Malik Ghulam et al. 2013). In *Marchantia* we identified a TSS 73 bp upstream of the *atp*I gene and an internal TSS (*atp*I) 143 bp upstream of the *atp*H gene (Fig. 2e).

### Comparisons with other bryophyte plastid genomes

Over 4,500 plastid genomes have been sequenced to date, and the overwhelming majority of these belong to angiosperm plants (Gutmann et al. 2020; Tonti-Filippini et al. 2017; Y. Yu et al. 2019). Sequence comparisons between the plastid genomes of land plants have revealed gross gene rearrangements, but individual coding regions and a number of gene clusters are recognisably conserved. In addition, certain cis-regulatory sequences, such as PPR-binding sites, are conserved and often located near the 5’ termini of mRNA transcripts (Zhelyazkova, Hammani, et al. 2012). However, the small size and apparent sequence redundancy of the domains makes them difficult to identify by comparison between divergent species. At the initial phase of our investigation, only eight bryophyte plastid genomes were publicly available. To overcome this limitation we expanded the sampling to 51 plastid genomes from bryophytes, and used comparative genomics to screen the *Marchantia* plastid genome for potential regulatory sequences.

We determined the complete sequences of 28 liverwort plastid genomes, 13 moss genomes and one hornwort genome. We also included in our analysis three recently published hornwort plastid genomes (F.-W. Li et al. 2020) and eight published liverwort plastid genome sequences (Supplementary Table 4 and 5), as well as three angiosperm plastid sequences for reference. The dataset comprised representatives of all three classes of liverworts, namely Haplomitriopsida, Marchantiopsida, and Jungermanniopsida (Söderström et al. 2016). In summary, we included representatives of six of the 15 liverwort orders, 13 of the 26 moss orders (Liu et al. 2019) and three of the five hornwort orders (Villarreal and Renner 2012) currently recognized (Fig. 3a and Table 1). Comparison of the newly generated bryophyte plastid genomes further supports the observation of a remarkable conservation of plastid genome structure among land plants (Liu et al. 2019).

**Table 1:** Sampling of land plant plastid genomes employed in this study.

| Lineages | Orders | Families | Orders sampled | Families sampled |
| --- | --- | --- | --- | --- |
| Hornworts | 5 | 11 | 3 | 3 |
| Liverworts | 15 | 87 | 7 | 21 |
| Mosses | 29 | 109 | 13 | 14 |
| Angiosperms | 64 | 418 | 3 | 4 |

**Fig. 3:**
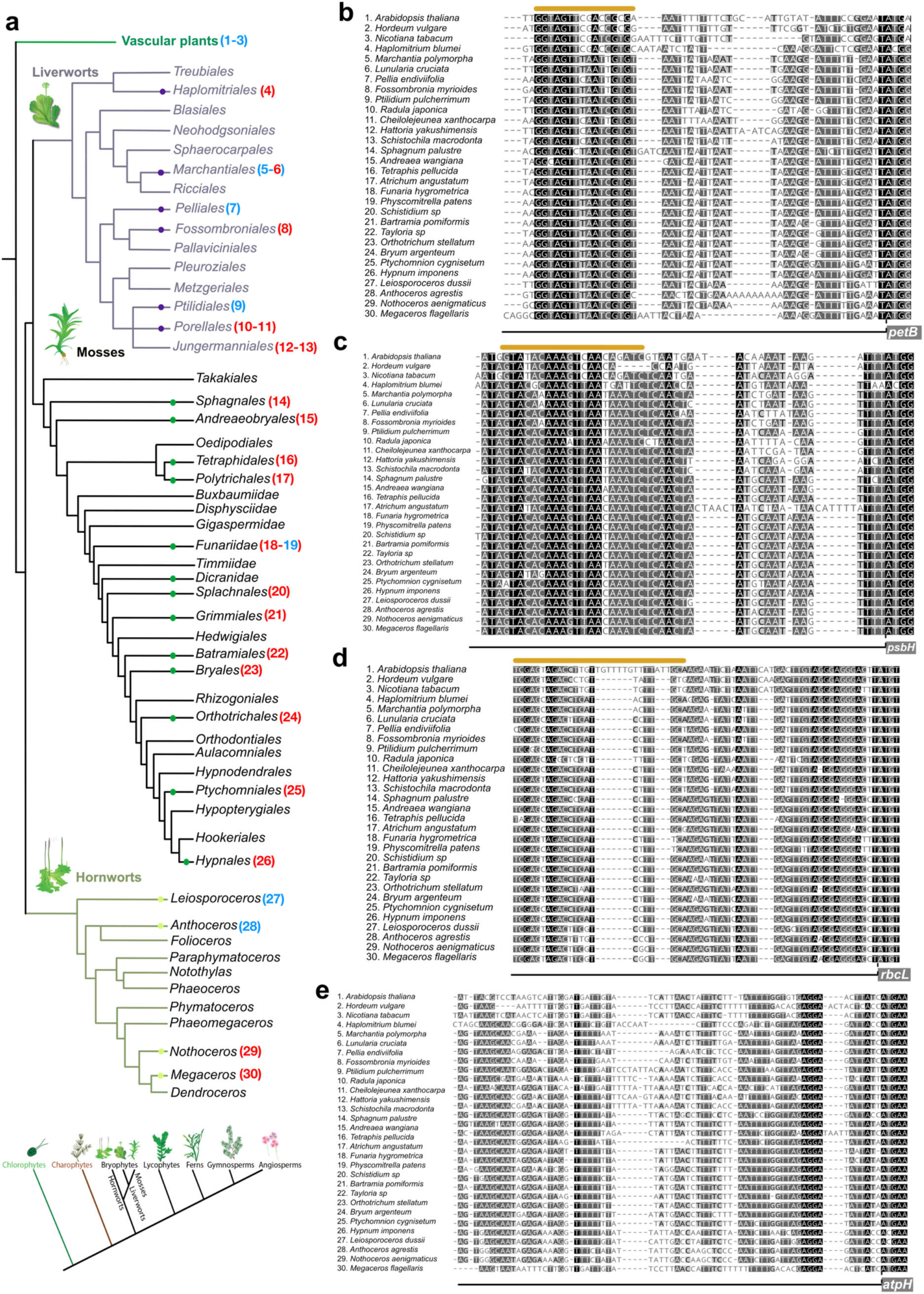
**a)** Bryophyte phylogeny modified from (Liu et al. 2019) using the most recent phylogenetic inference about the relationship of bryophytes (F.-W. Li et al. 2020). Numbers next to Order names indicate sampled species which were included in our analysis. Bottom: Land plant phylogenetic tree based on (F.-W. Li et al. 2020) with bryophytes being monophyletic and hornworts being sister to mosses and liverworts **b-e)** Multiple sequence alignments, using MUSCLE (Edgar 2004a), of upstream nucleotide sequences of *petB* **(b)**, *psbH* **(c)**, *rbcL* **(d),** and *atpH* **(e)** genes from 27 different bryophyte species and three angiosperms. Numbers next to species names correspond to the phylogenetic Order in (a). ATG site is indicated with a dashed line. Coding sequence is indicated with a grey box. The predicted PPR binding site is highlighted by an orange line above. The coloring used for that column depends on the fraction of the column that is made of letters from this group. Black: 100% similar, dark-grey 80-100% similar, lighter grey: 60%-80% similar, white: less than 60% similar.

### Identification of putative PPR protein binding sites

In order to identify conserved sequences that could be important for mRNA function in the chloroplast, we performed a phylogenetic comparison of mRNA sequences (up to ~100 bp) upstream of the predicted initiator codon of the highly expressed *pet*B, *rbc*L, *atp*H and *psb*H coding regions (Fig. 3 b-e). It is known that similar regions within the corresponding angiosperm mRNA sequences encode binding sites for specific PPR proteins. The HCF152 PPR protein binds to a sequence located in the 5’ untranslated region of the *pet*B chloroplast mRNA. It has been experimentally demonstrated that binding of the protein to RNA transcripts stabilises them against 5’→3’ riboexonuclease degradation in *Arabidopsis* (Meierhoff et al. 2003). We also included in our analysis the High Chlorophyll Fluorescence 107 (HCF107) protein, which is a member of the family of PPR proteins that contain domains similar to histone acetyltransferases (HAT). HCF107 stabilizes the *psb*H transcript and activates *psb*H translation (Felder et al. 2001). The MRL1 PPR protein binds to a sequence located in the 5’ untranslated region of the *rbc*L chloroplast gene. In *Arabidopsis*, MRL1 is necessary for the stabilization of the *rbc*L processed transcript, likely because it acts as a barrier to 5 ‘→3 ‘ degradation (Johnson et al. 2010). The PPR10 protein binds to a sequence located in the 5’ untranslated region of the *atp*H chloroplast gene and has been found to play a role in controlling translation by defining and stabilising the 5’ terminus, protecting it from exonuclease activity (Pfalz et al. 2009). The *Marchantia* nuclear genome encodes 75 PPR proteins (Pfalz et al. 2009; Bowman et al. 2017) including recognizable homologs of High Chlorophyll Fluorescence 152 (HCF152), Maturation of *rbcL*1 (MRL1), PPR10 and HCF107 (Supplementary Fig. 3).

We used the new bryophyte plastid genome alignments to search for conserved mRNA sequence motifs across both bryophyte and angiosperm plant species. Fig. 3b-e shows the alignments of 30 plastid genome segments from bryophytes and key angiosperm species (alignments for all bryophyte species used in this study in Supplementary Fig. 4). The alignments correspond to the 5’ sequences of *pet*B, *psb*H and *rbc*L and *atp*H mRNAs. The relevant PPR protein binding sites have been experimentally determined in certain angiosperms, and the binding footprints are indicated (Zhelyazkova, Hammani, et al. 2012).

These footprints coincide with conserved nucleotide sequences at the binding site. These sequences appear highly conserved across the angiosperms and bryophytes for the *Marchantia petB, psbH* and *rbc*L chloroplast mRNAs, although less so for the *atp*H mRNA.

The nucleotide sequence similarity of these putative binding sites, and existence of homologous PPR proteins in *Marchantia* suggests that the functional relationship between nuclear-encoded PPR proteins and regulation of chloroplast mRNA stability may be conserved for (at least) *pet*B*, psb*H *and rbc*L across the land plants. Further, these putative PPR protein binding sites in *Marchantia* might be transplanted into engineered chloroplast genes and confer improved mRNA stability. We built and tested hybrid gene genes to test this hypothesis.

### Creating artificial leader sequences

We have developed an open source DNA toolkit for facile engineering of nuclear and plastid genomes in *Marchantia* (Sauret-Güeto et al. 2020). The toolkit is based on Loop assembly (Pollak et al. 2019), a Type IIS method for DNA construct generation that employs a recursive strategy to greatly simplify the process of plasmid assembly. It allows rapid and efficient production of large DNA constructs from DNA parts that follow a common assembly syntax. Unlike other systems that require elaborate sets of vectors, Loop assembly requires only two sets of four complementary vectors. In a series of reactions, standardized DNA parts can be assembled into multi-transcriptional units (Fig. 4a). The system includes DNA vectors and basic parts for transformation of the *Marchantia* plastid genome (Fig. 4b).

**Fig. 4:**
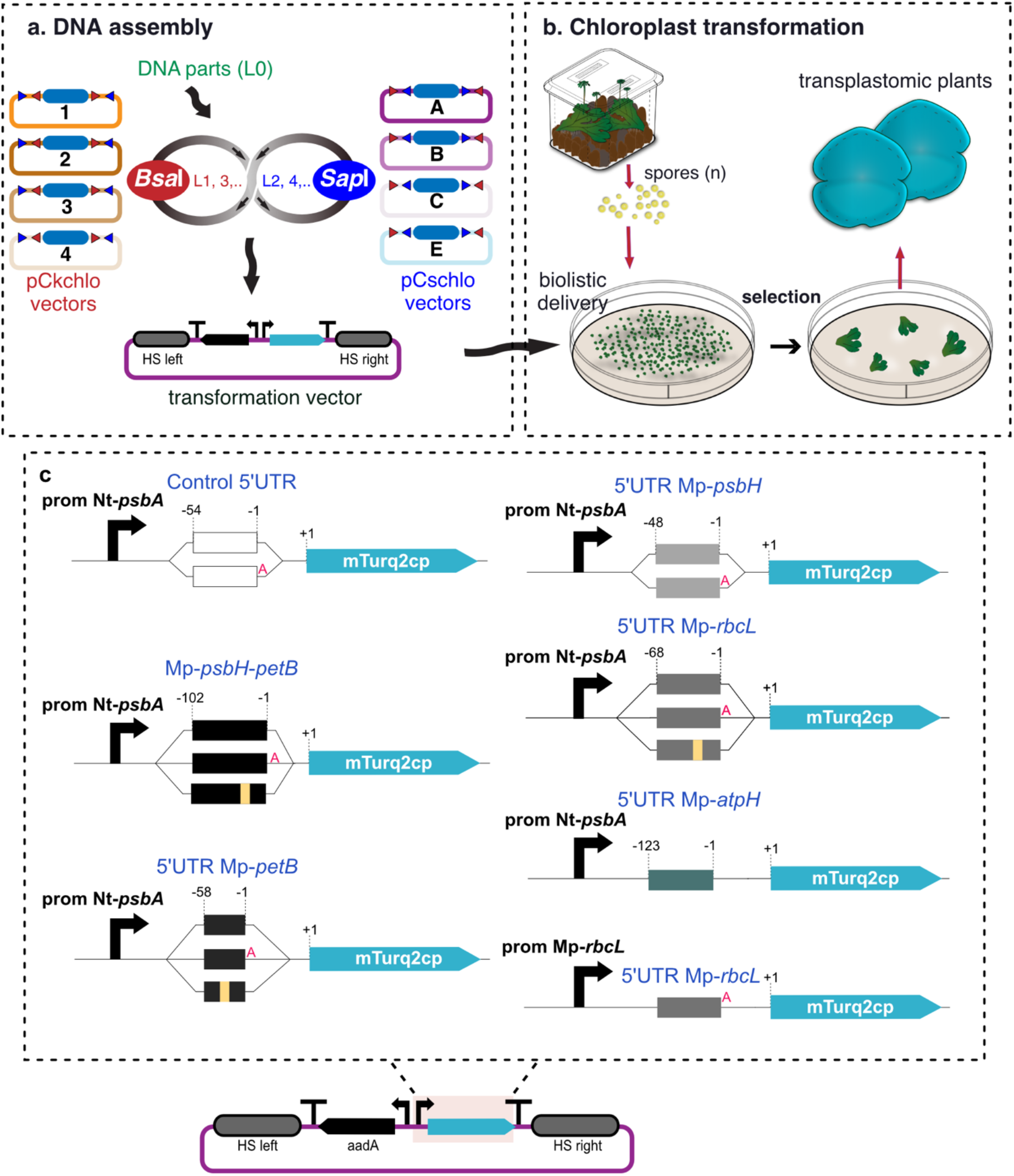
**a) Chloroplast Loop assembly overview** Level 0 (L0) DNA parts are assembled in Level 1 (L1) transcription units (TUs) into one of the four pCkchlo vectors, depicted with numbered circles, by *Bsa*I-mediated Type IIS assembly (sequential restriction enzyme digestion and ligation reactions). L1 TUs are assembled to Level 2 (L2) multi-TUs into one the four pCschlo vectors by *Sap*I-mediated Type IIS assembly. The recursive nature of Loop assembly means that this workflow can be repeated for higher level assemblies (L3, L4 etc). *Bsa*I and *Sap*I recognition site represented with red and blue triangles respectively. HS: homologous sequences, bended arrows: promoters, arrows: coding sequences and “T”: terminators. Blue filled rectangle: LacZ bacteria selection cassette. **b)** Microboxes are used to produce spores. 7 day old sporelings are bombarded with DNAdel™ nanoparticles coated with the desired DNA construct. After bombardment, sporelings are plated on selective media, and after 4 weeks successful transformants start to be visible. After a second round of selection (4 weeks), gemmae are produced and can be tested for homoplasmy by genotyping PCR. **c)** Top: Schematic representation of different constructs. Boxes represent the 5’ UTR region used. Numbers above the boxes correspond to the nucleotide position in relation to the CDS first nucleotide. Red “A” indicates the extra adenine nucleotide introduced by the common syntax. Cloned the region between Mp-*psbH* and Mp-*petB* (Mp-*psbH-petB*, 104 bp in length), 58 bp upstream of the Mp-*petB*, 48 bp upstream of *psbH*, 68 bp upstream of *MprbcL* and 123 bp upstream of Mp-*atpH* and. The amplified sequences were then fused with the Nt-*psbA* promoter (61 bp) and the mTurq2cp fluorescent protein coding sequence. The promoter and 5’UTR region (185 bp) of Mp-*rbcL* was also fused to mTurq2cp. All constructs were generated using the OpenPlant kit and Loop assembly. Bottom: Schematic representation of a L2 Loop construct to express the chloroplast codon optimized mTurq2cp fluorescent protein (Boehm et al., 2016) under the control of the tobacco Nt-*psbA* promoter and different combinations of PPR binding sequences (top figure) using the left and right homologous sequences for integration in the chloroplast *rbcL-trnR* intergenic region (Sauret-Güeto et al. 2020).

We cloned the intergenic region between the *Marchantia psb*H and *pet*B *genes* (104 bp in length) and sequences corresponding to the 5’ UTRs of the *pet*B gene (58 bp), *rbc*L (68 bp), *atp*H (123 bp) and *psb*H (48 bp). The amplified sequences were then fused downstream of the tobacco (*Nicotiana tabacum*) *psb*A promoter (61 bp). The intact Nt *psbA* promoter has been reported to have activity in *Marchantia*, albeit with low expression levels (Boehm et al. 2016). The hybrid promoter elements were assembled with a chloroplast codon optimised turquoise fluorescent protein reporter (mTurq2cp) (Boehm, et al., 2016).

Chloroplast protein synthesis is mediated by bacterial-type 70S ribosomes, and translation initiation is mediated by ribosome-binding sites, adjacent to the start codon on an mRNA. The sequence and spacing between the ribosome-binding sequence and the start codon is known to be important for the efficiency of translation initiation in cyanobacteria and chloroplasts **XXX** (Weiner et al. 2020). The default common syntax for Type IIS assembly DNA parts (Patron et al. 2015) introduces extra sequences at the termini of each element. The assembly of a 5’ UTR part can introduce an extra adenosine (A) nucleotide upstream of the ATG start codon. To test whether this has an effect on the expression efficiency of the transgene in *Marchantia* chloroplasts we generated two versions of the constructs, a version for standard assembly with an extra “A” and customised versions without. For the latter, we generated new L0 parts with ATGg as the 3’ overhang and mTurq2cp L0 constructs with ATGg as the 5’ overhang. We also generated constructs with mutant PPR binding sites, which contained sequence changes in the putative PPR protein binding site (Fig. 4c-d and Supplementary Table 6). As an additional control, we used a construct with the *Nt-psbA* core promoter fused to a 54 bp sequence containing the multi-cloning site from the pUC18 vector (45 bp) and a synthetic ribosome binding sequence (Hayashi et al. 2003) (hereafter called “control 5’UTR”). Transplastomic plants containing this construct showed very low levels of fluorescence.

The modified genes were assembled in chloroplast transformation vectors that contained the aadA spectinomycin resistance gene and flanking sequences for insertion by homologous recombination into the *rbc*L-*trn*R intergenic region of the *Marchantia* plastid genome. Chloroplasts were transformed by particle bombardment of germinating *Marchantia* spores, which are relatively easy to harvest in large numbers after sexual crossing, and stored indefinitely in a cold, desiccated state before use. DNAdel™ (Seashell Technology) nanoparticles were used as plasmid DNA carriers for the biolistic delivery into chloroplasts. The use of DNAdel™ reduces the time and labour required for loading of the plasmid DNA onto the microcarrier used for DNA delivery, compared to conventional metal carriers.

Three weeks after bombardment successful transformants were visible under a fluorescence stereomicroscope. After 6–8 weeks on antibiotic selection, plants were tested for homoplasticity (Supplementary Fig. 5). Five independent homoplastic lines for each construct were obtained. Little variation in levels of fluorescence was seen between the independent homoplastic lines, when examined using a stereo fluorescent microscope. Plants transformed with the 5’UTR *Mp-psbH* exhibited similar levels of expression to the control 5’UTR and were not further characterised (Supplementary Fig. 6).

### Testing the leader sequences

Three independent homoplastic lines for each construct were selected for further investigation. We developed and applied a three-step image processing pipeline to quantify chloroplast fluorescence intensity. This consisted of (i) acquisition of two-channel fluorescent micrographs using a confocal microscope, with a blue channel tuned to capture cyan fluorescent protein (CFP) fluorescence and a red channel tuned for chlorophyll autofluorescence, (ii) automated segmentation using a custom Fiji macro to identify regions of interest (ROI), and (iii) quantification of fluorescence intensity levels in each channel within each ROI. Mean CFP fluorescence intensity within each ROI was normalized by chlorophyll autofluorescence to account for fluorescence signal attenuation for plastids deeper within the sample (Markel 2018) (Supplementary Fig. 6 and Fig. 7). First we report the results from transformants containing custom 5’ UTR parts with native sequence and spacing adjacent to the start codon of the reporter gene. The highest levels of mTurq2cp fluorescence were measured in plants transformed with constructs containing the 5’UTR-*Mp-rbcL* leader sequence (Fig. 5a and Supplementary Table 7). Plants transformed with constructs containing mutations in the putative MRL1 PPR binding site within the 5’UTR-*MprbcL* sequence showed a reduction, but not complete loss of fluorescent protein levels. This is not unexpected since the ribosome binding sequence and promoter were still present.

**Fig. 5:**
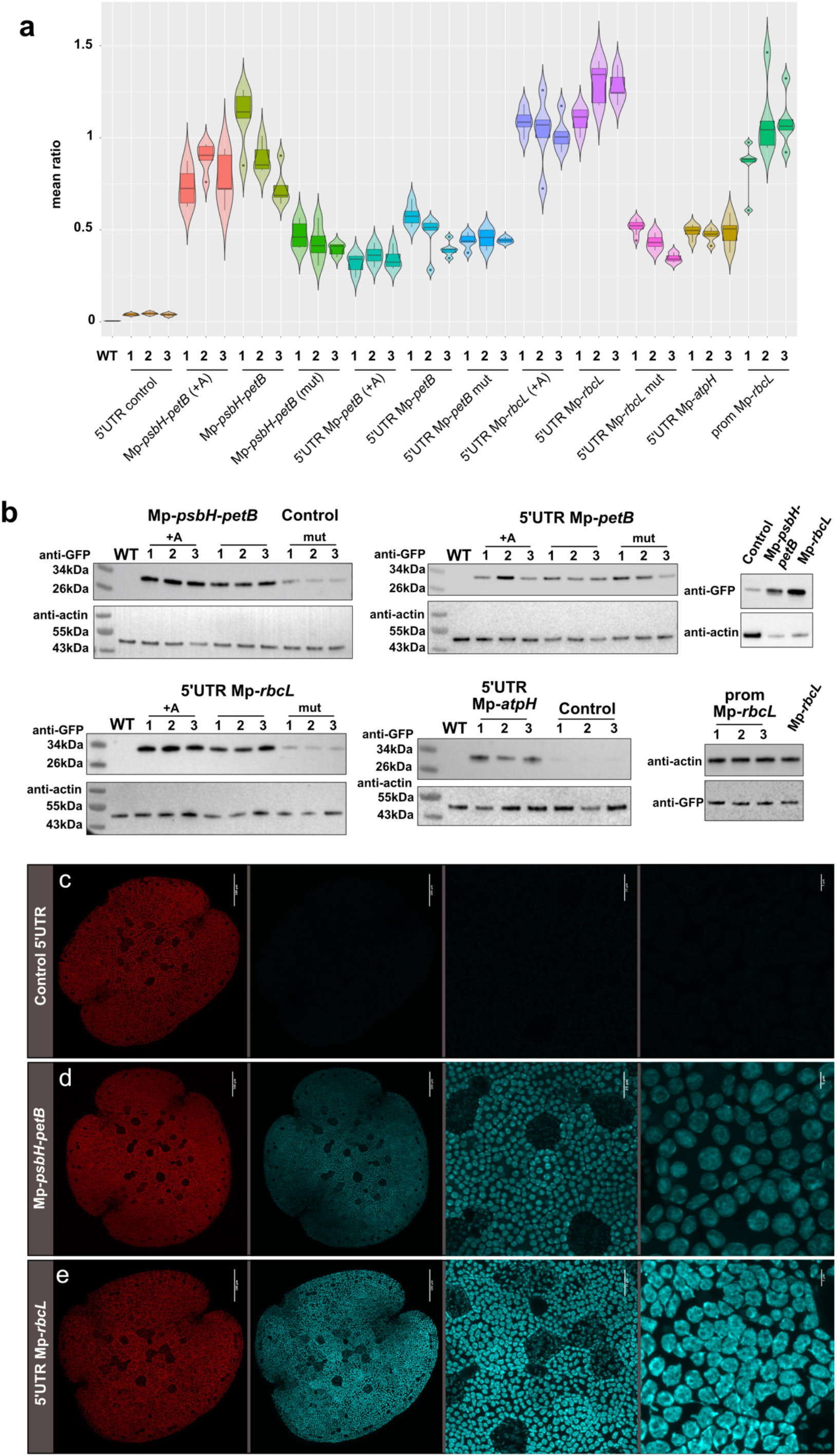
Foreign protein accumulation in transplastomic lines harbouring various candidate stabilisation elements. **a) Mean ratio of cyan and chlorophyll fluorescence** Five gemmae per line, for three lines per construct, were imaged and the ratio of cyan to chlorophyll fluorescence was calculated. 5’UTR Mp-*rbcL* confers the highest levels of expression followed by Mp-*psbH-petB*. 5’UTR-Mp*petB* and 5’UTR Mp-*atpH* have similar levels of expression. Expression levels are reduced for both when the predicted PPR binding sequence is mutated. The addition of an adenine between the 5’UTR and the mTurq2cp coding sequence does not affect the expression of mTurq2cp. **b) Western blots** Immunoblot analysis of mTurq2 accumulation in transplastomic lines. Total cellular proteins were separated by denaturing gel electrophoresis, blotted and probed with anti-GFP and anti-actin antibodies. Control 5’UTR, Mp-*psbH-petB*, 5’UTR Mp-*petB*, 5’UTR Mp-*rbcL* and 5’UTR Mp-*atpH*. +A: Adenine introduced by the common syntax present between the 5’UTR and the mTurq2cp coding sequence, Mut: predicted PPR binding sequence mutated. Numbers correspond to three independent lines per construct used. **c-e)** Microscopy images of Marchantia transplastomic 0 day gemmae expressing the mTurq2cp fluorescent protein under the control of the Nt-*psbA* promoter fused to different candidate stabilisation sequences: control 5’UTR, Mp-*psbH*-*petB*, and 5’UTR Mp*rbcL*. From left to right; first panel: Chlorophyll autofluorescence channel (Scale bars: 100 μm), second Panel: mTurq2cp channel (Scale bars: 100 μm), third and fourth panel: Higher magnification images showing mTurq2cp accumulation inside the chloroplasts of all cells (Scale bars: 20 μm and 5 μm, respectively). All images acquired using identical instrument settings. 5’UTR Mp*rbcL* confers the highest levels of expression followed by *Mp-psbH*-Mp-*petB*.

The Mp-*psbH-petB* intergenic region also conferred high levels of fluorescence, although levels were lower than those of plants containing the 5’UTR Mp-*rbcL* sequence. Plants transformed with constructs containing a 10bp mutation in the putative HCF152 PPR binding site (Supplementary Table 6) in the *Mp-psbH-petB* sequence showed reduced fluorescence levels. The 5’UTR Mp-*petB* sequence also conferred higher levels of fluorescence protein expression compared to the control 5’UTR but lower than that of the *Mp-psbH-petB* intergenic region. Plants transformed with constructs containing the 5’UTR *Mp-petB* sequence with a 15 bp mutation that removed the putative binding site for HCF152 did not show significant reduction in fluorescence (Supplementary Table 6). The 5’UTR *Mp-atpH* sequence produced levels of mTurq2cp fluorescence similar to that of 5’UTR *Mp-petB*.

The standardized syntax for Type IIS assembly of plant genes contains a site for gene fusions at the ATG initiation codon, which requires the sequence AATG to be placed at the junction of 5’UTR and coding sequence. We also tested the activity of constructs assembled this way, bearing an additional A residue adjacent to the start codon, in order to determine any effects on the efficiency of gene expression. Fluorescence levels were only slightly lowered compared to plants transformed with constructs containing the 5’UTR-(ATGg) sequences, indicating that the extra “A” introduced by the common syntax overhang did not have major effects on expression of the marker transgene. These observations were further supported by western blot studies of fluorescent protein levels in the plants.

Detergent soluble proteins were extracted from three independent lines for each construct, and fractionated by SDS polyacrylamide gel electrophoresis. mTurq2cp protein levels were assayed by western blotting using an anti-GFP antibody, and an anti-actin antibody was used to measure levels of endogenous actin protein as a loading control (Fig. 5b). Consistent with the results obtained using ratiometric imaging, the 5’UTR Mp-*rbcL* leader sequence conferred the highest levels of protein accumulation followed by the Mp-*psbH-petB* intergenic region. Constructs containing 5’UTR Mp-*petB* and 5’UTR *Mp-atpH* similar, lower levels of expression. The addition of an extra “A” between the 5’UTR and the mTurq2cp coding sequence did not greatly affect the expression of mTurq2cp in these experiments. However, substantially lower levels of fluorescent protein were seen in plants bearing mutations in the predicted PPR binding sites in 5’UTRs derived from Mp-*rbcL* and the Mp-*psbH-petB* intergenic region.

Based on these analyses the mRNA leader sequences corresponding to the *Mp-psbH-petB* intergenic region and 5’UTR Mp-*rbcL* were selected as the best candidates for generating high level gene expression in *Marchantia* chloroplasts. (Fig. 5c-e).

### *Marchantia rbcL* native promoter

The selected mRNA leader sequences with PPR-binding sites were all tested with the *N. tabacum psb*A promoter. To test the importance of the promoter in driving transgene expression, we cloned the entire promoter and 5’UTR region from the Mp-*rbcL* gene (185 bp upstream of the start codon), in order to compare it with the Nt-*psbA* promoter-driven version. Native transcripts from the Mp-*rbcL* promoter were found to accumulate at notably high levels in *Marchantia* (Fig. 2c). The native promoter was fused to the mTurq2cp coding sequence, and transformed into the *Marchantia* chloroplast genome as described for the other gene fusions. Confocal microscopy of the transformed plants confirmed (i) the exclusive chloroplast localisation of the expressed transgene, and (ii) high levels of fluorescent protein expression. High levels of mTurq2cp protein accumulation were further confirmed by ratiometric fluorescence measurements and a western blot analysis. However, the levels were not significantly over those conferred by the *Nt-psbA* promoter fusion (Fig.5a and Supplementary Fig. 6). This indicated that either both promoters had similar properties in *Marchantia* chloroplasts, or that rates of RNA transcription, mRNA stability, translation or protein stability might be saturated, and rate limiting.

### Quantification of transgene expression

In order to estimate the amount of protein produced in transplastomic *Marchantia* plants we expressed His6-tagged mTurquoise2 in *E. coli* under the control of the T7 promoter and purified the protein by affinity chromatography. (Supplementary Fig. 8). Serial dilutions of the purified mTurquoise2 were used to create a standard curve (fluorescence emission versus protein concentration) to allow accurate measurement of protein levels. Total protein was extracted from the *Marchantia* thallus tissue of plants harbouring different constructs (see Materials). The CFP fluorescence for each sample was then measured using a Clariostar plate reader and the protein concentration was calculated based on the standard curve. Up to 460 μg per g of tissue (~15% total soluble protein) was obtained from homoplastic plants harbouring the construct containing the *Nt-psbA* promoter and *Mp-rbcL* leader sequence (Fig.6a-b).

### Growth rates of transplastomic plants

Growth defects have been observed in plant species with high levels of chloroplast transgene expression (Hennig et al. 2007; Lössl et al. 2005). Very high levels of expression of a stable protein can lead to delayed plant growth (Oey et al. 2009). To test whether the accumulation of foreign proteins had an effect on *Marchantia* growth, we compared the growth of wild-type gemmae with those of lines transformed with the different constructs (Fig. 6c-k). The accumulation of fresh and dry weights were measured after one month of growth on agar-based media. Plants transformed with the construct containing the 5’UTR Mp-r*bcL, w*hich resulted in the highest levels of mTurq2cp accumulation, showed an approximately 35% biomass reduction compared to wild type. Interestingly, in comparison to other systems, *Marchantia* showed a higher tolerance to foreign protein accumulation in the chloroplast. For example, the potato showed significant biomass decrease in response to green fluorescence protein (GFP) overexpression. (Q. Yu, Barkan, and Maliga 2019). Interestingly, plants transformed with the construct containing the 5’UTR Mp-r*bcL c*onstruct showed lower size reduction than those transformed with the Mp-p*sbH* - Mp-p*etB* containing construct, despite higher levels of transgene accumulation.

**Fig. 6:**
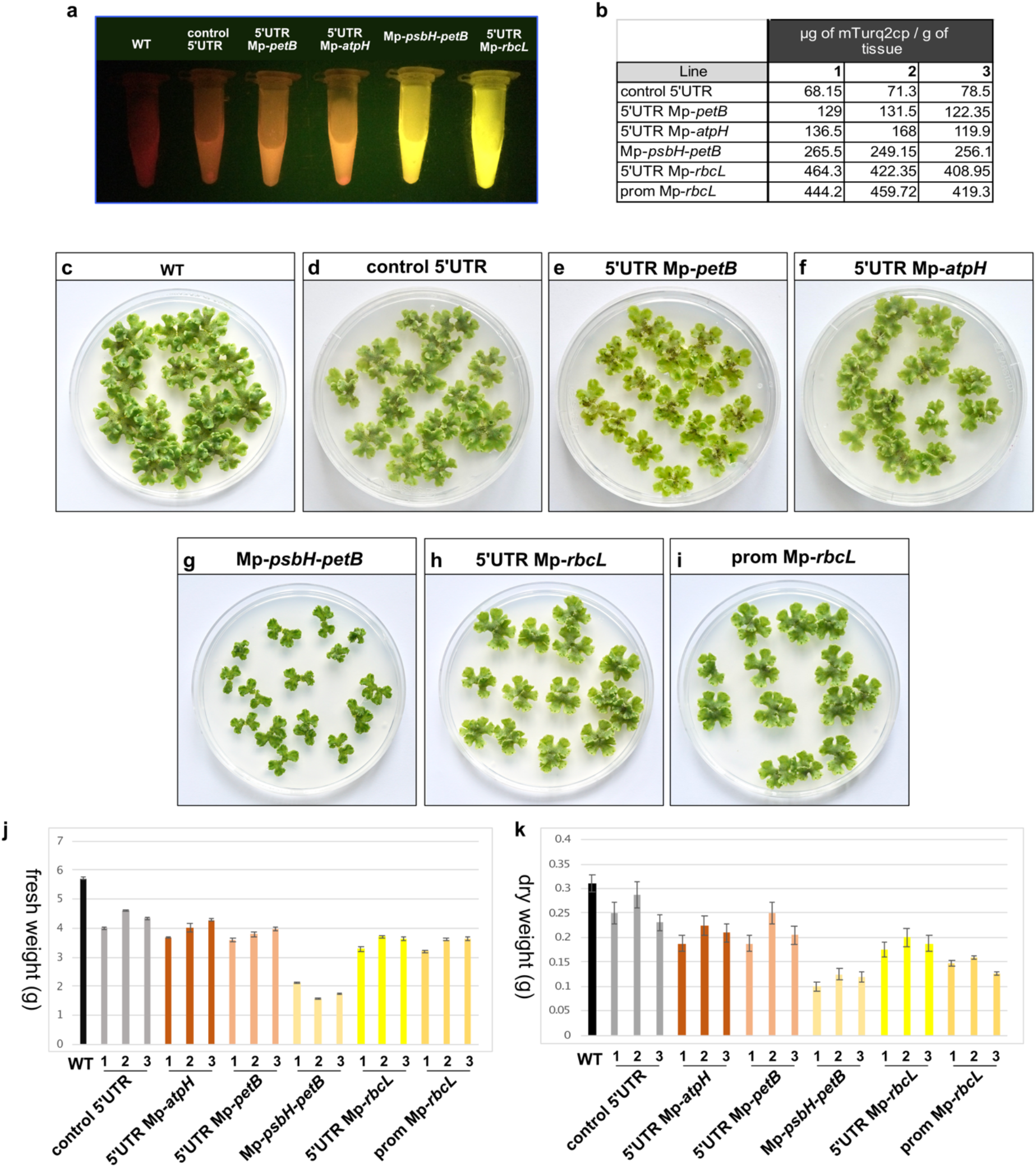
**a)** Total protein extract from 200mg of 1 month old gemmae under blue light transillumination. Red corresponds to chlorophyll autofluorescence and yellow to mTurq2cp fluorescence. Extract from plants transformed with the construct containing 5’UTR Mp*rbcL* are exhibiting the brightest fluorescence. **b**) Estimation of μg of mTurq2cp/g of fresh tissue, for three independent lines per construct. **c-i**) Comparison of growth between wild type and transplastomic *Marchantia* one month old gemmae expressing different constructs. All transplastomic plants showed a reduction in growth and biomass. Plants transformed with the construct containing the Mp-*psbH* - 5’UTR Mp-*petB* sequence showed the most extreme growth reduction phenotype. **j-k**) The fresh and dry weight was measured for 30 one month old gemmae, and the average values of two different experiments are shown on graphs I and J. Plants transformed with the construct containing the 5’UTR Mp-*rbcL*, even though they express the highest levels of mTurq2cp, they only showed an approximately 35% reduction in biomass. Error bars: standard error.

## DISCUSSION

Chloroplasts are attractive vehicles for transgene hyper-expression. Chloroplasts are sites for energy generation and high level protein expression and play a major role in metabolite production in plant cells. Plastid genes are present in high gene copy numbers per cell, can be highly transcribed, and are not subject to gene silencing. The plastid genome is compact and conserved across the terrestrial plants, and shows great promise as a platform for low-cost, large scale bioproduction.

*Marchantia* is one of the few plant species that have well-established methods for chloroplast transformation. It provides a facile testbed for chloroplast engineering, with simple culture requirements for culture (i.e. no need for glasshouses and expensive or specialised infrastructure for plant growth), and offers the benefits of spontaneous regeneration at high efficiency in the absence of phytohormones, fast selection for homoplasmy, simple microscopic observation and rapid amplification and propagation through both asexual and sexual life cycles. Further, a standardised system for rapid, semi-automated DNA assembly and libraries of modular DNA parts are available for *Marchantia*.

Recent work in the field has demonstrated the requirement for proper post-transcriptional regulation for high level gene expression in the chloroplasts of angiosperm plants (Prikryl et al. 2011; Legen et al. 2018; Pfalz et al. 2009; Zhelyazkova, Hammani, et al. 2012). In particular, nuclear-encoded PPR-proteins play a direct role in stabilizing the termini of specific mRNAs by direct binding, likely to protect the mRNAs against exoribonuclease degradation. The plastid genomes of early divergent plants, like *Marchantia*, possess a coding capacity broadly similar to gymnosperm and angiosperm species. However, gene regulation is different in a number of respects, such as the absence of RNA editing in *Marchantia*. Further, non-coding sequences in the plastid genome have diverged markedly. These key determinants of expression levels remain inaccessible to genetic manipulation, owing to insufficient understanding of native regulation and very limited availability of characterised parts. In order to fully exploit the potential benefits of the *Marchantia* system, we needed to “domesticate” important regulatory functions that allow properly regulated and high-level gene expression.

In this work, we describe the mapping of transcription patterns on the plastid genome of light-grown *Marchantia*. This allowed us to obtain empirical evidence for levels of transcription across the plastid genome. We precisely identified the promoter start sites for a number of highly expressed chloroplast genes. These genes have homologues in better-studied model systems, like tobacco, maize and *Arabidopsis*, where terminal sites for PPR protein binding to mRNAs have been characterized recently. However, sequence drift and the limited size of these functionally important sequences make them difficult to identify by inspection in widely divergent species.

While thousands of gymnosperm and angiosperm plastid genomes are available to build phylogenetic comparisons, the record for bryophytes has been sparse. We have engaged in a program of plastid genome sequencing to expand the data available for liverworts, hornworts and mosses. We have contributed 30 new bryophyte plastid genome sequences, and here, have used the newly expanded set to draw phylogenetic comparisons across the 5’ non-coding sequences of high abundance *Marchantia* transcripts. These regions correspond to mRNA termini that contain PPR protein binding sites in well-characterized angiosperm model systems. These fine-detail comparisons revealed conserved nucleotide sequences that may correspond to binding sites in *Marchantia*, and reflect an ancient origin for PPR-mediated control of gene expression in chloroplasts.

The identification of these conserved domains, which are putative PPR protein binding elements in the 5’ UTRs of chloroplast mRNAs, has allowed us to assemble a modular library of DNA parts that could confer transcript stability. In order to test the function of these novel 5’ UTR elements, the candidate sequences were each assembled as components of gene fusions between a chosen promoter and the mTurq2cp fluorescent protein coding sequence and terminator. The novel genes were incorporated into chloroplast transformation vectors, and homoplasmic transformants were generated. The levels of fluorescent protein expression in transformed plants were measured by microscopy-based ratiometric imaging, western blot analysis and protein extraction and quantitation. The presence of putative PPR protein binding sites at the 5’ termini of artificial mRNAs conferred markedly higher levels of reporter gene expression. Mutations within the putative binding domains reduced levels of gene expression. Highest levels of gene expression were seen in plants with reporter genes containing active promoters and the 5’ UTRs of *Mp-rbcL* and the *Mp-psbH-petB* intergenic region. A single inserted gene of interest could produce up to 15% of total soluble protein. Analysis of the growth rates of these plants showed that there was some penalty for hyperexpression in the form of slower growth. Lowered growth rates did not correspond directly to the level of ectopically expressed fluorescent protein, and it is possible that the mRNA transcripts themselves may interfere with growth, perhaps through competition with native transcripts for the different target PPR proteins. This indicates that conditional expression may be useful, through regulation of transcription in the chloroplast, regulation of mRNA stability through conditional expression of heterologous PPR proteins or supplementary expression of any limiting PPR proteins.

The identification and domestication of these mRNA stabilizing elements allows the prospect of enhanced gene design for engineering of the *Marchantia* plastid genome, to take advantage of the speed of this experimental system. Both the hybrid *Nt-psbA* promoter-5’ UTR *Mp-rbcL* and native *Mp-rbcL* promoter-5’ UTR sequences show high activity with minimal deleterious effects on growth, and look promising for future work in *Marchantia*. The transformation, regeneration and rescue of homoplasmic transformants in tobacco may take 6-9 months, while a similar experiment can take 8 weeks in *Marchantia*. Further, the vegetative life cycle for *Marchantia* can take a little as 2 weeks, and a single cycle through the sexual phase will give rise to millions of progeny as plant spores. *Marchantia* is a weed and can grow quickly. It may be useful as a cheap, easy to maintain and high yielding platform for small-scale bioproduction. Further, the DNA toolkit developed and characterized in *Marchantia* may function in plastids from a wide variety of plants.

## MATERIALS & METHODS

### Chloroplast isolation

Chloroplast isolation buffer (CIB) composition: 50 mM HEPES-KOH pH7.5, 0.33 M sorbitol, 1 mM MgCl2, 1 mM MnCl_2_, 2 mM EDTA. 5 mM Na-ascorbate and 1% (w/v) BSA (final concentration) were added immediately before use. Percoll^®^ (#17-0891-02, GE Healthcare) gradients were prepared as follows: 20mL 30% (v/v) Percoll solution was prepared by mixing 6ml Percoll and 14mL CIB. 10mL 70% (v/v) Percoll solution was prepared by mixing 7mLPercoll and 3mL CIB. For the preparation of 30%:70% (v/v) Percoll gradient, 15mL of 30% (v/v) Percoll were placed into a 50mL Falcon tube and 6mL of 70% (v/v) Percoll solution was carefully underlaid using a 5mL Gilson pipette.

Plants were grown in a 12h light : 12h dark cycle, and thallus tissue was harvested 2-3 hours after the start of the light cycle to minimise the amount of starch accumulated in chloroplasts. 40g of tissue was split into four equal parts and each homogenised using a mortar and pestle in 100mL of CIB. The homogenate was filtered through two layers of Miracloth (#475855, Millipore) into six 50mL Falcon tubes and centrifuged at 1200 g for 7 min. The supernatant was discarded and the pellet from each tube was carefully resuspended in 2mL of CIB using a paint brush. The resuspended pellet was transferred to the top of a Percoll gradient using a cut-off 1 mL pipette tip, and spun at 7000 g for 17 min at 4°C using slow acceleration and deceleration. Broken chloroplasts resided in the top fraction, while intact chloroplasts accumulated at the interface of the two Percoll layers. Chloroplasts from the interface were transferred to a 50mL falcon tube. 25mL of CIB was added, and tubes were centrifuged at 1500 g for 5 min at 4°C. The supernatant was discarded and the pellet was flash frozen in liquid N2.

### RNA extraction

RNA extraction was performed using the *mir*Vana™ miRNA Isolation Kit (#AM1560, ThermoFisher/Ambion) according to manufacturer instructions. After RNA extraction, samples were treated with DNase I using the TURBO DNA-fr*ee*™Kit (#AM1907, ThermoFisher/Ambion) following the manufacturer’s instruction. The integrity of the DNAse treated RNA was confirmed by capillary electrophoresis using the Agilent Bioanalyzer and the Agilent RNA 6000 Nano kit (#5067-1511, Agilent) according to the manufacturer’s instructions.

### Differential RNA-sequencing

Samples were treated and sequenced by vertis Biotechnologie AG, Germany. Detailed protocol in Supplementary Fig. 2.

### Differential RNA-sequencing processing

FASTQ read files were mapped against the Cam-1/2 plastid assembly (Genbank accession no. MH635409) using STAR-2.7.3a (Dobin et al. 2013). First, we generated a STAR index for the MH635409 assembly using the FASTA file of the assembly and existing genome annotation in GTF format (with settings: --runMode genomeGenerate --sjdbOverhang 74 -- genomeSAindexNbases 7). We then used multi-sample two pass mapping. In the first pass, samples were pooled and jointly mapped against the index to enable detection of unannotated transcripts and splice junctions. We supplied the genome annotation at this step and used conservative filtering of potential novel splice sites, (with settings: --alignIntronMax 800 -- outSJfilterCountUniqueMin 40 40 40 40 --outSJfilterCountTotalMin 50 50 50 50 -- sjdbOverhang 74). For the second pass we mapped each library against the index using both the existing genome annotation and the list of novel junctions generated by the first pass, using the same parameters as before. Mapping statistics for each library are provided in Supplementary Table 1.

We split the SAM output files into reverse and forward mapped reads using samtools view (H. Li et al. 2009) **XXX** and converted them to BAM format. Each file was sorted using samtools sort and per base coverage calculated using samtools depth. Base coverage was normalised and expressed as coverage per million mapped reads for each library. Coverage, data processing and visualisation was performed in R version 3.5.1. Plots were generated using ggplot2, ggbio (Yin, Cook, and Lawrence 2012) and circlize (Gu et al. 2014) packages.

Gene expression was quantified using kallisto (Bray et al. 2016). Protein coding transcript sequences were extracted from the MH635409 assembly sequence and used to build a kallisto index. FASTQ files from control libraries were processed using kallisto quant. Levels of gene expression were reported in units of transcripts per million (TPM).

### TSS identification

A 5’ end was annotated as a TSS when it had: i) a coverage in both TEX+/TEX-libraries of at least > 2 per million mapped reads, ii) a start at the same genomic position (nucleotide) in both libraries and iii) an enrichment > 1 in the TEX+ library (109 putative TSS in total). A 5’ end that was not enriched in TEX+ libraries was accepted as a TSS if it extended into an annotated gene (65 putative TSSs in total). We assigned 12 additional TSS that do not fall into the above categories when they extended into an annotated gene and a PEP promoter motif was predicted using MEME (Bailey et al. 2015).

### Marchantia PPR homolog prediction

Orthofinder was used (Emms and Kelly 2015) for the identification of PPR and HAT homologs between *M. polymorpha* and *A. thaliana* and *maize*.

### DNA extraction, sequencing and *de novo* bryophyte plastid genome assemblies

DNA extraction, sequencing and *de novo* assembly of plastid genomes were performed according to (Y. Yu et al. 2019). In addition, NGS data generated for a previous study (Y. Yu et al. 2019; Liu, Medina, and Goffinet 2014) were used for *de novo* assembly of *Anomodon attenuates, Atrichum angustatum, Bartramia pomiformis, Bryum argenteum, Entosthodon attenuates, Funaria hygrometrica, Hypnum imponens, Orthotrichum stellatum, Ptychomnion cygnisetum, Sphagnum palustre, Tetraphis pellucida* and *Ulota hutchinsiae* moss plastid genome sequences. Assemblies were performed using GetOrganelle (J.-J. Jin et al. 2020) (Supplementary Table 5). and annotated using GeSeq (Tillich et al. 2017). Genome alignments were performed using MUSCLE (Edgar 2004a).

### Marchantia chloroplast DNA manipulation

Genomic DNA was extracted according to (Sauret-Güeto et al. 2020). Constructs were generated using DNA parts and vectors from the OpenPlant kit (Sauret-Güeto et al. 2020). Construct sequences are listed in Supplementary Table 6. Primers used for construct generation are listed in Supplementary Table 8. Chloroplast transformation was performed as previously described in (Sauret-Güeto et al. 2020). The genotyping of transplastomic lines was performed as previously described in (Sauret-Güeto et al. 2020). Genotyping primers used are listed in Supplementary Table 8.

### Imaging

Gemmae were plated on half strength Gamborg B5 plus vitamins (#G0210, Duchefa Biochemie) with 1.2% (w/v) agar plates and placed in a growth cabinet for 3 days under continuous light with 150 μE m^−2^ s^−1^ light intensity at 21 °C. A gene frame (#AB0576, ThermoFisher) was positioned on a glass slide and 30 μL of half strength Gamborg B5 1.2% (w/v) agar placed within the gene frame. 5 gemmae were then placed within the media filled gene frame, 30 μL of milliQ water was added and then a cover slip was used to seal the geneframe. Plants were then imaged immediately using an SP8 fluorescent confocal microscope All images were acquired using the same instrument setting, Cyan and chlorophyll. 16 Z stacks, 3 μm thickness.

Images were acquired on an upright Leica SP8X confocal microscope equipped with a 460-670 nm supercontinuum white light laser, 2 CW laser lines 405nm, and 442 nm, and 5 channel spectral scanhead (4 hybrid detectors and 1 PMT). Imaging was conducted using either a 20x air objective (HC PL APO 20x/0.75 CS2) or a 40x water immersion objective (HC PL APO 40x/1.10 W CORR CS2). Excitation laser wavelength and captured emitted fluorescence wavelength window were as follows: for mTurq2cp (442 nm, 460-485 nm) and for chlorophyll autofluorescence (488 or 515, 670-700 nm). Chlorophyll autofluorescence was imaged simultaneously with mTurq2cp.

### Plastid segmentation pipeline

Plastid segmentation was achieved using an automated Fiji macro as described previously (Markel 2018), the source code is included in Supplementary Fig. 7c. In brief, the chlorophyll autofluorescence channel was duplicated, and the new copy subjected to a series of smoothing and thresholding steps using the Phansalkar algorithm (Phansalkar et al., n.d.), and the subsequent segmented regions were split using a watershed algorithm. Regions of interest were then used for quantification of marker gene and chlorophyll fluorescence and analysis of plastid parameters such as size and shape. Analysis in Figure 5 is based on the average fluorescence intensity within each ROI, with the CFP channel normalized by the chlorophyll channel. The full dataset (including additional parameters such as maximum and minimum fluorescence intensity within each ROI as well as area of ROIs) is included as Supplementary Table 7.

### Western blotting

*Marchantia* thallus tissue (100 mg) was excised from plants grown for 4 weeks on half strength Gamborg B5 medium including vitamins with 1.2% (w/v) agar, at 21 °C in continuous light, 150 μE m^−2^ s^−1^) and ground in liquid nitrogen. The tissue powder was resuspended in 500 μL 5× Laemmli loading buffer (0.2 M Tris-Hcl pH 6.8, 5 % w/v SDS, 25 % v/v glycerol, 0.25 M DTT, 0.05 % w/v bromophenol blue) with added Roche cOmplete protease inhibitor (# 11836170001, Roche. Samples were further diluted 21 times in 5x Laemmli loading buffer containing Roche protease inhibitor, heated at 95 °C for 5 minutes and centrifuged at 1000 g for 10 minutes. The supernatant was transferred to a new tube. Equal amounts of proteins were separated by denaturing electrophoresis in NuPAGE gel (#NP0322BOX, Invitrogen) and electro-transferred to nitrocellulose membranes using the iBlot2 Dry Blotting System (ThermoFisher). mTurq2cp was immunodetected with anti-GFP antibody (1:4000 dilution) (JL-8, #632380, Takara) and anti-mouse-HRP (1:15000 dilution) (#A9044, Sigma) antibodies. Actin was immunodetected with anti-actin (plant) (1:1500 dilution) (#A0480, Sigma) and (1:15000 dilution) anti-mouse-HRP (#A9044, Sigma) antibodies, using the iBind™ Western Starter Kit (#SLF1000S, ThermoFisher). Western blots were visualised using the ECL™ Select Western Blotting Detection Reagent (#GERPN2235, GE) following the manufacturer’s instructions. Images were acquired using a Syngene Gel Documentation system G:BOX F3.

### Plant biomass estimation

For each line 30 gemmae were placed on two petri dishes with 25mL of media (half strength Gamborg B5 plus vitamins) and grown for a month, at 21 °C, with continuous light, 150 μE m^−2^ s^−1^. The fresh and dry weight was measured using a scale.

### Total soluble protein estimation

Marchantia thallus tissue (200 mg) from 4 week old plants grown on half strength Gamborg B5 medium including vitamins and 1.2% (w/v) agar, at 21 °C in continuous light,150 μE m^−2^ s^−1^ was ground in liquid nitrogen and resuspended in 700 μL protein extraction buffer (50 mM Tris-HCl pH 7.5, 150 mM NaCl, TWEEN 20 0.1% (v/v), 10 % (v/v) glycerol, 1 mM DTT) plus Roche cOmplete protease inhibitor (# 11836170001, Roche). Total soluble protein concentration was estimated using a Pierce™ 660nm Protein Assay Kit as above (#22662, Thermo Scientific).

### Protein yield estimation

*E. coli* BL21 Star (DE3) (#C601003, Invitrogen,) was transformed with the pCRB SREI6His plasmid (Boehm et al., 2016) to express the mTurquoise2 protein. A culture of 10 mL was used to inoculate 250 mL of LB medium supplemented with ampicillin and grown in 2.5L baffled Tunair™ shake flasks (#Z710822, Sigma Aldrich) at 37°C with vigorous shaking (200 rpm). Cultures were monitored by spectrophotometry until OD600 reached 0.6. T7 RNA polymerase expression was induced by the addition of IPTG to a final concentration of 1 mM. Cultures were grown for 5 h at 30 °C, with shaking at 200 rpm. Cells were then harvested by centrifugation at 5000g for 12 min at 4 °C. To purify the recombinant protein under native conditions, the pellet was processed using the Ni-NTA Fast Start Kit (#30600, Qiagen), and cells were disrupted by lysozyme and detergent treatment according to the manufacturer’s instructions. Purified protein was concentrated using an Amicon Filter 3K (#UFC500324, Millipore). In order to avoid any interference with downstream procedures, imidazole was removed using a Zeba™ spin desalting column (#89882, Thermo Scientific) following the manufacturer’s protocol. Purified protein was stored in 50 mM sodium phosphate, pH 7.4 with 5 mM benzamidine at −20 °C.

The concentration of the mTurquoise2 protein was determined using a Pierce™ 660nm Protein Assay Kit (#22662, Thermo Scientific) and used as reference to build a mTurquoise2 standard curve (linear regression) based on fluorescence (random fluorescence units (RFU)) against concentration.

This curve was employed to estimate mTurq2cp protein amount in Marchantia samples (prepared following the same steps described in the total soluble protein estimation) per gram of tissue. Samples values were adjusted by subtracting the fluorescence values of the blank. In all the cases, a CLARIOstar (BMG) plate reader was used with an excitation and emission wavelength appropriate for mTurq2cp measurement (excitation:430-20 nm, emission: 474-20 nm, gain 500 nm).

## Supporting information

transcription data

transcription start sites

## Acknowledgements

We are thankful to Suvi Honkanen for advice throughout the project.

This work was funded as part of the BBSRC/EPSRC OpenPlant Synthetic Biology Research Centre Grant BB/L014130/1 to J.H., BBSRC BB/F011458/1 for confocal microscopy to J.H., BBSRC Research Studentships for M.R. (1943399) and National Natural Science Foundation of China (NSFC) (No. 31970227) to YY.

## Author contributions

JH and EF designed the project. EF and MR analysed the dRNAseq data, EF, JR and SWT performed cloning, EF and SSG carried out imaging, KM developed and performed imaging analysis, EF, AP and FGC performed protein western blotting and protein yield estimation, YY and HS sequenced bryophyte plastid genomes, YL and BG provided the NGS data for the moss plastid genome assemblies. JH and EF wrote the manuscript and all authors commented on the manuscript.

