## Supplementary information for "Comparative analysis of early divergent land plants and construction of DNA tools for hyper-expression in *Marchantia* chloroplasts"

### SUPPLEMETARY INFORMATION

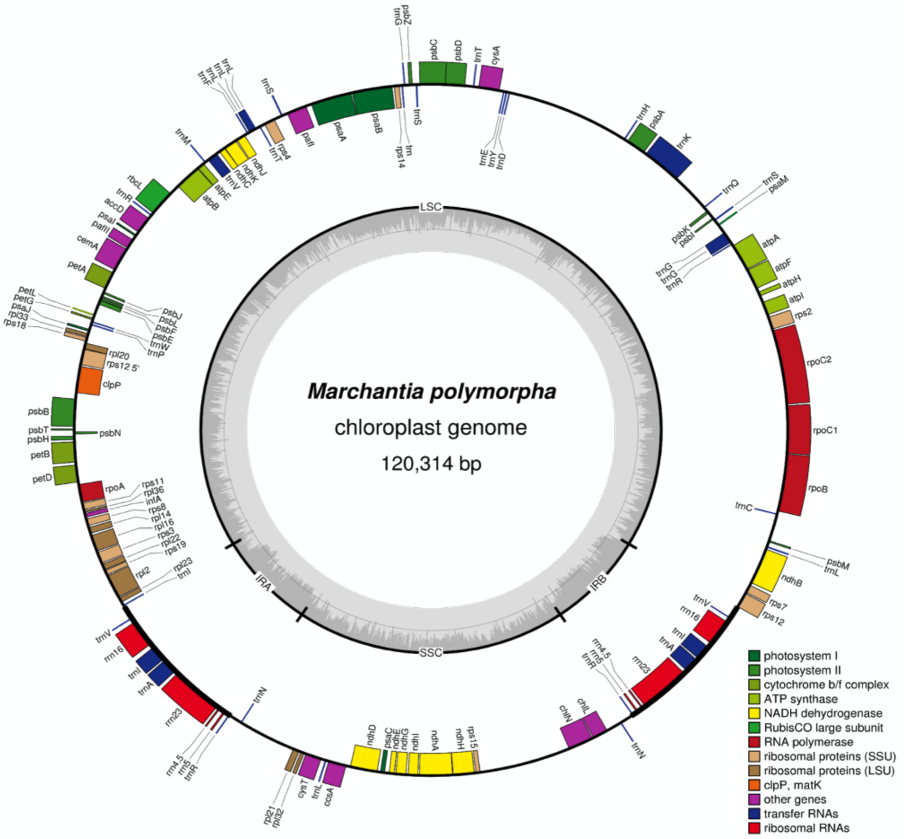

**Supplementary Fig. 1: Operon and gene map of the *Marchantia* plastid genome.**
The outer circle depicts the gene organization of the *Marchantia* plastid genome (MH635409). The graph was generated using OGDraw (Lohse et al. 2013). Genes are color coded based on their function listed at the bottom right of the figure. The Cam-1/2 plastome assembly was validated by comparison to both Sanger sequencing data covering ~10% of the plastome and the newly published Kit-2 plastome (NC_037507.1) assembly (Bowman et al. 2017). In both cases, validation supports a highly accurate assembly process.

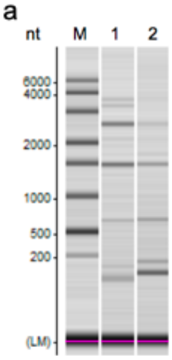

Analysis of the total RNA samples on a Shimadzu MultiNA microchip electrophoresis system. M: ladder

b

| No | Sample | Conc. (ng/µl) | Amount (µg) | Ratio 23S/16S |
|---|---|---|---|---|
| 1 | A | 123 | 29.6 | 0.9 |
| 2 | B | 44 | 7.7 | N/A |

Description of the RNA samples

c

| No. | Treatment | 5' Barcode | 3' Barcode | PCR cycles (FL) | PCR cycles (5') |
|---|---|---|---|---|---|
| 1+ | +TEX | ATAGAGAG | GAATTCGT | 12 | 6 |
| 2+ | +TEX | AGAGGATA | GAATTCGT | 12 | 6 |
| 1- | -TEX | CTCCTTAC | GAATTCGT | 12 | 6 |
| 2- | -TEX | TATGCAGT | GAATTCGT | 12 | 6 |

Description of the cDNA samples

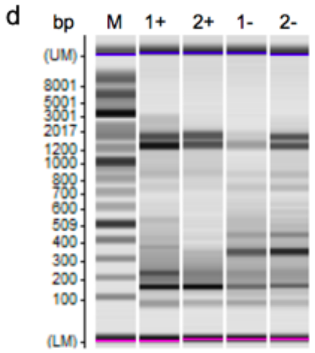

Analysis of the PCR-amplified full-length cDNAs on a Shimadzu MultiNA microchip electrophoresis system. M = ladder

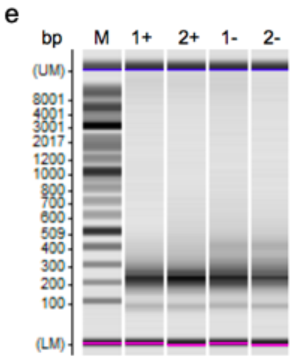

Analysis of the PCR-amplified 5' fragment cDNAs on a Shimadzu MultiNA microchip electrophoresis system. M = ladder

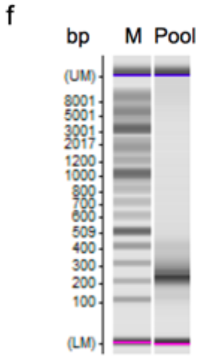

Analysis of the size fractionated cDNA libraries on a Shimadzu MultiNA microchip electrophoresis system. M = ladder

**Supplementary Fig. 2: Preparation of +/- TEX cDNA libraries for Illumina sequencing**

**a-b) Analysis of total RNA**

The total RNA samples were examined by capillary electrophoresis and RNA concentration was determined.

**c) cDNA synthesis from +/-TEX treated RNA**

The total RNA samples were split into two halves and one half was subjected to Terminator exonuclease (TEX) treatment. 2U of TEX enzyme (#TER51020, Lucigen) per 500 ng of RNA were used. Incubation was 1 hour at 30°C. The other half was left untreated (-TEX). The + and -TEX treated RNAs were poly(A)-tailed using poly(A) polymerase. The 5'PPP were converted to 5’P structures using RNA 5' Polyphosphatase (#RP8092H, Epicentre). 5’ Illumina sequencing adaptor was ligated to the 5'P of the +/-TEX treated RNA. First- strand cDNA synthesis was performed using an oligo(dT)-adapter primer and the M-MLV reverse transcriptase. The resulting cDNAs were PCR-amplified to about 10-20 ng/µl using a high-fidelity DNA polymerase (cycle numbers are indicated in the table).

**d)** The cDNAs were purified using the Agencourt AMPure XP kit (Beckman Coulter Genomics) and were analyzed by capillary electrophoresis

**e)** For Illumina sequencing, 100 – 300 bp long 5' fragments were isolated from the full-length cDNAs. For this purpose, the cDNA preparations were fragmented and the 5'-cDNA fragments were then bound to streptavidin magnetic beads. The bound cDNAs were blunted and the 3' Illumina sequencing adapter was ligated to the 3' ends of the cDNA fragments. The bead bound cDNAs were finally PCR-amplified. The PCR cycles performed and the barcode sequences, which are attached to the 5' and 3' ends of the cDNAs, are described in the table at (c).

**f) Pool generation and size fractionation**

For Illumina NextSeq sequencing, the samples were pooled in approximately equimolar amounts. The library pool was fractionated in the size range of 200-500 bp using a preparative agarose gel. An aliquot of the size fractionated cDNA pool was analyzed by capillary electrophoresis. The cDNAs have a size of about 200 – 500 bp. The primers used for PCR amplification were designed for TruSeq sequencing according to the instructions of Illumina. The following adapter sequences flank the DNA insert:

TruSeq_Sense_primer i5 Barcode 5'-AATGATACGGCGACCACCGAGATCTACAC-NNNNNNNN-ACACTCTTTCCCTACACGACGCTCTTCCGATCT-3’
TruSeq_Antisense_primer i7 Barcode 5’-CAAGCAGAAGACGGCATACGAGAT-NNNNNNNN-GTGACTGGAGTTCAGACGTGTGCTCTTCCGATCT-3’
The combined length of the flanking sequences is 136 bases.

The cDNA pool was single end sequenced on an Illumina NextSeq 500 system using 1x75 bp read length.

Sample A used for TSS predictions.
Both Sample A and B used for the identification of highly expressed genes.

**a**

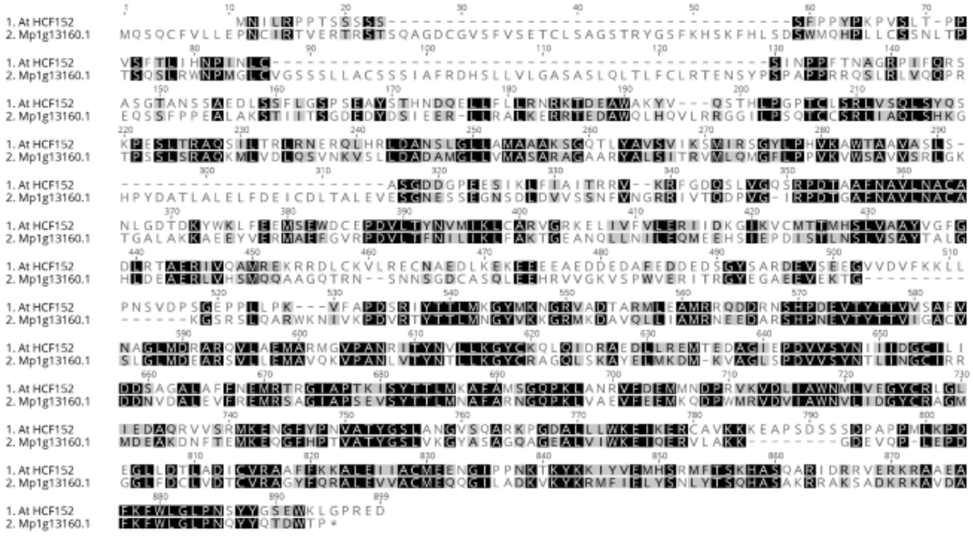

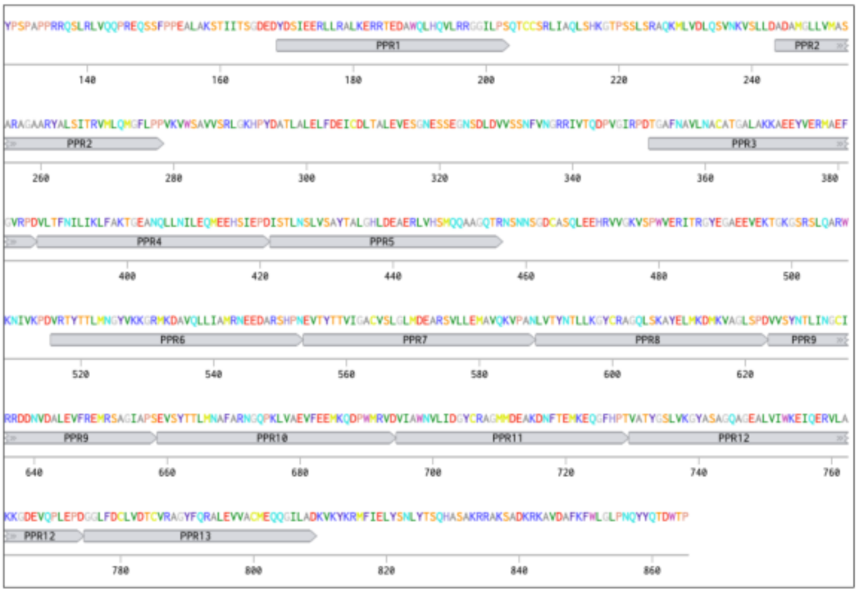

| P | P | P | P | P | P | P | P | P | P | P | P | P | Type |
|---|---|---|---|---|---|---|---|---|---|---|---|---|---|
| E | G | N | N | N | T | T | N | N | T | N | G | D | 5 |
| S | P | D | D | R | N | N | D | S | D | T | D | D | Last |
| . | . | U | U | C | A | A | U | C | G | C | U | . | Match 5 |
| | | 0.61 | 0.61 | 0.15 | 0.61 | 0.61 | 0.61 | 0.51 | 0.68 | 0.32 | 0.09 | | Prob |
| . | . | A | A | A,G,U | G | G | A | U | A,U | U | C | . | Match Last |
| | | -0.61 | -0.61 | -0.05 | -0.25 | -0.25 | -0.61 | -0.10 | -0.29 | 0.17 | 0.03 | | Prob |

**b**

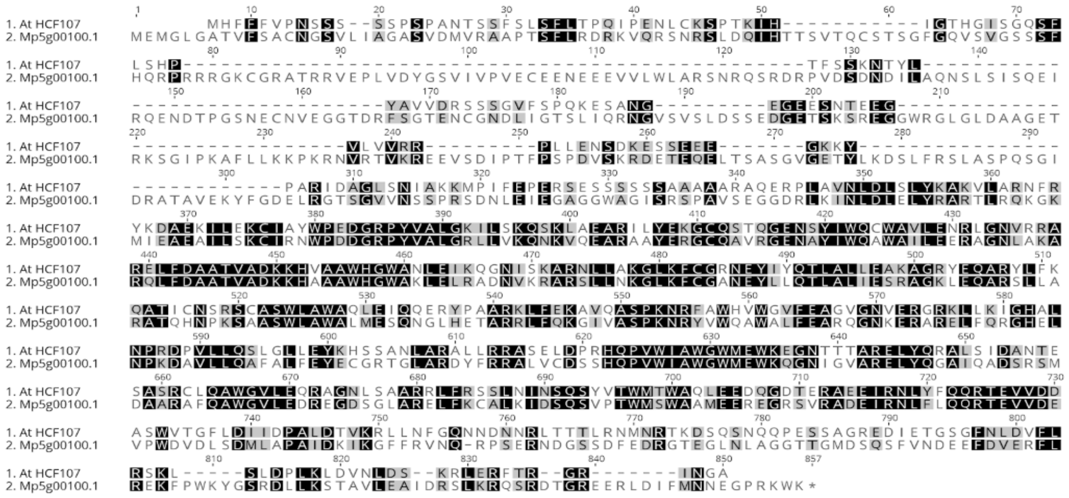

**c**

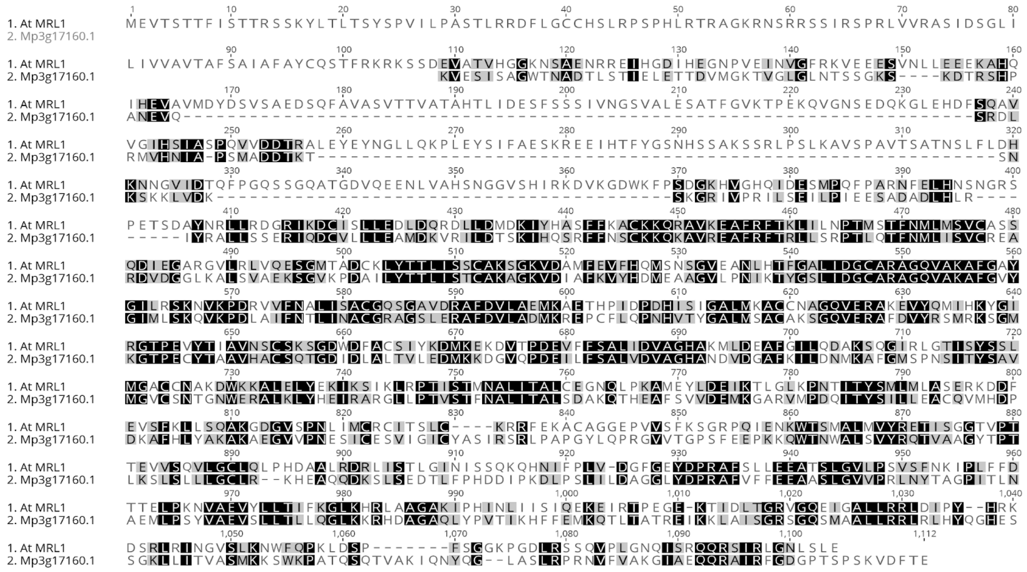

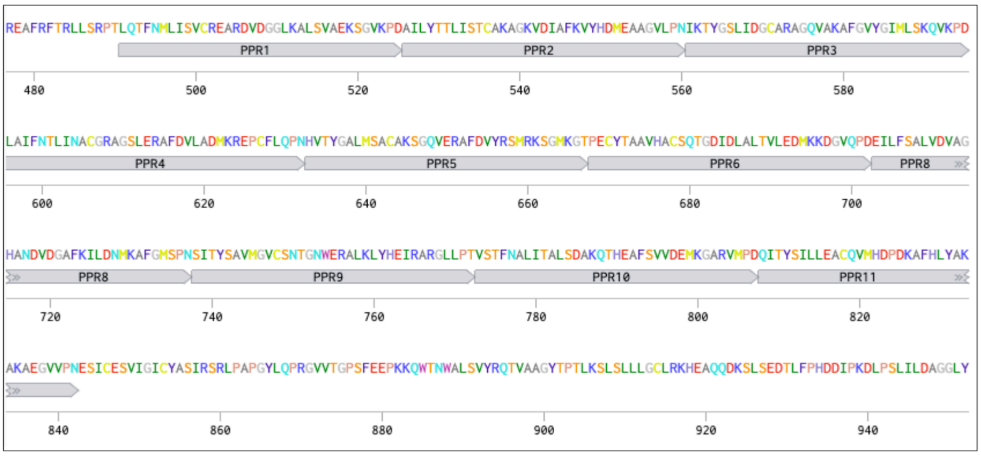

| P | P | P | P | P | P | P | P | P | P | Type |
|---|---|---|---|---|---|---|---|---|---|---|
| N | T | G | N | G | T | S | S | N | S | 5 |
| D | N | D | N | T | D | N | T | D | N | Last |
| U | A | U | C | . | G | A | . | U | A | Match 5 |
| 0.61 | 0.61 | 0.09 | 0.63 | | 0.68 | 0.44 | | 0.61 | 0.44 | Prob |
| A | G | C | U | . | A,U | C | . | A | C | Match Last |
| -0.61 | -0.25 | 0.03 | 0.16 | | -0.29 | -0.19 | | -0.61 | -0.19 | Prob |

d

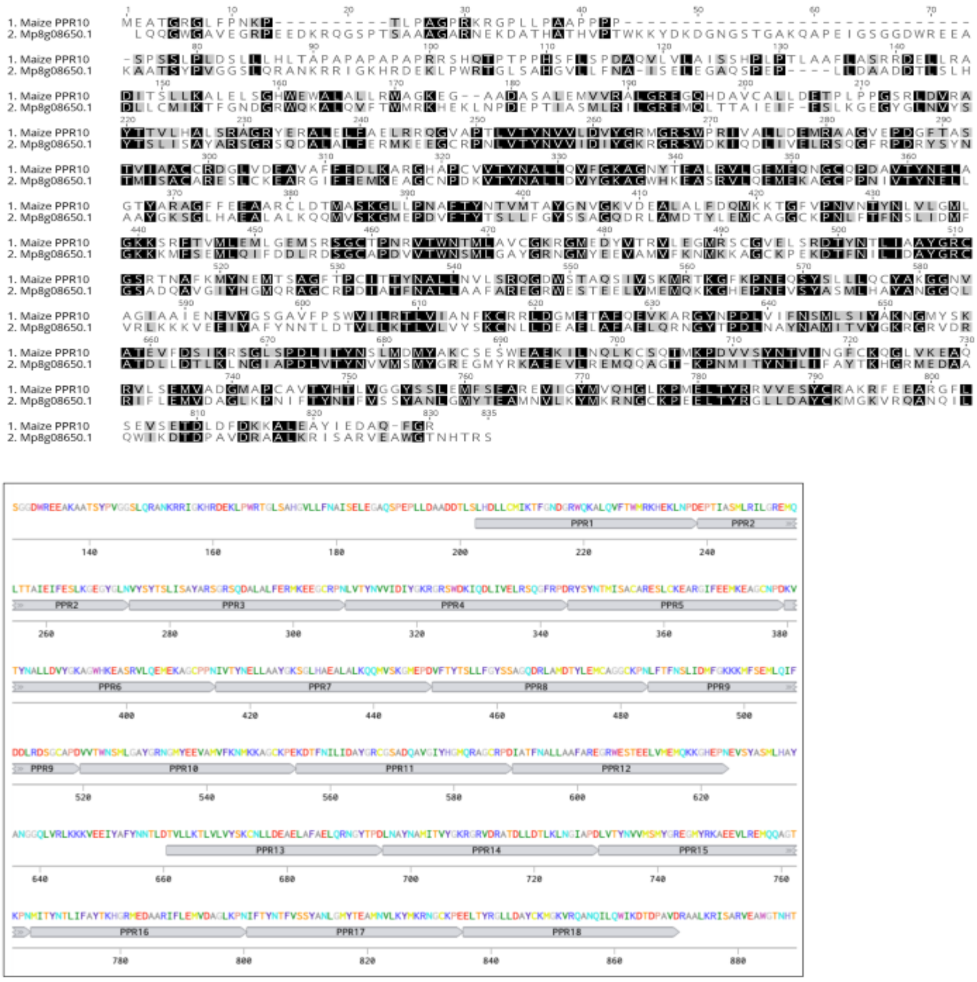

| | | | | | | | | | | | | | | | | | | |
|---|---|---|---|---|---|---|---|---|---|---|---|---|---|---|---|---|---|---|
| P | P | P | P | P | P | P | P | P | P | P | P | P | P | P | P | P | P | Type |
| L | A | T | N | N | N | N | T | N | N | N | N | K | N | N | N | N | R | 5 |
| D | N | N | D | D | N | D | N | D | E | D | N | D | D | N | N | E | D | Last |
| . | C | A | U | U | C | U | A | U | G | U | C | . | U | C | C | G | . | Match 5 |
| | 0.11 | 0.61 | 0.61 | 0.61 | 0.63 | 0.61 | 0.61 | 0.61 | 0.06 | 0.61 | 0.63 | | 0.61 | 0.63 | 0.63 | 0.06 | | Prob |
| . | A | G | A | A | U | A | G | A | U | A | U | . | A | U | U | U | . | Match Last |
| | 0.07 | -0.25 | -0.61 | -0.61 | 0.16 | -0.61 | -0.25 | -0.61 | 0.04 | -0.61 | 0.16 | | -0.61 | 0.16 | 0.16 | 0.04 | | Prob |

**Supplementary Fig. 3: Marchantia HCF152, HCF107, MRL1 and PPR10 homologs**

Marchantia PPR homolog predictions were made using Orthofinder (Emms and Kelly 2015)

a) Top: Amino acid sequence alignments of AtHCF152 (AT3G09650) and Mp*PPR_16* (Mp1g13160.1), using MUSCLE (Edgar 2004). Marchantia HCF152 homolog PPR domain (middle) and binding site (bottom) prediction based on Cheng et al. 2016.

b) Top: Amino acid sequence alignments of AtHCF107 (AT3G17040) and Mp5g00100.1, using MUSCLE.

c) Top: Amino acid sequence alignments of AtMRL1 (AT4G34830) and Mp*PPR_28* (Mp3g17160.1), using MUSCLE. Marchantia MRL1homolog PPR domain (middle) and binding site (bottom) prediction based on Cheng et al., 2016.

d) Top: Amino acid sequence alignments of ZmPPR10 (Genbank: AQK81740) and Mp*PPR_41* (Mp8g08650.1), using MUSCLE. Marchantia PPR10homolog PPR domain (middle) and binding site (bottom) prediction based on Cheng et al., 2016.

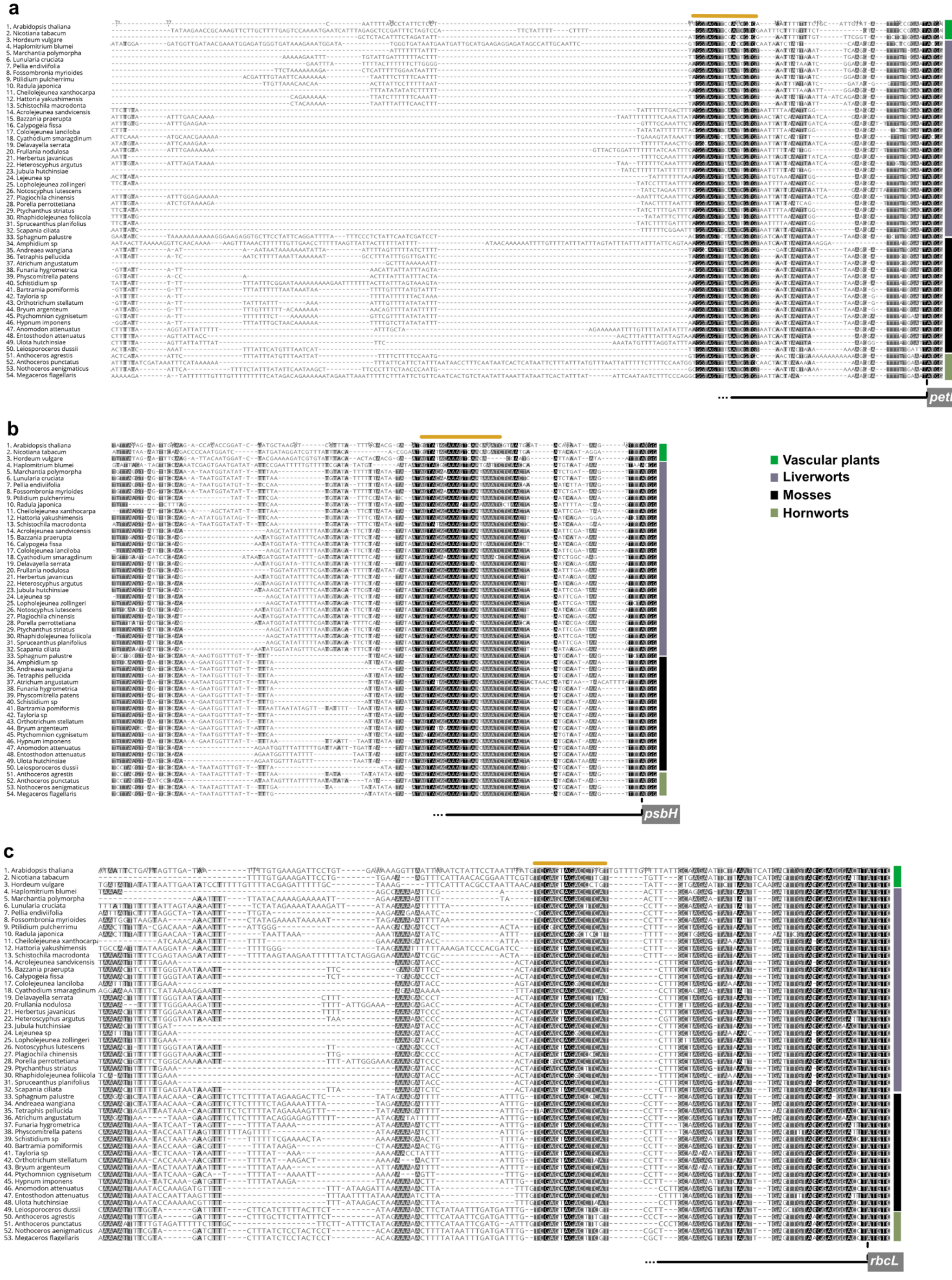
a
1. Arabidopsis thaliana
2. Nicotiana tabacum
3. Hordeum vulgare
4. Haplomitrium blumei
5. Marchantia polymorpha
6. Lunularia cruciata
7. Pellia endiviifolia
8. Fossombronia myrioides
9. Ptilidium pulcherrimu
10. Radula japonica
11. Cheilolejeunea xanthocarpa
12. Hattoria yakushimensis
13. Schistochila macrodonta
14. Acrolejeunea sandvicensis
15. Bazzania praerupta
16. Calypogeia fissa
17. Cololejeunea lanciloba
18. Cyathodium smaragdinum
19. Delavayella serrata
20. Frullania nodulosa
21. Herbertus javanicus
22. Heteroscyphus argutus
23. Jubula hutchinsiae
24. Lejeunea sp
25. Lopholejeunea zollingeri
26. Notoscyphus lutescens
27. Plagiochila chinensis
28. Porella perrottetiana
29. Ptychanthus striatus
30. Rhaphidolejeunea foliicola
31. Spruceanthus planifolius
32. Scapania ciliata
33. Sphagnum palustre
34. Amphidium sp
35. Andreaea wangiana
36. Tetraphis pellucida
37. Atrichum angustatum
38. Funaria hygrometrica
39. Physcomitrella patens
40. Schistidium sp
41. Bartramia pomiformis
42. Tayloria sp
43. Orthotrichum stellatum
44. Bryum argenteum
45. Ptychomnion cygnisetum
46. Hypnum imponens
47. Anomodon attenuatus
48. Entosthodon attenuatus
49. Ulota hutchinsiae
50. Leiosporoceros dussii
51. Anthoceros agrestis
52. Anthoceros punctatus
53. Nothoceros aenigmaticus
54. Megaceros flagellaris
petB
b
psbH
c
rbcL
Vascular plants
Liverworts
Mosses
Hornworts

d

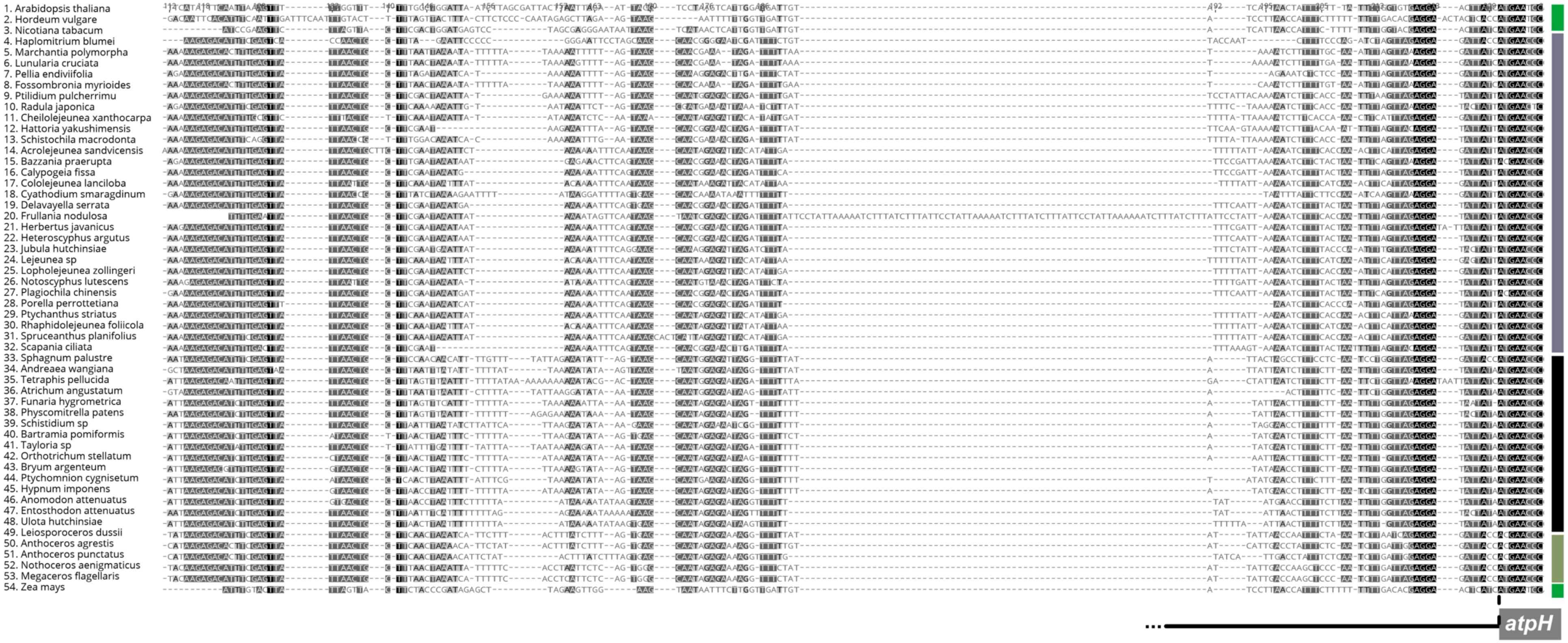
1. Arabidopsis thaliana
2. Hordeum vulgare
3. Nicotiana tabacum
4. Haplomitrium blumei
5. Marchantia polymorpha
6. Lunularia cruciata
7. Pellia endiviifolia
8. Fossombronia myrioides
9. Ptilidium pulcherrimu
10. Radula japonica
11. Cheilolejeunea xanthocarpa
12. Hattoria yakushimensis
13. Schistochila macrodonta
14. Acrolejeunea sandvicensis
15. Bazzania praerupta
16. Calypogeia fissa
17. Cololejeunea lanciloba
18. Cyathodium smaragdinum
19. Delavayella serrata
20. Frullania nodulosa
21. Herbertus javanicus
22. Heteroscyphus argutus
23. Jubula hutchinsiae
24. Lejeunea sp
25. Lopholejeunea zollingeri
26. Notoscyphus lutescens
27. Plagiochila chinensis
28. Porella perrottetiana
29. Ptychanthus striatus
30. Rhaphidolejeunea foliicola
31. Spruceanthus planifolius
32. Scapania ciliata
33. Sphagnum palustre
34. Andreaea wangiana
35. Tetraphis pellucida
36. Atrichum angustatum
37. Funaria hygrometrica
38. Physcomitrella patens
39. Schistidium sp
40. Bartramia pomiformis
41. Tayloria sp
42. Orthotrichum stellatum
43. Bryum argenteum
44. Ptychomnion cygnisetum
45. Hypnum imponens
46. Anomodon attenuatus
47. Entosthodon attenuatus
48. Ulota hutchinsiae
49. Leiosporoceros dussii
50. Anthoceros agrestis
51. Anthoceros punctatus
52. Nothoceros aenigmaticus
53. Megaceros flagellaris
54. Zea mays
atpH

**Supplementary Fig. 4**

Multiple sequence alignments of regions upstream the 5′ UTR of (a) *petB*, (b) *psbH*, (c) *rbcL and* (d) *atpH*, of the 51 bryophyte plastid genomes used in this study and key angiosperms, performed with MUSCLE (Edgar 2004). ATG site is indicated with a dashed line. Coding sequence is indicated with a grey box. The predicted PPR binding site is highlighted by an orange line above. The coloring used for that column depends on the fraction of the column that is made of letters from this group. Black: 100% similar, dark-grey 80->100% similar, lighter grey: 60%-80% similar, white: less than 60% similar. All letters from this group are assigned this one color, and all letters outside of the group are not colored.

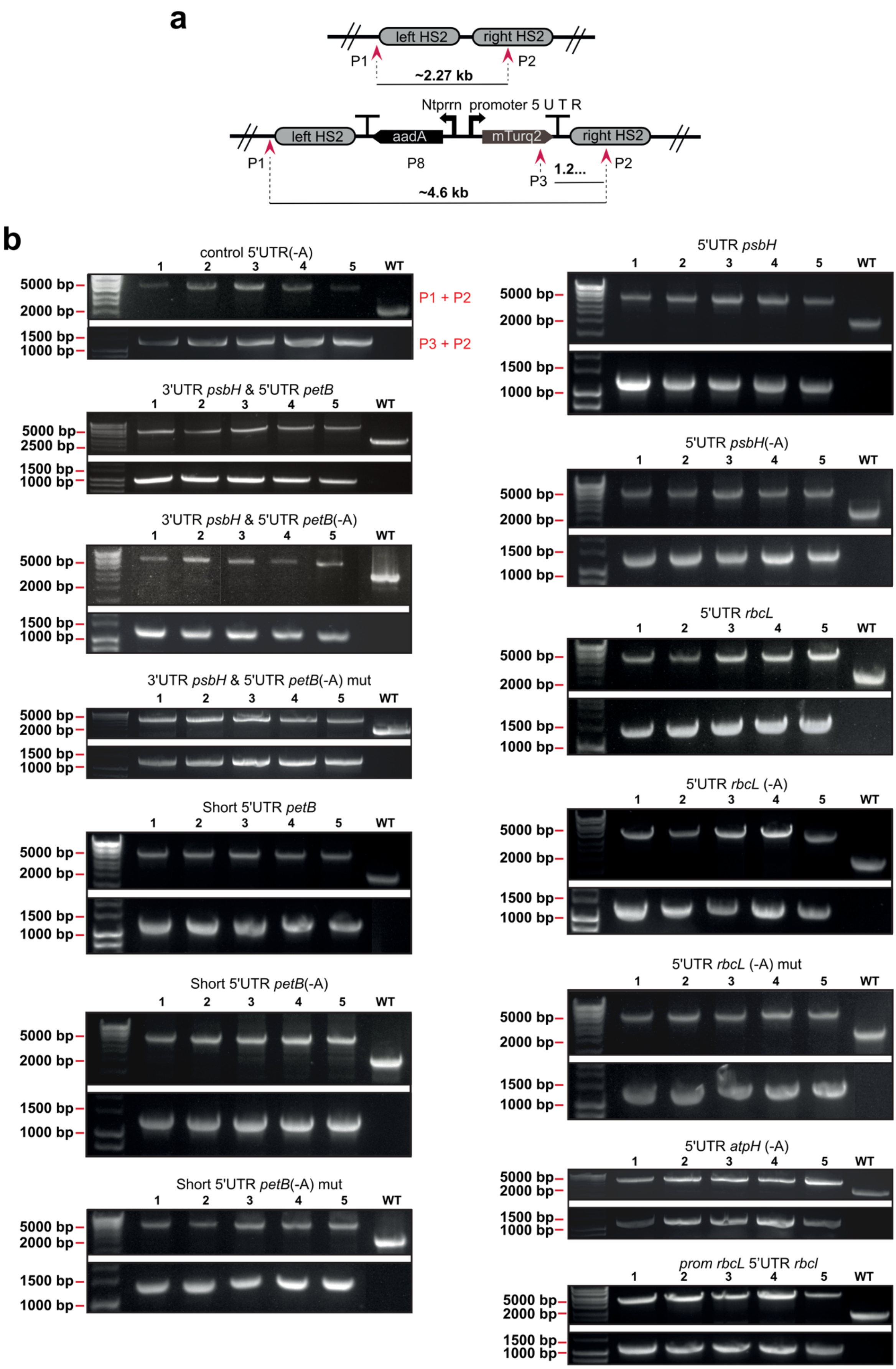
a
left HS2
right HS2
P1
P2
~2.27 kb
Ntprrn promoter 5 U T R
left HS2
aadA
mTurq2
right HS2
P1
P8
P3
1.2...
P2
~4.6 kb
b
control 5'UTR(-A)
P1 + P2
P3 + P2
3'UTR psbH & 5'UTR petB
3'UTR psbH & 5'UTR petB(-A)
3'UTR psbH & 5'UTR petB(-A) mut
Short 5'UTR petB
Short 5'UTR petB(-A)
Short 5'UTR petB(-A) mut
5'UTR psbH
5'UTR psbH(-A)
5'UTR rbcL
5'UTR rbcL (-A)
5'UTR rbcL (-A) mut
5'UTR atpH (-A)
prom rbcL 5'UTR rbcl
1 2 3 4 5 WT
5000 bp
2500 bp
2000 bp
1500 bp
1000 bp

**Supplementary Fig. 5**

Validation of DNA parts and vectors for chloroplast transformation. a) Schematic representation of the *rbcL-trnR* target region (flanked by left homologous sequence 2 (HS2) and right homologous sequence 2 (HS2) (Sauret-Güeto et al. 2020)) in the wild type chloroplast genome (top) and the same region after integration of the DNA construct II (bottom). Red arrowheads indicate the position of the PCR primers used for the detection of wild type or homoplasmic transplastomic lines. Maps not to scale. b) PCR analysis of genomic DNA isolated from wild type and transplastomic plants. Homoplasmy (bottom, primers P1+P2) and Integrity of the reporter gene (top, primers P2+P3) were confirmed for transplastomic lines after 2 months of subculture under selective conditions. The primer pair used for each PCR are shown next to the gel images.

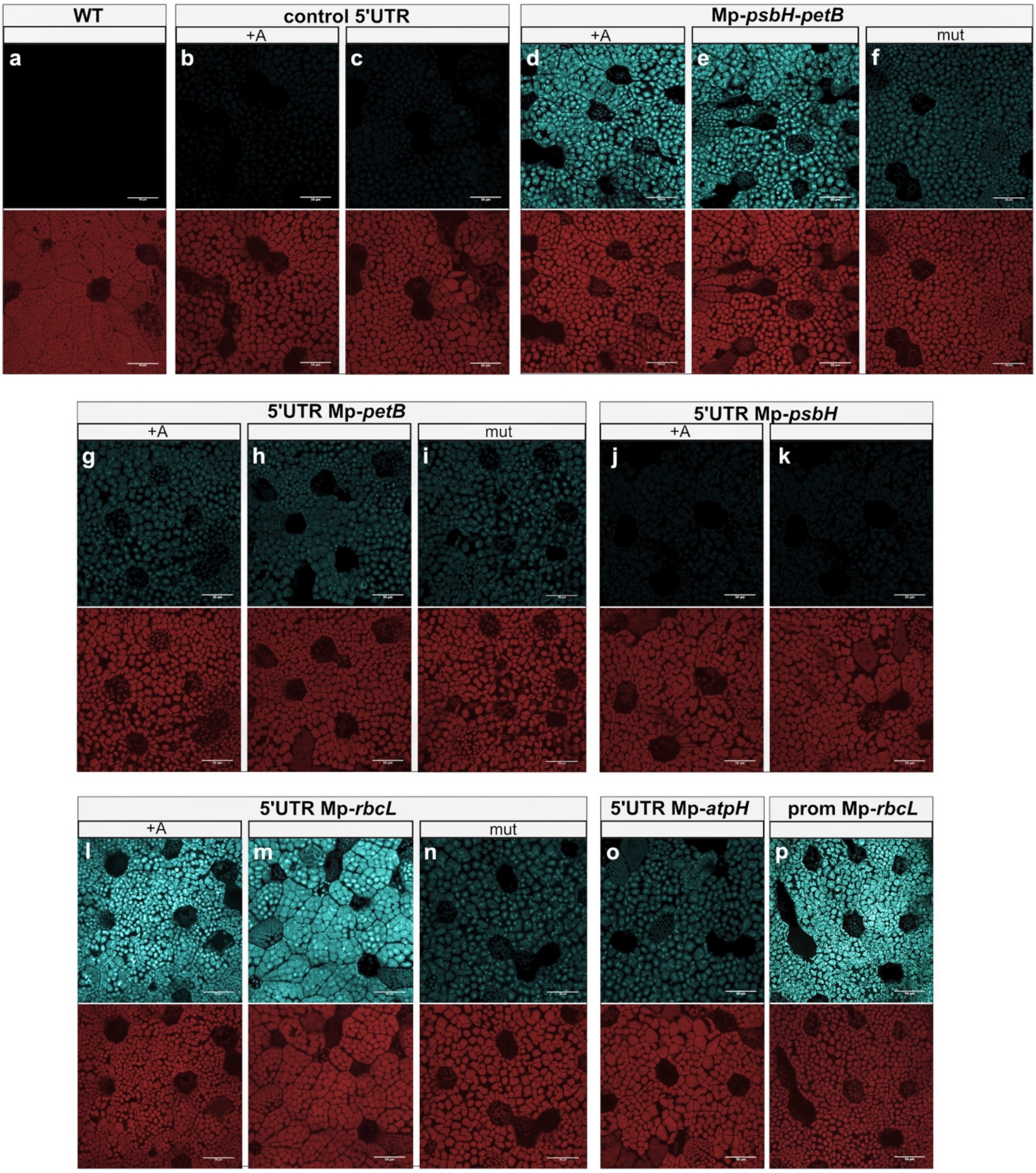
WT
a
control 5'UTR
+A
b
c
Mp-psbH-petB
+A
mut
d
e
f
5'UTR Mp-petB
+A
mut
g
h
i
5'UTR Mp-psbH
+A
j
k
5'UTR Mp-rbcL
+A
mut
l
m
n
5'UTR Mp-atpH
o
prom Mp-rbcL
p

**Supplementary Fig. 6**

Microscopy images of *Marchantia* transplastomic 3 day gemmae expressing the mTurq2cp fluorescent protein under the control of the Nt-*psbA* promoter fused to different candidate stabilisation sequences: control 5'UTR, Mp-*psbH-petB,* 5'UTR Mp-*petB*, 5'UTR Mp-*psbH,* 5'UTR Mp-*rbcL* and 5'UTR Mp-*atpH*. +A: Adenine introduced by the common syntax present between the 5'UTR and the mTurq2cp coding sequence. Mut: predicted PPR binding sequence mutated. Panel top: Chlorophyll autofluorescence channel, Panel bottom: mTurq2cp channel. All images acquired using identical instrument settings. 5'UTR Mp-*rbcL* confers the highest levels of expression followed by Mp-*psbH*-*petB*. 5'UTR Mp-*petB* and 5'UTR Mp-*atpH.* 5'UTR Mp-*psbH* expression levels are similar to those of the control 5'UTR. The addition of an extra "A" between the 5'UTR and the mTurq2cp coding sequence does not affect the expression of mTurq2cp. Expression levels are reduced for both when the predicted PPR binding sequence is mutated.

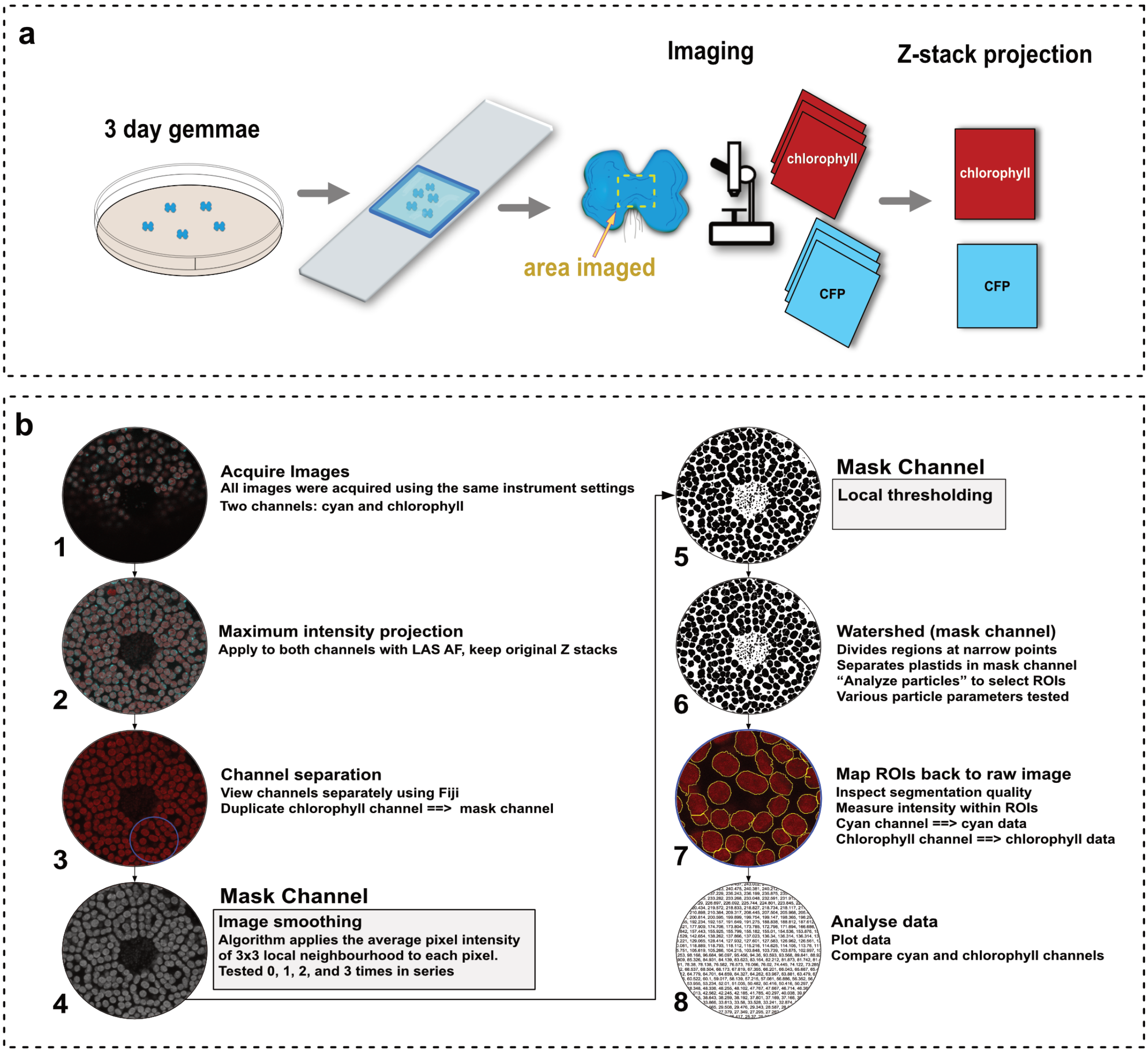

**c**

Macro (plain text between the bold start and stop lines)

```
run("Duplicate...", " ");
run("Smooth");
run("Smooth");
run("Auto Local Threshold", "method=Phansalkar radius=15 parameter_1=0 parameter_2=0 white");
run("Watershed");
run("Clear Results");
run("Analyze Particles...", "size=250-1500 circularity=0.60-1.00 display exclude clear add");
close();
run("Clear Results");
roiManager("Measure");
String.copyResults();
run("Clear Results");
\\ end
```

**Supplementary Fig. 7**: **Schematic of sample preparation and plastid segmentation pipeline**

a) Gemmae were plated on half strength Gamborg B5 1.2% (w/v) agar plates and placed in a growth cabinet for 3 days under continuous light at 21 $^{o}$C. A gene frame was positioned on a glass slide and 30 µL of half strength Gamborg B5 1.2% (w/v) agar were placed within the gene frame. 5 gemmae are then placed within the media filled gene frame, 30 µL of milliQ water are added and then a cover slip is used to seal the geneframe. Plants were then imaged immediately using an SP8 fluorescent confocal microscope.

b) Image processing pipeline. Steps 1 and 2 were performed on the Leica SP8, steps 3-7 were performed in Fiji using a custom macro (included in supplement), step 8 was performed using Google Sheets for spreadsheet data-preparation and RStudio for statistical analysis. All images were acquired using identical instrument settings. Images consisted of 16 Z stacks of 3 µm thickness.

c) Fiji Macro.

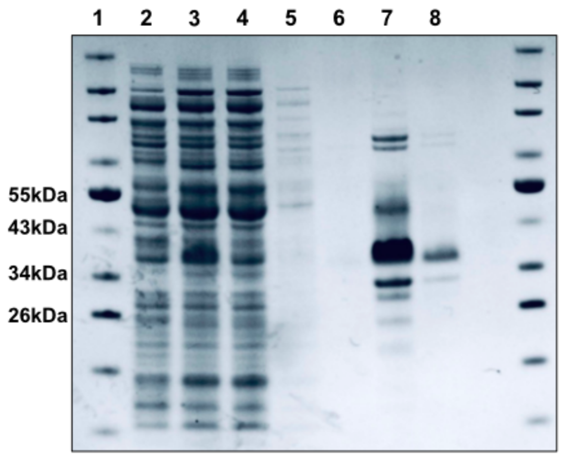

**Supplementary Fig. 8**
**Coomassie Blue–Stained Protein Gel of mTurquoise2 recombinant protein and its purification**.
Lane 1 PageRuler Pre-stained NIP protein ladder (#26635, ThermoFisher); lane 2 total cell extract from no transformed cells; lane 3 total extract from IPTG-induced bacteria cells for protein expression; lane 4 flow-through in the first step of the protein purification; lane 5 and 6, washes 1 and 2 respectively. Lane 7, purified mTurquoise2 recombinant protein in the first and second (lane 8) elution fraction.

| Sample | Total Reads | Forward mapped reads | Reverse mapped reads |
|---|---|---|---|
| A-TEX | 10.636.145 | 6,310,844 | 3,566,280 |
| A | 9.988.769 | 6,208,786 | 5,291,442 |
| B-TEX | 8.517.797 | 5,109,165 | 1,883,046 |
| B | 9.063.993 | 2,303,814 | 4,747,933 |

**Supplementary Table 1**

**Supplementary Table 2**: List of TSSs identified using dRNAseq
*Separate excel file*

**Supplementary Table 3**: List of MEME identified promoter motifs
*Separate excel file*

| | Liverworts | Order | Family | Genbank Accesion |
|---|---|---|---|---|
| 1 | *Cheilolejeunea xanthocarpa* | Porellales | Lejeuneaceae | MH064504 |
| 2 | *Cololejeunea lanciloba* | Porellales | Lejeuneaceae | MH064505 |
| 3 | *Schistochila macrodonta* | Jungermanniales | Schistochilaceae | MH064506 |
| 4 | *Porella perrottetiana* | Porellales | Porellaceae | MH064507 |
| 5 | *Radula japonica* | Porellales | Radulaceae | MH064508 |
| 6 | *Jubula hutchinsiae* | Porellales | Jubulaceae | MH064509 |
| 7 | *Frullania nodulosa* | Porellales | Frullaniaceae | MH064510 |
| 8 | *Plagiochila chinensis* | Jungermanniales | Plagiochilaceae | MH064511 |
| 9 | *Bazzania praerupta* | Jungermanniales | Lepidoziaceae | MH064512 |
| 10 | *Scapania ciliata* | Jungermanniales | Scapaniaceae | MH064513 |
| 11 | *Calypogeia fissa* | Jungermanniales | Calypogeiaceae | MH064514 |
| 12 | *Heteroscyphus argutus* | Jungermanniales | Lophocoleaceae | MH064515 |
| 13 | *Lunularia cruciata* | Marchantiales | Lunulariaceae | this study |
| 14 | *Lejeunea sp* | Porellales | Lejeuneaceae | this study |
| 15 | *Spruceanthus planifolius* | Porellales | Lejeuneaceae | this study |
| 16 | *Acrolejeunea sandvicensis* | Porellales | Lejeuneaceae | this study |
| 17 | *Rhaphidolejeunea foliicola* | Porellales | Lejeuneaceae | this study |
| 18 | *Ptychanthus striatus* | Porellales | Lejeuneaceae | this study |
| 19 | *Lopholejeunea zollingeri* | Porellales | Lejeuneaceae | this study |
| 20 | *Herbertus javanicus* | Jungermanniales | Herbertaceae | this study |
| 21 | *Delavayella serrata* | Jungermanniales | Delavayellaceae | this study |
| 22 | *Cyathodium smaragdinum* | Marchantiales | Cyathodiaceae | this study |
| 23 | *Fossombronia myrioides* | Fossombroniales | Fossombroniaceae | this study |
| 24 | *Hattoria yakushimensis* | Jungermanniales | Scapaniaceae | this study |
| 25 | *Notoscyphus lutescens* | Jungermanniales | Notoscyphaceae | this study |
| 26 | *Haplomitrium blumei* | Haplomitriales | Haplomitriaceae | MH064516 |
| 27 | *Pellia endiviifolia* | Pelliales | Pelliaceae | NC_019628.1 |
| 28 | *Ptilidium pulcherrimum* | Ptilidiales | Ptilidiaceae | HM222519.1 |
| 29 | *Marchantia polymorpha* | Marchantiales | Marchantiaceae | MH635409.1 |
| | **Mosses** | **Order** | **Family** | |
| 30 | *Bryum argenteum* | Bryales | | this study |
| 31 | *Sphagnum palustre* | Sphagnales | | this study |
| 32 | *Atrichum angustatum* | Polytrichopsida | | this study |
| 33 | *Tetraphis pellucida* | Tetraphidopsida | | this study |
| 34 | *Funaria hygrometrica* | Funariales | Funariaceae | this study |
| 35 | *Entosthodon attenuatus* | Funariales | Funariaceae | this study |
| 36 | *Bartramia pomiformis* | Bartramiales | | this study |
| 37 | *Ulota hutchinsiae* | Orthotrichales | Orthotrichaceae | this study |
| 38 | *Orthotrichum stellatum* | Orthotrichales | Orthotrichaceae | this study |
| 39 | *Ptychomnion cygnisetum* | Ptychomniales | | this study |
| 40 | *Hypnum imponens* | Hypnales | Callicladiaceae | this study |
| 41 | *Anomodon attenuatus* | Hypnales | Anomodontaceae | this study |
| 42 | *Andreaea wangiana* | Andreaeopsida | | this study |
| 43 | *Amphidium sp* | Orthotrichales | | this study |
| 44 | *Schistidium sp* | Grimmiales | | this study |
| 45 | *Tayloria sp* | Splachnales | | this study |
| 46 | *Physcomitrella patens* | Funariales | Funariaceae | NC_005087.2 |
| | **Hornworts** | **Order** | **Family** | |
| 47 | *Megaceros flagellaris* | Dendrocerotales | Megaceros | this study |
| 48 | *Anthoceros agrestis* | Anthocerotales | Anthoceros | NC_049002.1 |
| 49 | *Anthoceros punctatus* | Anthocerotales | Anthoceros | NC_049001.1 |
| 50 | *Leiosporoceros dussii* | Leiosporocerotales | Leiosporoceros | NC_039750.1 |
| 51 | *Nothoceros aenigmaticus* | Dendrocerotales | Nothoceros | NC_020259.1 |

**Supplementary Table 4: Bryophyte plastid genomes used in this study**

| Name | assembly | length bp |
|---|---|---|
| ***Anomodon attenuatus*** | complete | 115682 |
| ***Atrichum angustatum*** | complete | 116350 |
| ***Bartramia pomiformis*** | complete | 116167 |
| ***Bryum argenteum*** | complete (small repeats) | 114181 |
| ***Entosthodon attenuatus*** | complete (one repeat) | 113759 |
| ***Funaria hygrometrica*** | complete (one repeat) | 113244 |
| ***Hypnum imponens*** | complete | 115808 |
| ***Orthotrichum stellatum*** | complete | 113708 |
| ***Ptychomnion cygnisetum*** | complete | 114393 |
| ***Sphagnum palustre*** | complete | 128848 |
| ***Tetraphis pellucida*** | complete | 117992 |
| ***Ulota hutchinsiae*** | complete (not circular) | 114069 |

**Supplementary Table 5: Moss genome assemblies overview**

| | Sequence description | Nucleotide sequence cloned |
|---|---|---|
| | PROM Nt-psbA | agcggccaattcgagctcttggttgacacgagtatataagtcatgttatactgttgaataa |
| | control 5'UTR | |
| | Mp-psbH-petB | TAAATAAAAAATAAAAAAATGAATTGCTGCTAAAATAGCAGCAATTCATTTTTTTATTTTAGGTAGTTTAATTGTG TAATTATTAAATTCAAGGATTTTTGAAT |
| | Mp-psbH-petB mut | TAAATAAAAAATAAAAAAATGAATTGCTGCTAAAATAGCAGCAATTCATTTTTTTATTTTAGGTAGGGGCCGGCG CTAATTATTAAATTCAAGGATTTTTGAAT |
| | 5'UTR Mp-petB | CATTTTTTTATTTTAGGTAGTTTAATTGTGTAATTATTAAATTCAAGGATTTTTGAAT |
| | 5'UTR Mp-petB mut | CATTTTTTTATGAGCTGTAGGGGCCGGCGCTAATTATTAAATTCAAGGATTTTTGAAT |
| | 5'UTR Mp-psbH | TTAGGTAGTTCCAAAATAATGGTATATTTTTAATGTATATTTTTTAATATAATAGTACAAAAAGTTAATAAATCTC AACTAATCTGATAAGTTTT |
| | 5'UTR Mp-rbcL | AGAAAAAAAATTTTTATCGAGCAGACCTCATACTTGCAAGAATATTATTTGATTTGTAGGGAGGGACTT |
| | 5'UTR Mp-rbcL mut | AGAAAAAAAATTTTTATCTCATAtACAGGCGACTtgcaaGAATATTATTTGATTTGTAGGGAGGGACTT |
| | 5'UTR Mp-atpH | AAAAAAGAGACACTTTGAGTTATTAACTGCTTTAATTAAAATATTTTTATAAAAAATTTTTAGTAAGCAACGAAATA GATTTTTAATAAAATCTTTTTTGCAAATTTAGTTAAAGGAGATTATC |
| | prom Mp-rbcL | TTGATTTAATAATAAAATAACTAGGTTGCATTACATATAAAAAACAATATACAATAATAATGTTTTATTATTGGAA AAAATTTTTACTTAAAAAATTTTTTATACAAAAGAAAAATTAGAAAAAAAATTTTTATCGAGCAGACCTCATACTTG CAAGAATATTATTTGATTTGTAGGGAGGGACTT |
| | **Construct description** | **Notes** |
| 1 | pNt-psbA-control 5'UTR:mTurq2cp | Sauret-Gueto et al., 2020 |
| 2 | pNt-psbA-control 5'UTR (ATGg):mTurq2cp | |
| 3 | pNt-psbA-Mp-psbH-petB:mTurq2cp | |
| 4 | pNt-psbA-Mp-psbH-petB(ATGg):mTurq2cp | |
| 5 | p-Nt-psbA-Mp-psbH-petB(mut_ATGg)::mTurq2cp | mutated 10 bp (TTTAATTGTG->GGGCCGGCGC) |
| 6 | p-Nt-psbA-5'UTR-Mp-petB:mTurq2cp | |
| 7 | p-Nt-psbA-5'UTR-Mp-petB(ATGg):mTurq2cp | |
| 8 | p-Nt-psbA-5'UTR-Mp-petB(mut_ATGg):mTurq2cp | mutated 15 bp (TTTAGGTAGTTTAATTGTG-> GAGCTGTAGGGGCCGGCGC) |
| 9 | p-Nt-psbA-5'UTR-Mp-psbH:mTurq2cp | |
| 10 | p-Nt-psbA-5'UTR-Mp-psbH(ATGg):mTurq2cp | |
| 11 | p-Nt-psbA-5'UTR-Mp-rbcL:mTurq2cp | |
| 12 | p-Nt-psbA-5'UTR-Mp-rbcL(ATGg):mTurq2cp | |
| 13 | p-Nt-psbA-5'UTR-Mp-rbcL(mut_ATGg):mTurq2cp | mutated 10bp (GAGCAGACCTCAT->TCATAtACAGGCG) |
| 14 | p-NtpsbA-5U-MpatpH(ATGg):mTurq2cp | |
| 15 | p-Mp-rbcL:mTurq2cp | |

#### Supplementary Table 6: List of constructs

#### Supplementary Table 7: Microscopy image quantification *Separate excel file*

| Description - For amplifcation of: | Part type | Fw primer | 5'->3' | Rev primer | 5'->3' | Template |
|---|---|---|---|---|---|---|
| **Mp cam1 rbcL promoter region** | | EF240 | CCAACACCTGCTTTAGTCTCCG | EF239 | GGAAAAATTGTTAAGGTCAATGTGTATC | gDNA |
| **Mp cam1 rbcL promoter region domesticated** | PROM5 | EF616 | TAGAGCTCTTCGTCTCTGGAGTTGATTTAATAATAAAATAACTAGG | EF269 | ACTAGCTCTTCGTCTCACATTAAGTCCCTCCCTACAAATC | Mp cam1 rbcL promoter region |
| **Mp cam1 atpH promoter region** | | EF223 | AGGCGAATCTATGGAAGGGCAT | EF224 | CTTCTGCTTCAGGCTGTCTTGC | gDNA |
| **5'UTR Mp-atpH** | 5UTR | AT001 | taGAGCTCTTCgtctcTTACTAAAAAAGAGACACTTTGAGTTATTAACTGC | EF610 | AGCAGCTCTTCGTCTCTCCATGATAATCTCCTTTAACTAAATTTGCAAAAAAG | Mp cam1 atpH promoter region |
| **Mp-psbH-petB** | 5UTR | AT012 | taGAGCTCTTCgtctcTTACTTAAATAAAAAATAAAAAAATGAATTGCTGCTAAAATAGC | AT011 | AGCAGCTCTTCGTCTCTCATTATTCAAAAATCCTTGAATTTAATAATTACACAATTAAAC | gDNA |
| **Mp-psbH-petB (-A)** | 5UTR-A | AT012 | taGAGCTCTTCgtctcTTACTTAAATAAAAAATAAAAAAATGAATTGCTGCTAAAATAGC | EF612 | AGCAGCTCTTCGTCTCTcCATATTCAAAAATCCTTGAATTTAATAATTACAC | Mp-psbH-petB |
| **Mp-psbH-petB** | 5UTR | AT012 | taGAGCTCTTCgtctcTTACTTAAATAAAAAATAAAAAAATGAATTGCTGCTAAAATAGC | EF615 | AGCAGCTCTTCGTCTCTCCATATTCAAAAATCCTTGAATTTAATAATTAGCGCCGGCCCCTACCTAAAA | Mp-psbH-petB |
| **5'UTR Mp-petB** | 5UTR | PP01 | taGAGCTCTTCgtctcTTACTCATTTTTTTATTTTAGGTAGTTTAATTGTG | PP02 | AGCAGCTCTTCGTCTCTCATTATTCAAAAATCCTTGAATTTAATAATTAC | gDNA |
| **5'UTR Mp-petB** | 5UTR-A | PP01 | taGAGCTCTTCgtctcTTACTCATTTTTTTATTTTAGGTAGTTTAATTGTG | PP03 | AGCAGCTCTTCGTCTCTcCATATTCAAAAATCCTTGAATTTAATAATTAC | 5'UTR Mp-petB |
| **5'UTR Mp-petB mutated** | 5UTR-A | PP04 | taGAGCTCTTCgtctcTTACTCATTTTTTTATGAGCTGTAG | PP05 | AGCAGCTCTTCGTCTCTcCATATTCAAAAATCCTTGAATTTAATAATTAGCGCCGGCCC | 5'UTR Mp-petB |
| **5'UTR Mp-psbH** | 5UTR | PP06 | taGAGCTCTTCgtctcTTACTTATAATAGTACAAAAAGTTAATAAATC | PP07 | AGCAGCTCTTCGTCTCTCATTAAAACTTATCAGATTAGTTG | gDNA |
| **5'UTR Mp-psbH** | 5UTR-A | PP06 | taGAGCTCTTCgtctcTTACTTATAATAGTACAAAAAGTTAATAAATC | PP08 | AGCAGCTCTTCGTCTCTcCATAAAACTTATCAGATTAGTTG | 5'UTR Mp-psbH |
| **5'UTR Mp-rbcL** | 5UTR | MRL1-F | TAGAGCTCTTCGTCTCTTACTAGAAAAAAATTTTTATCGAGCAGACCTC | MRL1-R1 | AGCAGCTCTTCGTCTCTCATTAAGTCCCTCCCTACAAATCAAATAATATTCTTG | gDNA |
| **5'UTR Mp-rbcL** | 5UTR-A | MRL1-F | TAGAGCTCTTCGTCTCTTACTAGAAAAAAATTTTTATCGAGCAGACCTC | MRL1-R2 | AGCAGCTCTTCGTCTCTCCATAAGTCCCTCCCTACAAATCAAATAATATTCTTG | 5'UTR Mp-rbcL |
| **5'UTR Mp-rbcL mutated** | 5UTR-A-SCR | MRL1-scr-F3 | TAGAGCTCTTCGTCTCTTACTAGAAAAAAATTTTTATCTCATATACAGGCGACTTG | MRL1-R3 | AGCAGCTCTTCGTCTCTCCATAAGTCCCTCCCTACAAATCAAATAATATTC | 5'UTR Mp-rbcL |
| **Nt-psbA (-A)** | PROM5-A | AT004 | taGAGCTCTTCgtctcTGGAGagcggccaattcgagctc | EF614 | AGCAGCTCTTCGTCTCTcCATTTTTTCCTCCGGATCCCCCGG | pNt-psbA-control 5'UTR:mTurq2cp |
| **mTurq2cp (-A)** | CDS-A | EF613 | taGAGCTCTTCgtctcTatggtttctaaaggagaagaactttttactg | EF611 | CAGCTCTTCGTCTCTAAGCttatttgtacaattcatccattcctaatg | pNt-psbA-control 5'UTR:mTurq2cp |
| **Nt-psbA (-A)** | PROM | AT004 | TAGAGCTCTTCGTCTCTGGAGAGCGGCCAATTCGAGCTC | AT005c | AGCAGCTCTTCGTCTCTAGTATTATTCAACAGTATAACATGACTTATATACTCG | pNt-psbA-control 5'UTR:mTurq2cp |

| | | | | | |
|---|---|---|---|---|---|
| **Genotyping primers** | | | | | |
| | | KM14 (P1) | CGGAGACTAAAGCAGGTGTTGG | EF513 (P2) | ACAGATCCCATACTACCGCC |
| **P1, P2 and P3 in Supp Fig. 5** | | | | | |
| | | EF344 (P3) | ttacctgtaccatggcctac | | |

**Supplementary Table 8: List of primers used in this study**

Edgar, Robert C. 2004. “MUSCLE: A Multiple Sequence Alignment Method with Reduced Time and Space Complexity.” *BMC Bioinformatics* 5 (August): 113.

Emms, David M., and Steven Kelly. 2015. “OrthoFinder: Solving Fundamental Biases in Whole Genome Comparisons Dramatically Improves Orthogroup Inference Accuracy.” *Genome Biology* 16 (August): 157.

Lohse, Marc, Oliver Drechsel, Sabine Kahlau, and Ralph Bock. 2013. “OrganellarGenomeDRAW—a Suite of Tools for Generating Physical Maps of Plastid and Mitochondrial Genomes and Visualizing Expression Data Sets.” *Nucleic Acids Research* 41 (W1): W575–81.

Sauret-Güeto, Susanna, Eftychios Frangedakis, Linda Silvestri, Marius Rebmann, Marta Tomaselli, Kasey Markel, Mihails Delmans, Anthony West, Nicola J. Patron, and Jim Haseloff. 2020. “Systematic Tools for Reprogramming Plant Gene Expression in a Simple Model,.” *ACS Synthetic Biology* 9 (4): 864–82.
