## Supplementary material for "Comparative analysis of early divergent land plants and construction of DNA tools for hyper-expression in *Marchantia* chloroplasts": transcription data

| | Start | End | strand of gene | | Gene name | Gene function | starnd of trascript | Genomic coordinates of TSS | TSS relatively to ATG | Transcipt relatively to gene | TSS type | TSS group | Control_fwd | TEX_fwd | Control_rev | TEX_rev |
|---|---|---|---|---|---|---|---|---|---|---|---|---|---|---|---|---|
| 1 | 93 | 854 | + | | rps12 | ribosomal protein S12 | + | 1 | | | | 1 | 14.0638317 | 20.7539794 | 0 | 0 |
| | 93 | 324 | | exon 1 | | | + | 111 | | rps12 intron 1 | | 1 | 13.90756687 | 18.07149224 | 0.781323982 | 0.423550599 |
| | 829 | 854 | | exon 2 | | | + | 496 | | rps12 exon2 | | 2 | 24.22104343 | 15.24782157 | 0 | 0 |
| 2 | 904 | 1371 | + | gene | rps7 | 30S ribosomal protein S7 | + | 1303 | -209 | rps7 inter (ndhB) | | 2 | 20.47068832 | 9.035746118 | 0.625059185 | 0.847101199 |
| 3 | 1512 | 3535 | + | gene | ndhB (ndh2) | NADH dehydrogenase subunit 2 | + | 1696 | | ndhB exon 1-inter | | 2 | 3.594090315 | 2.964854195 | 0.468794389 | 0.282367066 |
| | | | | | | | + | 2209 | | ndhB exon 1-inter | | 2 | 46.09811492 | 4.09432246 | 0.468794389 | 0.423550599 |
| | 1512 | 2237 | | exon 1 | | | + | 2756 | | ndhB exon 2 -inter | | 2 | 43.28534858 | 4.800240125 | 0.312529593 | 0.282367066 |
| | 2756 | 3535 | | exon 2 | | | + | 3090 | | ndhB exon 2 -inter | | 2 | 17.50165719 | 15.95373924 | 0.312529593 | 0 |
| | | | | | | | + | 3192 | | ndhB exon 2 -inter | | 2 | 140.6383167 | 117.6058831 | 2.812766334 | 1.694202397 |
| | | | | | | | – | 3462 | | ndhB exon 2 -inter | | 3 | 1.406383167 | 1.129468265 | 1.406383167 | 3.67077186 |
| 4 | 3633 | 3712 | + | tRNA | tRNA-Leu-CAA | | + | 3610 | -23 | tRNA-Leu | | 1 | 77.19480938 | 84.85130339 | 0.625059185 | 0 |
| | | | | | | | + | 3633 | 1 | tRNA-Leu | | 1 | 359.5652963 | 649.0207016 | 0.781323982 | 0 |
| 5 | 3954 | 4058 | + | gene | psbM | photosystem II protein M | + | 3775 | -179 | psbM | | 2 | 20.47068832 | 12.00060031 | 0 | 0.141183533 |
| | | | | | | | + | 3954 | | psbM-0 | | 2 | 69.22530477 | 28.37789015 | 0.156264796 | 0 |
| 6 | 4182 | 5073 | + | gene | ycf66 | hypothetical protein | | | | | | | | | | |
| | 4182 | 4287 | | exon 1 | | | | | | | | | | | | |
| | 4772 | 5073 | | exon 2 | | | | | | | | | | | | |
| 7 | 5202 | 5113 | - | gene | ycf (ORF29) | | – | 5299 | | orf29 inter | | 1 | 0 | 0 | 3.125295926 | 3.67077186 |
| | | | | | | | – | 5406 | -204 | orf29 | | 2 | 313.779711 | 266.4133269 | 78.75745735 | 44.89636352 |
| 8 | 5601 | 5531 | - | tRNA | tRNA-Cys-GCA | | – | 5601 | 0 | tRNA-cys | | 1 | 1.71891276 | 1.976569463 | 752.8837887 | 3263.457367 |
| | | | | | | | – | 5613 | -12 | tRNA-cys | | 1 | 0.937588778 | 0.988284732 | 289.7149324 | 292.9558312 |
| 9 | 5740 | 8937 | + | gene | rpoB | RNA polymerase beta subunit | + | 5405 | -191 | rpoB? | | 2 | 311.2794743 | 263.3072892 | 78.44492775 | 44.89636352 |
| | | | | | | | – | 6111 | | rpoB-internal | | 2 | 0.781323982 | 0.282367066 | 7.65697502 | 6.494442522 |
| | | | | | | | – | 6574 | | rpoB-internal | | 2 | 0.781323982 | 0 | 3.281560723 | 2.823670662 |
| 10 | 8968 | 11627 | + | gene | rpoC1 | RNA polymerase beta' subunit | + | 8579 | | rpoB-internal,rpoC1? | | 1 | 100.6345288 | 151.489931 | 0 | 0.282367066 |
| | 8968 | 9399 | | exon 1 | | | – | 8887 | | rpoB-internal | | 1 | 0 | 0 | 3.437825519 | 5.788524857 |
| | 10005 | 11627 | | exon 2 | | | – | 9417 | | rpoC1 intron 1 | | 2 | 0.468794389 | 0.423550599 | 4.062884704 | 2.823670662 |
| | | | | | | | – | 9757 | | rpoC1 intron 1 | | 1 | 0 | 0 | 6.250591853 | 6.353258989 |
| 11 | 11700 | 15854 | + | gene | rpoC2 | RNA polymerase beta'' subunit | + | 12024 | rpoC1 intro | rpoC2-inter | | 1 | 100.6345288 | 134.547907 | 0.156264796 | 0.141183533 |
| | | | | | | | – | 12453 | | rpoC2-inter | | 2 | 0.468794389 | 0.141183533 | 30.78416488 | 30.63682668 |
| | | | | | | | – | 13106 | | rpoC2-inter | | 1 | 0 | 0 | 3.750355112 | 4.659056592 |
| | | | | | | | – | 13985 | | rpoC2-inter | | 1 | 0.156264796 | 0.141183533 | 2.656501537 | 2.682487129 |
| 12 | 15937 | 16644 | + | gene | rps2 | 30S ribosomal protein S2 | + | 14817 | -1120 | rpoC2-inter (rps2?) | | 1 | 5.93806226 | 6.917993121 | 0 | 0 |
| 13 | 16773 | 17519 | + | gene | atpI | ATP synthase CF0 A subunit | + | 16700 | -73 | atpI | | 1 | 2.812766334 | 3.106037728 | 0 | 0 |
| 14 | 17889 | 18134 | + | gene | atpH | ATP synthase CF0 C chain | + | 17748 | -141 | atpH? | | 2 | 372.3790096 | 43.48452819 | 0 | 0 |
| 15 | 18356 | 19483 | + | gene | atpF | ATP synthase CF0 B subunit | | | | | | | | | | |
| | 18356 | 18500 | | exon 1 | | | | | | | | | | | | |
| | 19074 | 19483 | | exon 2 | | | – | 19218 | | atpF-internal exon2 | | 1 | 5.781797464 | 4.09432246 | 2.96903113 | 8.612195518 |
| 16 | 19526 | 21049 | + | gene | atpA | ATP synthase CF1 alpha subunit | | | | | | | | | | |
| | | | | | | | – | 20068 | | atpA-internal | | 1 | 4.375414297 | 4.376689526 | 12.03238932 | 17.36557457 |
| 17 | 21184 | 21113 | - | tRNA | tRNA-Arg-UCU | | – | 21184 | 1 | tRNA-arg-UCU | | 1 | 6.406856649 | 7.482727254 | 400.9754674 | 2322.610303 |
| | | | | | | | – | 21295 | 1 | tRNA-Gly-UCU exon 2 | | 1 | 16.40780361 | 20.33042876 | 1527.488384 | 10761.85599 |
| 18 | 21899 | 21877 | - | tRNA | tRNA-Gly-UCC | exon 1 | – | 21899 | 1 | tRNA-Gly-UCC | | 1 | 0.156264796 | 0 | 2356.473129 | 11296.09448 |
| | 21295 | 21248 | | | tRNA-Gly-UCC | exon 2 | + | 21926 | | intergenic | | 2 | 10.93853574 | 2.258936529 | 0.625059185 | 6.494442522 |
| 19 | 22116 | 22015 | - | gene | ycf12 | | – | 22108 | | ycf12 internal | | 1 | 0.468794389 | 0.564734132 | 90.47731707 | 103.2051627 |
| | | | | | | | – | 22231 | | ycf45 internal | | 2 | 0.468794389 | 0.423550599 | 21.4082771 | 21.3187135 |
| 20 | 22278 | 22141 | - | gene | ycf45 | | – | 22317 | -39 | ycf45 | | 2 | 9.844682168 | 9.741663783 | 29.6903113 | 25.55421949 |
| 21 | 22372 | 22470 | + | gene | psaM | photosystem I protein M | + | 22246 | -126 | psaM? (ycf45 inter) | | 1 | 24.53357302 | 25.27185242 | 28.90898732 | 20.4716123 |
| | | | | | | | + | 22654 | -64 | tRNA-Ser GUC | | 2 | 76.72601499 | 51.95554018 | 0 | 0 |
| 22 | 22718 | 22805 | - | tRNA | tRNA-Ser-GCU | | + | 22718 | 1 | tRNA-Ser GUC | | 1 | 450.0426134 | 737.9663274 | 3.594090315 | 5.082607191 |
| 23 | 22933 | 22823 | - | gene | psbI | photosystem II protein I | – | 22976 | -43 | psbI | | 2 | 2.812766334 | 0.847101199 | 362.8468571 | 8.753379051 |
| 24 | 23430 | 23263 | - | gene | psbK | photosystem II protein K | – | 23515 | -85 | psbK | | 2 | 0.312529593 | 0.141183533 | 11.87612452 | 6.917993121 |
| 25 | 23629 | 23700 | + | tRNA | tRNA-Gln-UUG | | + | 23620 | -9 | tRNA-Gln UUG | | 1 | 387.0679005 | 588.4529659 | 1.406383167 | 1.694202397 |
| | | | | | | | – | 23780 | | intergenic | | 1 | 0.781323982 | 0.282367066 | 13.90756687 | 14.68308744 |
| | | | | | | | + | 23845 | | intergenic | | 1 | 72.0380711 | 100.8050426 | 0 | 0 |
| 26 | | | | | petD?? | | + | 24369 | -217 | petD | | 2 | 3.125295926 | 5.92970839 | 0.156264796 | 0 |
| 27 | 25563 | 25715 | + | gene | orf50 | | + | 25168 | -395 | ycf ORF50? | | 2 | 6.406856649 | 6.070891923 | 0.156264796 | 0 |
| 28 | 27948 | 27982 | + | | tRNA-Lys-UUU | | + | 25787 | -47 | tRNA-Lys UUU | | 1 | 581.7738367 | 713.2592092 | 0.312529593 | 0.282367066 |
| | | | | | | | – | 25997 | | intergenic | | 1 | 2.187707149 | 0.847101199 | 9.532152576 | 11.01231558 |
| 29 | 28128 | 29189 | + | gene | psbA | photosystem II protein D1 | + | 28074 | -54 | psbA | | 1 | 20707.89828 | 51473.53959 | 0 | 0 |
| | | | | | | | + | 28286 | | psbA internal | | 1 | 445.6671991 | 474.9414053 | 0.156264796 | 0.282367066 |
| 30 | 29353 | 29427 | + | tRNA | tRNA-His-GUG | | | | | | | | | | | |
| | | | | | | | – | 33822 | | integenic | | 2 | 0.156264796 | 0 | 8.125769409 | 7.482727254 |
| | | | | | | | + | 33844 | | intergenic | | 1 | 12.6574485 | 13.41243564 | 0 | 0 |
| 31 | 36152 | 36079 | - | tRNA | tRNA-Asp-GUC | | – | 36152 | 1 | tRNA-Asp | | 1 | 0 | 0.141183533 | 1662.501168 | 5530.441358 |
| 32 | 36313 | 36232 | - | tRNA | tRNA-Tyr-GUA | | – | 36180 | -28 | tRNA-Asp | | 1 | 0 | 0 | 5.156738279 | 11.71823325 |
| 33 | 36456 | 36384 | - | tRNA | tRNA-Glu-UUC | | – | 36466 | -10 | tRNA-Glu | | 1 | 0 | 0 | 4357.756375 | 5287.605681 |
| 34 | 36680 | 37798 | + | gene | cysA | sulfate ABC transporter protein | + | 36361 | -319 | CysA | | 1 | 4.844208686 | 4.941423658 | 39.53499347 | 26.11895362 |
| | | | | | | | | 36662 | -18 | CysA | | | 0 | 0.423550599 | 0.156264796 | 0 |
| | | | | | | | + | 38004 | -36 | tRNA-Thr GGU | | 1 | 114.5420957 | 191.7272379 | 0.156264796 | 0 |
| 35 | 38040 | 38111 | + | tRNA | tRNA-Thr-GGU | | + | 38040 | 1 | tRNA-Thr GGU | | 1 | 357.6901188 | 1454.755125 | 0.156264796 | 0.282367066 |
| 36 | 38539 | 39600 | + | gene | psbD | photosystem II protein D2 | + | 38291 | -248 | psbD | | 2 | 56.09906188 | 24.56593476 | 0 | 0 |
| | | | | | | | + | 38395 | -144 | psbD | | 2 | 62.97471292 | 6.494442522 | 0.937588778 | 0.141183533 |
| 37 | 39548 | 40969 | + | gene | psbC | photosystem II 44 kDa protein | + | 39541 | -7 | psbC | | 1 | 140.0132575 | 147.5367921 | 0 | 0 |
| 38 | 41178 | 41091 | - | tRNA | tRNA-Ser-UGA | | – | 41178 | 1 | tRNA-ser | | 1 | 14.37636126 | 13.41243564 | 44.22293736 | 108.0054028 |
| 39 | 41354 | 41542 | + | gene | psbZ | photosystem II protein Z | + | 41325 | -29 | psbZ | | 2 | 82.50781246 | 56.47341323 | 0.781323982 | 0.705917665 |
| | | | | | | | – | 41589 | | intergenic | | 2 | 1.093853574 | 7.059176654 | 174.5477775 | 147.1132415 |
| 40 | 41747 | 41817 | + | tRNA | tRNA-Gly-GCC | | + | 41747 | 1 | tRNA-Gly GCC | | 1 | 752.1024647 | 1392.21082 | 0.156264796 | 0.564734132 |
| 41 | 41947 | 41874 | - | tRNA | tRNA-fMet-CAU | | – | 41947 | 1 | tRNA-fMet-CAU | | 2 | 2.812766334 | 2.682487129 | 17046.92663 | 73955.18185 |
| | | | | | | | – | 41958 | -11 | tRNA-fMet-CAU | | 1 | 2.96903113 | 2.541303596 | 315.9674182 | 422.8446816 |
| 42 | 42353 | 42051 | - | gene | rps14 | 30S ribosomal protein S14 | – | not detected | | | | | | | | |
| 43 | 44646 | 42442 | - | gene | psaB | PSI P700 apoprotein A2 | – | not detected | | | | | | | | |
| | | | | | | | + | 45424 | | psaA internal | | 1 | 6.406856649 | 11.43586618 | 24.84610262 | 16.09492277 |
| 44 | 46925 | 44673 | - | gene | psaA | PSI P700 apoprotein A1 | – | 47074 | -149 | psaA | | 1 | 0.156264796 | 0 | 527.862482 | 749.2610101 |
| 45 | 48316 | 47207 | - | gene | pafI | photosystem I-related pafI Ycf3 | – | | | | | | | | | |
| | 48316 | 48193 | | exon 1 | | | – | | | | | | | | | |
| | 47586 | 47207 | | exon 2 | | | + | 47495 | | pafI inter | | 2 | 31.72175365 | 29.22499135 | 1.71891276 | 0.847101199 |
| | | | | | | | – | 48278 | | pafI inter | | 3 | 0 | 0 | 0.625059185 | 3.388404794 |
| 46 | 48563 | 48650 | + | tRNA | tRNA-Ser-GGA | | + | 48548 | -15 | tRNA-Ser GGA | | 1 | 87.50828594 | 102.499245 | 0.312529593 | 0 |
| | | | | | | | + | 48563 | 1 | tRNA-Ser GGA | | 1 | 419.7272429 | 2548.362772 | 0.625059185 | 0 |
| 47 | 49667 | 49059 | - | gene | rps4 | ribosomal protein S4 | + | 49325 | | rps4 internal | | 1 | 10.00094696 | 16.09492277 | 1.562647963 | 0.423550599 |
| | | | | | | | – | 49456 | | rps4 internal | | 1 | 0.156264796 | 0.282367066 | 9.688417372 | 10.02403085 |
| 48 | 49968 | 49896 | - | tRNA | tRNA-Thr-UGU | | – | 49968 | 1 | trnA-Thr | | 1 | 0 | 0.423550599 | 281.4328982 | 1295.92365 |
| | | | | | | | – | 50041 | -73 | trnA-Thr | | 2 | 0 | 0 | 74.53830785 | 51.95554018 |
| 49 | 50174 | 50208 | + | tRNA | tRNA-Leu-UAA | exon 1 | + | 50139 | -35 | tRNA-Leu UAA | | 1 | 326.2808947 | 351.6881809 | 0.156264796 | 0 |
| | | | | | | | + | | | | | | | | | |
| | | | | | | | – | 50804 | | integenic | | 1 | 0.468794389 | 0 | 3.906619908 | 5.364974257 |
| | 50523 | 50572 | | | | exon 2 | + | | | | | | | | | |
| | | | | | | | | 50855 | | integenic | | 2 | 8.594563798 | 5.92970839 | 0 | 0 |
| 50 | 50649 | 50721 | + | tRNA | tRNA-Phe-GAA | | + | 50649 | 1 | tRNA-Phe-GAA | | 1 | 209.0822975 | 572.7815937 | 0.625059185 | 0.705917665 |
| | | | | | | | + | 51683 | | ndhK inter | | 1 | 2.500236741 | 3.106037728 | 0.781323982 | 0.705917665 |
| 51 | 51393 | 50884 | - | gene | ndhJ | NADH dehydrogenase subunit J | – | 51356 | | internal ndhJ | | 2 | 0.312529593 | 0 | 8.438299001 | 7.76509432 |
| 52 | 52174 | 51437 | - | gene | ndhK | NADH dehydrogenase subunit K | – | 52245 | -71 | ndhK (internal ndhC) | | 2 | 0.156264796 | 0 | 28.75272252 | 22.73054883 |
| 53 | 52527 | 52165 | - | gene | ndhC | NADH dehydrogen | | | | | | | | | | |
| | | | | | | | – | 52728 | | tRNA Val | | 1 | 0.468794389 | 0.847101199 | 140.3257871 | 153.4665005 |
| 54 | 53292 | 53256 | - | tRNA | tRNA-Val-UAC | exon 1 | – | 53292 | | tRNA-Val UAC | | 1 | 0.156264796 | 0.141183533 | 1982.844001 | 2829.459187 |
| | 52728 | 52694 | | | tRNA-Val-UAC | exon 2 | – | 53330 | -38 | tRNA-Val UAC-- | | 1 | 0 | 0 | 1094.009839 | 1471.414782 |
| | | | | | | | + | 53430 | -10 | tRNA-Met CAU | | 1 | 144.0761422 | 170.549708 | 0.625059185 | 0.141183533 |
| 55 | 53440 | 53513 | + | tRNA | tRNA-Met-CAU | | + | 53440 | 1 | tRNA-Met CAU | | 1 | 364.8782994 | 840.606756 | 0.625059185 | 0.141183533 |
| 56 | 53994 | 53590 | - | gene | atpE | ATP synthase epsilon chain | | | | | | | | | | |
| 57 | 55478 | 54000 | - | gene | atpB | ATP synthase CF1 beta subunit | – | 55498 | -20 | atpB | | 1 | 0 | 0 | 17.65792198 | 49.97897071 |
| | | | | | | | – | 55748 | -270 | atpB | | 2 | 16.40780361 | 7.341543721 | 4.531679093 | 3.811955393 |
| 58 | 56027 | 57454 | + | gene | rbcL | rbcL | + | 55903 | -124 | rbcL | | 1 | 515.3612983 | 2357.482635 | 0 | 0 |
| 59 | 57573 | 57646 | + | tRNA | tRNA-Arg-CCG | | + | 57573 | 1 | tRNA-Arg CCG | | 1 | 102.3534416 | 254.6950937 | 0.468794389 | 0.141183533 |
| 60 | 57831 | 58778 | + | gene | accD | | + | 57700 | -131 | accD? | | 3 | 1.093853574 | 0.423550599 | 1.406383167 | 1.976569463 |
| 61 | 58954 | 59064 | + | gene | psaI | psaI | + | 58880 | -74 | psaI | | 2 | 16.09527402 | 5.364974257 | 0 | 0 |
| 62 | 59287 | 59841 | + | gene | pafII | photosystem I assembly protein Ycf4 | + | | | | | | | | | |
| | | | | | | | + | 59326 | | pafII inter | | 1 | 10.46974135 | 12.14178385 | 0.156264796 | 0.141183533 |
| | | | | | | | + | 59572 | | pafII inter 2/cemA? | | 1 | 26.56501537 | 32.89576321 | 0 | 0 |
| 63 | 59913 | 61217 | + | gene | cemA | envelope membrane protein | + | | | | | | | | | |
| 64 | 61403 | 62365 | + | gene | petA | apocytochrome f precursor | + | 61012 | | cemA inter - petA? | | 2 | 15.00142045 | 7.341543721 | 0.312529593 | 0.141183533 |
| 65 | 62680 | 62558 | - | gene | psbJ | PSII reaction center subunit X | – | 62708 | -28 | psbJ | | 2 | 0.156264796 | 0.141183533 | 126.8870146 | 81.74526566 |
| 66 | 62916 | 62800 | - | gene | psbL | PSII reaction center subunit XII | – | no detected | | | | | | | | |
| 67 | 63057 | 62938 | - | gene | psbF | photosystem II reaction center subunit VI | – | 63270 | -213 | psbF / psbE internal | | 1 | 0.312529593 | 0.282367066 | 52.66123636 | 71.58005128 |
| 68 | 63318 | 63067 | - | gene | psbE | cytochrome b559 alpha subunit | – | 63460 | -142 | psbE | | 1 | 0 | 0 | 376.7544239 | 864.3255896 |
| 69 | 63917 | 64012 | + | gene | petL | | + | 63772 | -145 | petL | | 3 | 0.625059185 | 0.988284732 | 0 | 0 |
| | | | | | | | + | 63876 | -41 | petL 2? | | 3 | 2.343971945 | 0.988284732 | 0 | 0 |
| 70 | 64134 | 64247 | + | gene | petG | cytochrome b6/f complex subunit V | + | 64088 | -46 | petG | | 3 | 2.031442352 | 1.694202397 | 0 | 0 |
| 71 | 64391 | 64318 | - | tRNA | tRNA-Trp-CCA | | – | 64391 | 1 | tRNA-Trp CCA | | 1 | 0.156264796 | 0.282367066 | 622.246419 | 3528.600042 |
| | | | | | | | + | 64445 | | | | 1 | 12.6574485 | 14.68308744 | 0.468794389 | 0.564734132 |
| 72 | 64544 | 64471 | - | tRNA | tRNA-Pro-UGG | | – | 64544 | 1 | tRNA-Pro UGG | | 1 | 0.468794389 | 1.129468265 | 1191.362807 | 1267.969311 |
| | | | | | | | – | 64559 | -15 | tRNA-Pro UGG | | 1 | 0.468794389 | 0.282367066 | 581.7738367 | 662.1507702 |
| | | | | | | | – | 64709 | -165 | tRNA-Pro UGG | | 2 | 0.156264796 | 0 | 9.219622983 | 4.376689526 |
| 73 | 64780 | 64908 | + | gene | psaJ | photosystem I subunit IX | + | 64732 | -48 | psaJ | | 2 | 31.09669447 | 27.24842189 | 0 | 0 |
| 74 | 65027 | 65224 | + | gene | rpl33 | ribosomal protein L33 | + | 64741 | -39 | psaJ 2/ rpl33? | | 1 | 82.35154766 | 105.4640992 | 0 | 0 |
| 75 | 65252 | 65479 | + | gene | rps18 | ribosomal protein S18 | + | 65033 | -219 | rpl33 inter (rps18) | | 2 | 3.594090315 | 3.106037728 | 0 | 0 |
| 76 | 65909 | 65559 | – | gene | rpl20 | 50S ribosomal protein L20 | – | not detected | | | | | | | | |
| | | | | | | | – | 66272 | | rps12 inter | | 1 | 0.156264796 | 0 | 15.78274443 | 18.63622637 |
| 77 | 66199 | 67098 | – | gene | rp112 | | – | 66447 | | rps12 intron | | 2 | 0.625059185 | 0.141183533 | 13.90756687 | 11.29468265 |
| | | | | | | | – | 66856 | rps12 | -47 | | 2 | 0.312529593 | 0 | 89.3834635 | 5.788524857 |
| 78 | 68388 | 66883 | - | gene | clpP | ATP-dependent Clp protease proteolytic subunit | – | 67134 | clpP inter | clp exon 3 | | 1 | 0 | 0.141183533 | 34.69078478 | 78.0744938 |
| | 68388 | 68318 | | exon 1 | | | – | 67805 | clpP inter | clp exon 2 | | 1 | 0.468794389 | 0.988284732 | 10.31347656 | 179.02072 |
| | 67805 | 67514 | | exon 2 | | | – | 68477 | clpP | -59 | | 1 | 0 | 0 | 0 | 0 |
| | 67134 | 66883 | | exon 3 | | | | | | | | | | | | |
| 79 | 68717 | 70243 | + | gene | psbB | photosystem II 47 kDa protein | 1 | 68573 | -144 | psbB | | 1 | 233.4596057 | 322.4631896 | 1.875177556 | 1.129468265 |
| 80 | 70361 | 70468 | + | gene | psbT | photosystem II protein T | + | | | | | | | | | |
| 81 | 70665 | 70534 | - | gene | psbN | photosystem II protein N | – | 70712 | -47 | psbN | | 2 | 1.562647963 | 1.553018864 | 1160.266113 | 926.8698947 |
| | | | | | | | – | 74208 | | rpoA internal | | 1 | 0 | 0.141183533 | 4.531679093 | 6.212075456 |
| 82 | 70763 | 70987 | + | gene | psbH | photosystem II protein H | + | 70726 | | psbH | | 2 | 13.12624289 | 6.917993121 | 0 | 0 |
| 83 | 71092 | 72222 | + | gene | petB | | + | 71049 | | petB | | 2 | 187.6740204 | 23.86001709 | 1.093853574 | 0.564734132 |
| | 71092 | 71097 | | exon 1 | | | | | | | | | | | | |
| | 71575 | 72222 | | exon 2 | | | | | | | | | | | | |
| 84 | 72369 | 73339 | + | gene | petD | | + | 71585 | | petB exon 2 internal | | 2 | 199.2376153 | 45.74346472 | 0.156264796 | 0.423550599 |
| | | | | | | | + | | | | | | | | | |
| | 72369 | 72376 | | exon 1 | | | | | | | | | | | | |
| | 72865 | 73339 | | exon 2 | | | | | | | | | | | | |
| 85 | 74480 | 73455 | + | gene | rpoA | RNA polymerase alpha subunit | – | | | | | | | | | |
| 86 | 74905 | 74513 | - | gene | rps11 | 30S ribosomal protein S11 | – | 74917 | -12 | rps11 | | 3 | 0.156264796 | 0 | 1.875177556 | 2.823670662 |
| 87 | 75066 | 74953 | - | gene | rpl36 | 50S ribosomal protein L36 | – | | | | | | | | | |
| 88 | 75337 | 75101 | - | gene | infA | translation initiation factor 1 | – | not detected | | | | | | | | |
| 89 | 75821 | 75423 | - | gene | rps8 | ribosomal protein S8 | – | not detected | | | | | | | | |
| 90 | 76271 | 75903 | - | gene | rpl14 | 50S ribosomal protein L14 | – | not detected | | | | | | | | |
| 91 | 77310 | 76367 | - | gene | rpl16 | ribosomal protein L16 | – | | | | | | | | | |
| | | | | | | | + | 77341 | | integenic | | 2 | 4.68794389 | 3.67077186 | 0 | 0.282367066 |
| 92 | 78021 | 77365 | - | gene | rps3 | 30S ribosomal protein S3 | – | 77713 | | rps3 internal | | 1 | 0.156264796 | 0 | 8.594563798 | 11.57704971 |
| | | | | | | | + | 78105 | | rpl22 | | 1 | 69.85036396 | 88.80444231 | 0.937588778 | 1.129468265 |
| 93 | 78431 | 78072 | - | gene | rpl22 | ribosomal protein L22 | – | not detected | | | | | | | | |
| 94 | 78727 | 78449 | - | gene | rps19 | 30S ribosomal protein S19 | – | not detected | | | | | | | | |
| | | | | | | | – | 80008 | | rpl12 int | | 1 | 0.156264796 | 0 | 23.12718986 | 25.13066889 |
| 95 | 80122 | 78765 | - | gene | rpl2 | 50S ribosomal protein L2 | – | 80190 | -68 | rpl23 inter (rpl2) | | 2 | 0.156264796 | 0 | 5.000473482 | 2.258936529 |
| 96 | 80434 | 80159 | - | gene | rpl23 | ribosomal protein L23 | – | 80473 | -39 | rpl23 | | 2 | 0 | 0 | 3.750355112 | 3.67077186 |
| 97 | 80670 | 80597 | - | tRNA | tRNA-Ile-CAU | | – | 80670 | 1 | tRNA-Ile CAU | | 1 | 0 | 0 | 52.66123636 | 135.2538247 |
| | | | | | | | – | 80711 | -41 | tRNA-Ile CAU | | 1 | 0 | 0 | 27.81513375 | 36.42535154 |
| | 80716 | 90620 | | repeat_region | | inverted repeat B | | | | | | | | | | |
| 98 | 81427 | 81498 | + | tRNA | tRNA-Val-GAC | | + | 81427 | 1 | tRNA-Val GAC | | 1 | 2.500236741 | 7.482727254 | 0 | 0 |
| 99 | 81722 | 83220 | + | rRNA | 16S ribosomal RNA | | + | 81572 | -150 | 16S rna -- | | 1 | 11.09480054 | 14.54190391 | 0.156264796 | 0 |
| | | | | | | | + | 81593 | -129 | 16S | | 1 | 932.7445693 | 2395.461006 | 0.156264796 | 0 |
| | | | | | | | + | 81687 | -35 | 16S rna - | | 2 | 737.7261034 | 214.8813374 | 0.312529593 | 0.141183533 |
| | | | | | | | + | 81722 | 1 | 16S rna | | 2 | 113999.8569 | 80000.51956 | 0 | 0.141183533 |
| | | | | | | | + | 83225 | | 16S internal | | 1 | 1224.334679 | 1374.704062 | 0 | 0 |
| 100 | 83441 | 83477 | + | tRNA | tRNA-Ile-GAU | exon 1 | + | 83441 | 1 | tRNA-Ile GUA | | 1 | 160.327681 | 364.5358824 | 0.156264796 | 0 |
| | 84270 | 84304 | | | | exon 2 | | 84305 | -76 | tRNA-Ala UGC | | 1 | 48.44208686 | 54.49684377 | 0 | 0 |
| 101 | 84381 | 84418 | + | tRNA | tRNA-Ala-UGC | exon 1 | + | 84381 | 1 | tRNA-Ala UGC | | 1 | 402.8506449 | 1090.078059 | 0 | 0 |
| | 85186 | 85220 | | | | exon 2 | + | 84650 | | tRNA-Ala UGC intron | | 1 | 1163.235144 | 1326.136926 | 0 | 0 |
| | | | | | | | + | 85154 | | tRNA-Ala UGC intron | | 1 | 1907.211839 | 2785.268741 | 0 | 0 |
| 102 | 85380 | 88197 | + | rRNA | | 23S ribosomal RNA | + | 85312 | -68 | 23S rna- | | 2 | 4276.186151 | 698.7173052 | 0 | 0 |
| 103 | 88305 | 88407 | + | rRNA | | 4.5S ribosomal RNA | + | 88302 | -3 | 4.5S rna | | 2 | 664.9067084 | 208.5280784 | 0 | 0 |
| 104 | 88633 | 88751 | + | rRNA | | 5S ribosomal RNA | + | 88630 | -3 | 5S rna | | 2 | 2326.626552 | 2284.914299 | 0 | 0 |
| | | | | | | | + | 88633 | 1 | 5S rna | | 1 | 22610.57844 | 47062.82484 | 0 | 0 |
| 105 | 88965 | 89038 | + | tRNA | tRNA-Arg-ACG | | + | 88965 | 1 | tRNA-Arg ACG | | 1 | 259.5558267 | 1603.139018 | 0.625059185 | 0.282367066 |
| | | | | | | | + | 89190 | | intergenic | | 1 | 23.43971945 | 29.22499135 | 0 | 0 |
| 106 | 89808 | 89737 | - | tRNA | tRNA-Asn-GUU | | – | 89808 | 1 | tRNA-Asn GUU | | 3 | 3.125295926 | 3.247221261 | 0.468794389 | 1.83538593 |
| | | | | | | | – | 89931 | -123 | tRNA-Asn GUU | | 2 | 3.437825519 | 0.988284732 | 3.281560723 | 3.388404794 |
| | | | | | | | – | 90537 | | intergenic | | 3 | 0 | 0 | 0.937588778 | 1.83538593 |
| | | | | | | | – | 91974 | | intergenic | | 2 | 0 | 0.282367066 | 67.03759762 | 55.20276144 |
| | | | | | | | – | 92494 | | intergenic | | 1 | 0.312529593 | 0 | 3.750355112 | 4.659056592 |
| 107 | 92927 | 93277 | + | gene | rpl21 | 50S ribosomal protein L21 | + | 92933 | | rpl21 inter | | 1 | 13.90756687 | 16.37728984 | 0 | 0 |
| 108 | 93358 | 93567 | + | gene | rpl32 | ribosomal protein L32 | | | | | | | | | | |
| 109 | 93655 | 94521 | + | gene | cysT | sulfate transport protein | | | | | | | | | | |
| 110 | 94741 | 94820 | + | tRNA | tRNA-Leu-UAG | | + | 94739 | -2 | tRNA-Leu UAG | | 1 | 52.81750116 | 118.4529843 | 0.156264796 | 0 |
| 111 | 94947 | 95909 | + | gene | ccsA | cytochrome c biogenesis protein | + | 94887 | -60 | ccsA | | 3 | 1.250118371 | 0.423550599 | 0 | 0 |
| | | | | | | | – | 90537 | | intergenic | | 3 | | | | |
| | | | | | | | – | 91974 | | intergenic | | 2 | | | | |
| 112 | 95246 | 95325 | – | | tRNA-Pro pseudo | | – | 94697 | | tRNA-Pro pseudo | | 1 | 0.156264796 | 0.141183533 | 20.47068832 | 53.22619197 |
| | | | | | | | + | 96387 | 1 | ndhD internal | | 1 | 3.750355112 | 4.517873059 | 0.156264796 | 0 |
| | | | | | | | + | 97067 | | ndhD internal | | 1 | 3.281560723 | 3.953138926 | 1.71891276 | 0.847101199 |
| 113 | 97628 | 96129 | - | gene | ndhD | NADH dehydrogenase subunit 4 | – | 97660 | -44 | ndhD (maybe -/+) | | 1 | 0 | 0 | 6.094327057 | 7.906277853 |
| 114 | 98022 | 97777 | - | gene | psaC | photosystem I subunit VII | – | 98066 | -44 | psaC (maybe -/+) | | 3 | 0 | 0 | 2.187707149 | 1.270651798 |
| 115 | 98551 | 98249 | - | gene | ndhE | NADH dehydrogenase subunit 4L | – | non detected | | | | | | | | |
| 116 | 99180 | 98605 | - | gene | ndhG | NADH dehydrogenase subunit 6 | – | non detected | | | | | | | | |
| 117 | 99821 | 99270 | - | gene | ndhI | NADH dehydrogenase subunit I | – | non detected | | | | | | | | |
| | | | | | | | + | 100895 | | ndhA internal | | 2 | 7.813239816 | 4.800240125 | 3.281560723 | 2.823670662 |
| 118 | 101685 | 99873 | - | gene | ndhA | NADH dehydrogenase subunit 1 | – | 102042 | -357 | ndhH inter (ndhA) | | 2 | 0.156264796 | 0 | 5.625532668 | 5.50615779 |
| | 101685 | 101130 | | exon 1 | | | | | | | | | | | | |
| | 100423 | 99873 | | exon 2 | | | | | | | | | | | | |
| 119 | 102865 | 101687 | - | gene | ndhH | NADH dehydrogenase subunit 7 | – | 102635 | | ndhH internal | | 1 | 0.312529593 | 0 | 6.250591853 | 7.200360187 |
| 120 | 103184 | 102918 | - | gene | rps15 | ribosomal protein S15 | | | | | | | | | | |
| 121 | 104751 | 103357 | - | gene | ORF464 | | | | | | | | | | | |
| 122 | 109509 | 108112 | - | gene | chlN | photochlorophyllide reductase subunit N | – | 108471 | | chlN inter | | 2 | 0 | 0 | 76.88227979 | 74.54490547 |
| 123 | 110416 | 109547 | - | gene | chlL | photochlorophyllide reductase subunit L | – | 110471 | -55 | chlL | | 3 | 0.312529593 | 0.423550599 | 1.562647963 | 2.682487129 |
| | | | | | | | + | 110491 | | | | 1 | 53.91135473 | 57.6028815 | 0 | 0 |
| | | | | | | | + | 110100 | | | | | 0.625059185 | 0.141183533 | 0.156264796 | 0.847101199 |
| | 120314 | 110410 | | repeat_region | | Inverted Repeat A | | | | | | | | | | |
| 124 | 111222 | 111293 | + | tRNA | tRNA-Asn-GUU | | + | 111222 | 1 | tRNA-Asn GUU | | 1 | 45.31679093 | 230.8350766 | 0.156264796 | 0 |
| 125 | 112065 | 111992 | - | tRNA | tRNA-Arg-ACG | | – | 112065 | 1 | tRNA-Arg ACG | | 1 | 0 | 0 | 3.125295926 | 36.84890214 |
| 126 | 112397 | 112279 | - | rRNA | | 5S ribosomal RNA | – | 112397 | 1 | 5S rna | | 1 | 0 | 0 | 156.5773259 | 241.282658 |
| | | | | | | | – | 112401 | -3 | 5S rna | | 1 | 0 | 0 | 0 | 0.141183533 |
| 127 | 112725 | 112623 | - | rRNA | | 4.5S ribosomal RNA | – | 112727 | -2 | 4.5S rna | | 2 | 0 | 0 | 9.219622983 | 2.823670662 |
| 128 | 115650 | 112833 | - | rRNA | | 23S ribosomal RNA | – | 115718 | -68 | 23S rna | | 2 | 0 | 0 | 446.2922583 | 78.921595 |
| | | | | | | | – | 115876 | | tRNA-Ala UGC intron | | 1 | 0 | 0 | 52.19244197 | 73.98017134 |
| | | | | | | | – | 116380 | | tRNA-Ala UGC intron | | 1 | 0 | 0 | 23.90851384 | 30.21327608 |
| 129 | 116649 | 116612 | - | tRNA | tRNA-Ala-UGC | exon 1 | – | 116649 | 1 | tRNA-Ala UGC | | 1 | 0 | 0 | 3.437825519 | 6.917993121 |
| | 115844 | 115810 | | | | exon 2 | | | | | | | | | | |
| 130 | 117589 | 117553 | - | tRNA | tRNA-Ile-GAU | exon 1 | – | 117589 | 1 | tRNA-Ile-GAU | | 1 | 0 | 0 | 2.187707149 | 9.741663783 |
| | 116760 | 116726 | | | | exon 2 | | | | | | | | | | |
| 131 | 119308 | 117810 | - | rRNA | | 16S ribosomal RNA | – | 119308 | 1 | 16S rna | | 2 | 0 | 0 | 25.00236741 | 15.24782157 |

| | | GROUP 1+2 | GROUP 3 |
|---|---|---|---|
| gTSS | | 99 | 9 |
| iTSS | | 39 | 1 |
| aTSS | | 21 | - |
| oTSS | | 15 | 2 |
| TOTAL | | | 186 |

| | | |
|---|---|---|
| TSS group 1 | A 5' end with: i) a coverage in both TEX+/TEX- libraries of at least > 2 per million mapped reads, ii) a start at the same genomic position (nucleotide) in both libraries and iii) an enrichment > 1 in the TEX+ library | 109 |
| TSS group 2 | A 5' end that was not enriched in TEX+ libraries was accepted as a TSS if it extended into an annotated gene | 65 |
| TSS group 3 | 5' ends that do not fall into the above categories but they extended into an annotated gene and a PEP promoter motif was predicted using MEME (Bailey et al. 2015). | 12 |
