## Supplementary material for "Comparative analysis of early divergent land plants and construction of DNA tools for hyper-expression in *Marchantia* chloroplasts": transcription start sites

| | Start | End | strand of gene | | Gene name | Gene function | starnd of trascript | Genomic coordinates of TSS | TSS relatively to ATG | Transcipt relatively to gene | starnd of trascript | 51 bp upstream potential TSS - **WITH BOLD PROMOTERS WITH PREDICTED MOTIFS** |
|---|---|---|---|---|---|---|---|---|---|---|---|---|
| 1 | 93 | 854 | + | | rps12 | ribosomal protein S12 | + | 1 | | | + | aaaaagaaaaataattgattgaatttaagaactaatatgttataaactaca |
| | 93 | 324 | | exon 1 | | | + | 111 | | rps12 intron 1 | + | ACGATTTAGGTAACTTATTTGTCAACTTTTCCACTACAACACCAAAAAAAC |
| | 829 | 854 | | exon 2 | | | + | 496 | | rps12 exon2 | + | GGCTATATGTATATAAAATAAAATATAATCTATATTATATACTATACAATA |
| 2 | 904 | 1371 | + | gene | rps7 | 30S ribosomal protein S7 | + | 1303 | -209 | rps7 inter (ndhB) | + | CTTTTAAACTTAGTTATGAATTAATTGATGCAGCTAGAGATAATGGAATTG |
| 3 | 1512 | 3535 | + | gene | ndhB (ndh2) | NADH dehydrogenase subunit 2 | + | 1696 | | ndhB exon 1-inter | | catctccttaacaagtttactaataagcataatattattatttcaata |
| | | | | | | | + | 2209 | | ndhB exon 1-inter | + | TATTTGTATTCTTGTAGGACTCGCATTTAAACTTTCTTTAGTTCCATTTCA |
| | 1512 | 2237 | | exon 1 | | | + | 2756 | | ndhB exon 2 -inter | + | CATGCACGGTTTTGAATGAGAGAAAAAAGAATTTTTTTTTCGACTCTAACT |
| | 2756 | 3535 | | exon 2 | | | + | 3090 | | ndhB exon 2 -inter | + | TTTTTTCTACATTTTTATGAATTTAGGAACATTTGCTTGTATTATATTATA |
| | | | | | | | + | 3192 | | ndhB exon 2 -inter | + | AAAAGATCCTTTATTAAGTTTTTCCTTAACATTATGCTTATTATCTTTAGG |
| 4 | 3633 | 3712 | + | tRNA | tRNA-Leu-CAA | | + | 3610 | -23 | tRNA-Leu | + | AAAACTTTATATCTATATCTATATATATATCTATAGATATATATATAGATA |
| | | | | | | | + | 3633 | 1 | tRNA-Leu | + | TATATATCTATAGATATATATATAGATATAGATATTATGTAGTAACTAGAG |
| 5 | 3954 | 4058 | + | gene | psbM | photosystem II protein M | + | 3775 | -179 | psbM | + | TTTTAGTTAAATTTTTTAGATAAATATTTTTTATGTTATACTATTCTATTG |
| | | | | | | | + | 3954 | | psbM-0 | + | ATCAGATGATATTTTTAATATTTTAAATAACCCTAATATAATATAATTA |
| 6 | 4182 | 5073 | + | gene | ycf66 | hypothetical protein | | | | | | |
| | 4182 | 4287 | | exon 1 | | | | | | | | |
| | 4772 | 5073 | | exon 2 | | | | | | | | |
| 7 | 5202 | 5113 | - | gene | ycf (ORF29) | | – | 5299 | | orf29 inter | – | ATTAGCAAAAAAATATCAAAAATTTTATAAAAAAAATATATAATATATATA |
| | | | | | | | – | 5406 | -204 | orf29 | – | ATAGCCTGATACGTTTGAAATGTTTTTTAATTTATTATATATTATAGACAACC |
| 8 | 5601 | 5531 | - | tRNA | tRNA-Cys-GCA | | – | 5601 | 0 | tRNA-cys | – | gccttgaattttaaaatatacattgtattatacaattaattaaaataaag |
| | | | | | | | – | 5613 | -12 | tRNA-cys | – | AATATTCAATATGCCTTGAATTTTAAAATAATACATTGTATTATACAATTA |
| 9 | 5740 | 8937 | + | gene | rpoB | RNA polymerase beta subunit | + | 5405 | -191 | rpoB? | + | AAAGAAATAATTAACTTGCTTTTTTTTTTTTATTGTTTTAATCTTATGATG |
| | | | | | | | – | 6111 | | rpoB-internal | – | gtatatataggattccattatgatctaattctgaattataataaattcca |
| | | | | | | | – | 6574 | | rpoB-internal | – | TATTCCAAATTTTAATTTGATTAAATAATCAACAGCTGCTAAAATATCTTG |
| 10 | 8968 | 11627 | + | gene | rpoC1 | RNA polymerase beta' subunit | + | 8579 | | rpoB-internal,rpoC1 | + | TTTTGAACAACCTATAACTATTGGAAAAGCTTATATGTTAAAATTAATTCA |
| | 8968 | 9399 | | exon 1 | | | – | 8887 | | rpoB-internal | – | TTAAATTTCTTTTAATTTTAATTTCAAATTTTTTTCACATATAATAACATG |
| | 10005 | 11627 | | exon 2 | | | – | 9417 | | rpoC1 intron 1 | – | TTAAAACTAAATTAAAATAACAGTTATTTATATCTGTTAAAATCAAAATCA |
| | | | | | | | – | 9757 | | rpoC1 intron 1 | – | ATTTGTTTATAGATGTTTATTTCTATATATTTTTTATTATATATATATTTA |
| 11 | 11700 | 15854 | + | gene | rpoC2 | RNA polymerase beta'' subunit | + | 12024 | rpoC1 intron 1 | rpoC2-inter | + | ACGCAGTAGAAAAATTGCGTCAACTAATAGAAACATGGTATGCTACAAGTG |
| | | | | | | | – | 12453 | | rpoC2-inter | – | AAGTGCGCCGATATCTTGATTTCGTGGAGCAATACATCTATGATCTATATA |
| | | | | | | | – | 13106 | | rpoC2-inter | – | ACAGATAATTCATTGTTTTTATTATGAAAATTTCCTGATAATATCCATATA |
| | | | | | | | – | 13985 | | rpoC2-inter | – | AATAAATCAAAAAGAGTTGATTTATTTAATTTTTTAGATATTTTATATTCA |
| 12 | 15937 | 16644 | + | gene | rps2 | 30S ribosomal protein S2 | + | 14817 | -1120 | rpoC2-inter (rps2?) | + | AAAAAATTAATAAATTGATTTTAGGAAAACATATAAATTATAATTTATTTA |
| 13 | 16773 | 17519 | + | gene | atpI | ATP synthase CF0 A subunit | + | 16700 | -73 | atpI | + | ATAAAAAATCATTACTACTTTCTAATAAAAAAATAGATTAAAATAATAACA |
| 14 | 17889 | 18134 | + | gene | atpH | ATP synthase CF0 C chain | + | 17748 | -141 | atpH? | + | ATTTGAATTTTTTCTTTTAAAAATTTAGTATTTGAAATTTAAAAAAAATATT |
| 15 | 18356 | 19483 | + | gene | atpF | ATP synthase CF0 B subunit | | | | | | |
| | 18356 | 18500 | | exon 1 | | | | | | | | |
| | 19074 | 19483 | | exon 2 | | | – | 19218 | | atpF-internal exon2 | – | CTGCATTAATTAAATCTTGTTTTTCTTTTTCCATTTGAGATAATCCATTTA |
| 16 | 19526 | 21049 | + | gene | atpA | ATP synthase CF1 alpha subunit | | | | | | |
| | | | | | | | – | 20068 | | atpA-internal | – | GCTACATAAACACATACTACATTTTGACCTTTTTGATTTAAAATAGTATCA |
| 17 | 21184 | 21113 | - | tRNA | tRNA-Arg-UCU | | – | 21184 | 1 | tRNA-arg-UCU | – | ATTTGAAATGATAAACTTAAAAAAATTTATCATTTCAAATTCTCTTTTAAG |
| | | | | | | | – | 21295 | 1 | tRNA-Gly-UCU exon 2 | – | TCACGTTCGGTTTTGAATTAGAGGCATTTTTTATAAATGTCGACTATAACC |
| 18 | 21899 | 21877 | - | tRNA | tRNA-Gly-UCC | exon 1 | – | 21899 | 1 | tRNA-Gly-UCC | – | GTTAAATAATATGATAAAAAACCTAAAATATAGTATATAAATAATTATATAG |
| | 21295 | 21248 | | | tRNA-Gly-UCC | exon 2 | + | 21926 | | intergenic | + | CTTTTACCACTAAACTATACCCGCTATATAATTATTATATATACTATATTTA |
| 19 | 22116 | 22015 | - | gene | ycf12 | | – | 22108 | | ycf12 internal | – | AAAAATTTAGAATTGTAAAAAAAAAAGAGAGGTTAAACCAGAATGAATTTA |
| | | | | | | | – | 22231 | | ycf45 internal | – | AGTATGAAATTAATTTTGAATAAAGAATATCGTCTTGTTATTATTGTACTG |
| 20 | 22278 | 22141 | - | gene | ycf45 | | – | 22317 | -39 | ycf45 | – | TTTTCTCCTTAATCATTAAAAAATAAAAAAAATATATATATATATACAATTA |
| 21 | 22372 | 22470 | + | gene | psaM | photosystem I protein M | + | 22246 | -126 | psaM? (ycf45 inter) | + | CAATAAAAAAATCTATAGCGATAATAAACTGAGATCAGTACAATAATAACA |
| | | | | | | | + | 22654 | -64 | tRNA-Ser GUC | + | ATAAAAAAAAAGATTGAAATAAATATTTTTTTTATAGTATAATTAATATA |
| 22 | 22718 | 22805 | - | tRNA | tRNA-Ser-GCU | | + | 22718 | 1 | tRNA-Ser GUC | + | ATATATACAATTTTTATTTTTATTCAAAAACAAATAAATTTTTATATTTG |
| 23 | 22933 | 22823 | - | gene | psbI | photosystem II protein I | – | 22976 | -43 | psbI | – | CCCTTTTTCTCCATTAAATTTAATGAAAAAAAATTTGTAAAATTTGAATT |
| 24 | 23430 | 23263 | - | gene | psbK | photosystem II protein K | – | 23515 | -85 | psbK | – | AAAAATTATAACCTCTTTAAAGCTCAAAAAACTTGTAGTAAGATGTACCTA |
| 25 | 23629 | 23700 | + | tRNA | tRNA-Gln-UUG | | + | 23620 | -9 | tRNA-Gln UUG | + | TTTTAATAAATAGATTGACAAAATAAAAAAATAATGATTACTAGAATTA |
| | | | | | | | – | 23780 | | intergenic | – | TATTTATATTATGACAATAAGATAAATATGGCAAGGATTTTTCTATGTTG |
| | | | | | | | + | 23845 | | intergenic | + | AATAAGATAAATATGGCAAGGATTTTTCTATGTTGTAAAATATAAAAATA |
| 26 | | | | | petD?? | | + | 24369 | -217 | petD | + | TTATCTTTAACAGAGAAACCATCAGCAAATATTATTGGTATATTTACATTA |
| 27 | 25563 | 25715 | + | gene | orf50 | | + | 25168 | -395 | ycf ORF50? | + | TAGGCTATGAAGGTACTAATCAAATAGCAGATTTAGTTTATAATTCATTTA |
| 28 | 27948 | 27982 | + | | tRNA-Lys-UUU | | + | 25787 | -47 | tRNA-Lys UUU | + | CTAATTAAAATAGAATTGACAATACTAATCCACATACATATAATTGTGTTG |
| | | | | | | | – | 25997 | | intergenic | – | TTATCCATTAACTTTAATTGACTTATTCTATGAAATTTTATTTAATTAAAA |
| 29 | 28128 | 29189 | + | gene | psbA | photosystem II protein D1 | + | 28074 | -54 | psbA | + | ATTTGAATATTAGTTGACATAATCATATGTTATGTGTAATACTATAAGTTA |
| | | | | | | | + | 28286 | | psbA internal | + | ATGATTCCTACTTTATTAACAGCAACTTCAGTATTCATTATTGCTTTTATT |
| 30 | 29353 | 29427 | + | tRNA | tRNA-His-GUG | | | | | | | |
| | | | | | | | – | 33822 | | integenic | – | ATATTCATCCAACTTTCTGTTATAACATCAGGTTTATTATATAATAGTTTG |
| | | | | | | | + | 33844 | | intergenic | + | ATGTTCCGTTATTCAAAATTTCGATAAACAAACTATTATATATATAAACCTG |
| 31 | 36152 | 36079 | - | tRNA | tRNA-Asp-GUC | | – | 36152 | 1 | tRNA-Asp | – | tattatgtcaaacatttttaacaatataaaaaaaaatgcatttatctaatg |
| 32 | 36313 | 36232 | - | tRNA | tRNA-Tyr-GUA | | – | 36180 | -28 | tRNA-Asp | – | AATAATTTATAGAAAAAAGTTGCCGTTTATTATGTCAAACATTTTTAACA |
| 33 | 36456 | 36384 | - | tRNA | tRNA-Glu-UUC | | – | 36466 | -10 | tRNA-Glu | – | TTTCTAAAAAACTATTGACAGGAAATCATATATGAGTAAGATAAAGTTA |
| 34 | 36680 | 37798 | + | gene | cysA | sulfate ABC transporter protein | + | 36361 | -319 | CysA | + | CCCAGATAGAAAAATTTTTTCTACCTGAAAAAAAGTATAAAATATTTATTA |
| | | | | | | | | 36662 | -18 | CysA | | TTAAACTTAAATAAAACATTTTTTATTTAAATATTGTTTAAAGTAGTTTTA |
| | | | | | | | + | 38004 | -36 | tRNA-Thr GGU | + | GTTTTTTTAATTAAGTTGACAATTTTGTAACTTTTGTTACACAATATGCTG |
| 35 | 38040 | 38111 | + | tRNA | tRNA-Thr-GGU | | + | 38040 | 1 | tRNA-Thr GGU | + | ttacacaatatgctgtgttaaaaaaaaaaaaaaaatagtttcttcttattg |
| 36 | 38539 | 39600 | + | gene | psbD | photosystem II protein D2 | + | 38291 | -248 | psbD | + | AAAAAAAAATTGGAACTTTCAACTTTCTATTTACTATGATAAAATAAATTA |
| | | | | | | | + | 38395 | -144 | psbD | + | AAACTATTAGATAATGAATTTAAAATAAAAAAGACATAAAAAAATTATTAC |
| 37 | 39548 | 40969 | + | gene | psbC | photosystem II 44 kDa protein | + | 39541 | -7 | psbC | + | AAAATATTTTATTAAATGAAGGTATTAGAGCTTGGATGGCAGCTCAAGATC |
| 38 | 41178 | 41091 | - | tRNA | tRNA-Ser-UGA | | – | 41178 | 1 | tRNA-ser | – | TTTTTATAATTTTTTAATAAAAAATTATTAGATTAAAAATGTTTACTATAG |
| 39 | 41354 | 41542 | + | gene | psbZ | photosystem II protein Z | + | 41325 | -29 | psbZ | + | TAATTTTTTGATAATTTTACAAAAACAGGATTTGATGGTAATATTTTCATA |
| | | | | | | | – | 41589 | | intergenic | – | TTTATCTAAAGCACTTGTTTCAAATGCATTTAGAATTTATAATTAATGTA |
| 40 | 41747 | 41817 | + | tRNA | tRNA-Gly-GCC | | + | 41747 | 1 | tRNA-Gly GCC | + | TAAAAAAAAATATATATTATATATATATAATAAGTATGTATAATTTATATACG |
| 41 | 41947 | 41874 | - | tRNA | tRNA-fMet-CAU | | – | 41947 | 1 | tRNA-fMet-CAU | – | ATCTTGCTTAATTCTATTTTTATAGAGTATACTTTTTTCGTTAATTATAAC |
| | | | | | | | – | 41958 | -11 | tRNA-fMet-CAU | – | CTTTAAAATATATCTTGCTTAATTCTATTTTTATAGAGTATACTTTTTTCG |
| 42 | 42353 | 42051 | - | gene | rps14 | 30S ribosomal protein S14 | – | not detected | | | | |
| 43 | 44646 | 42442 | - | gene | psaB | PSI P700 apoprotein A2 | – | not detected | | | | |
| | | | | | | | + | 45424 | | psaA internal | + | TACTACCAACTGCTATTACATCTCCACCACCCCATGTTAAACTAGTACTTG |
| 44 | 46925 | 44673 | - | gene | psaA | PSI P700 apoprotein A1 | – | 47074 | -149 | psaA | – | ATAAAATTTATGGTCCATTAAGCACCTTAAAATTGTGTCATAATAAATTTG |
| 45 | 48316 | 47207 | - | gene | pafI | photosystem I-related pafI Ycf3 | – | | | | – | |
| | 48316 | 48193 | | exon 1 | | | – | | | | – | |
| | 47586 | 47207 | | exon 2 | | | + | 47495 | | pafI inter | + | GCATGTTCTCCATTACTTGTATGAATAAGACCTATATTATATAGTATATAA |
| | | | | | | | – | 48278 | | pafI inter | – | TTTATCAATAAAATTATCATTTTTTTGAGATCTAGGCATAGTTAATTATA |
| 46 | 48563 | 48650 | + | tRNA | tRNA-Ser-GGA | | + | 48548 | -15 | tRNA-Ser GGA | + | CAAATTAAAAAATATTGCTATAATATATATATTTTCCATTAGAATATTTCCA |
| | | | | | | | + | 48563 | 1 | tRNA-Ser GGA | + | tgctataatatatatttttccattagaatatttccaagataaactatcttg |
| 47 | 49667 | 49059 | - | gene | rps4 | ribosomal protein S4 | + | 49325 | | rps4 internal | + | ACAATTATAACTTGGTATATCAACTGTATTATTATTTATTAAAATATGTCG |
| | | | | | | | – | 49456 | | rps4 internal | – | ATTACGTTTTCATTATGGATTAACCGAAAGACAGTTATTAAAATATGTACG |
| 48 | 49968 | 49896 | - | tRNA | tRNA-Thr-UGU | | – | 49968 | 1 | trnA-Thr | – | TAATGATAGATTATCAATCTGTTTTTCTAATAAATAAATTTGAATTATTTG |
| | | | | | | | – | 50041 | -73 | trnA-Thr | – | AATTAAAAAACAATTCTTGCTTTTTTAAATATATTATATATATAATATGTG |
| 49 | 50174 | 50208 | + | tRNA | tRNA-Leu-UAA | exon 1 | + | 50139 | -35 | tRNA-Leu UAA | + | ATTTTTTTTTAACTATTGCATAAAAAAAAAAAAAAACTATAATTAAATTTG |
| | | | | | | | + | | | | | |
| | | | | | | | – | 50804 | | integenic | – | ATTTTAATAATATATTGAAACAAAAATAAATGTATATATAAAATATACATA |
| | 50523 | 50572 | | | | exon 2 | + | | | | | |
| | | | | | | | | 50855 | | integenic | + | ATGTATATTTTATATATACATTATTTTTGTTTCAATATATTATTAAAATA |
| 50 | 50649 | 50721 | + | tRNA | tRNA-Phe-GAA | | | 50649 | 1 | tRNA-Phe-GAA | + | AAAGTTGACATATTTTTTCTTTTATTATGTTAATATAAAGGAAATAAAAATTG |
| | | | | | | | + | 51683 | | ndhK inter | + | TGAGCTATTTTTTTACGAAGTTTTATTATAGCATCTATAATTGCTTCTGGT |
| 51 | 51393 | 50884 | - | gene | ndhJ | NADH dehydrogenase subunit J | – | 51356 | | internal ndhJ | – | AAGTACAAATAATATGTTAAACATTTTAAAAAATAACAATAATAAAATACA |
| 52 | 52174 | 51437 | - | gene | ndhK | NADH dehydrogenase subunit K | – | **52245** | -71 | ndhK (internal ndhC | – | TTCTTTATCCTTGGGCTATGAGTTTTTATAATTTGGTATATCATCTTTTA |
| 53 | 52527 | 52165 | - | gene | ndhC | NADH dehydrogenase subunit 3 | | | | | | |
| | | | | | | | – | 52728 | | tRNA Val | – | ATGCATGTTGGGTTCTTAAAACAGTTTTTAATTTAAAAGGACTGTTTTACC |
| 54 | 53292 | 53256 | - | tRNA | tRNA-Val-UAC | exon 1 | – | 53292 | | tRNA-Val UAC | – | TATATATATATTGTATAGGAAATAATACGAATATAACTCGTCCGTATTAAA |
| | 52728 | 52694 | | | tRNA-Val-UAC | exon 2 | – | 53330 | -38 | tRNA-Val UAC-- | – | ATTGAATCTAAAACTTGACAATAAGTAATAAAAAAGTGTATATATATATTG |
| | | | | | | | + | 53430 | -10 | tRNA-Met CAU | + | TTTTTTTACTTATCTTGCATTAAAAGTTATTATATAGTTATAATAAGTTTG |
| 55 | 53440 | 53513 | + | tRNA | tRNA-Met-CAU | | + | 53440 | 1 | tRNA-Met CAU | + | tatcttgcattaaaagttattatatagttataataagtttgaaataaaata |
| 56 | 53994 | 53590 | - | gene | atpE | ATP synthase epsilon chain | | | | | | |
| 57 | 55478 | 54000 | - | gene | atpB | ATP synthase CF1 beta subunit | – | 55498 | -20 | atpB | – | TTATTAATTGATTTATTTGATACACAATATTTTTTTATTATAATTCATTA |
| | | | | | | | – | 55748 | -270 | atpB | – | TCCAATATAAAGATGTAAATAAATAAATAAATATATATATATATATATATA |
| 58 | 56027 | 57454 | + | gene | rbcL | rbcL | + | 55903 | -124 | rbcL | + | ATAAAATAACTAGGTTGCATTACATATAAAAACAATATACAATAATAATG |
| 59 | 57573 | 57646 | + | tRNA | tRNA-Arg-CCG | | + | 57573 | 1 | tRNA-Arg CCG | + | ttttatctgatatttcagataaaaaaaagaaaattacaaaacttttaaatg |
| 60 | 57831 | 58778 | + | gene | accD | | + | 57700 | -131 | accD? | + | TTTATTTGAAAATTTTTGCATAAAAAAAAAATTGTTCTATTATTAAATTA |
| 61 | 58954 | 59064 | + | gene | psaI | psaI | + | 58880 | -74 | psaI | + | TTTTATATTTTGTTATTGTTATTTATTAAAATAAGATATAATTTTTATTA |
| 62 | 59287 | 59841 | + | gene | pafII | photosystem I assembly protein Ycf4 | + | | | | | |
| | | | | | | | + | 59326 | | pafII inter | + | ATTCTTTTGTTATGAATTTACAAGTGGACCATATTAGAGTAGATTTTATAA |
| | | | | | | | + | 59572 | | pafII inter 2/cemA? | + | ATTTATGGTGCACTATTTGTTGGAATGTCGGTAGTGGCTATAATAAATTTG |
| 63 | 59913 | 61217 | + | gene | cemA | envelope membrane protein | + | | | | | |
| 64 | 61403 | 62365 | + | gene | petA | apocytochrome f precursor | + | 61012 | | cemA inter - petA? | + | AAGCGATACAATGAAAGCTTTTTTTATTCTTTTATTAACTGATTTATGTAT |
| 65 | 62680 | 62558 | - | gene | psbJ | PSII reaction center subunit X | – | 62708 | -28 | psbJ | – | AGACTTTTTTTATTATTTGTAACTGTTTATGTTTTGAGTTATAACTATTTA |
| 66 | 62916 | 62800 | - | gene | psbL | PSII reaction center subunit XII | – | no detected | | | | |
| 67 | 63057 | 62938 | - | gene | psbF | photosystem II reaction center subunit VI | – | 63270 | -213 | psbF / psbE interna | – | CCATGTCTGGAAATACGGGAGAGCGTCCTTTTGCTGATATAATTACCAGTA |
| 68 | 63318 | 63067 | - | gene | psbE | cytochrome b559 alpha subunit | – | 63460 | -142 | psbE | – | TTTTTTGAATTTATCTTGCTGCGTAAAAAGAACATTAGCTATACTAAGTTA |
| 69 | 63917 | 64012 | + | gene | petL | | + | 63772 | -145 | petL | + | TTATTTTAAATAATTTTTGTCAAAATAATATATATATATTAATATATATTA |
| | | | | | | | + | 63876 | -41 | petL 2? | + | CTTTATATATAAAAAAACCATTTTTATTTGAAAATTTTATATAATAATTTG |
| 70 | 64134 | 64247 | + | gene | petG | cytochrome b6/f complex subunit V | + | 64088 | -46 | petG | + | ATTTCATTGTATTTCTCAAACTTTTTTTGAGATTCATAGTAAACTACAATA |
| 71 | 64391 | 64318 | - | tRNA | tRNA-Trp-CCA | | – | 64391 | 1 | tRNA-Trp CCA | – | tatctatgtatataaataaaaatagaattcaaattaatactaattaataag |
| | | | | | | | + | 64445 | | | + | TTAATTAGTATTAATTTGAATTCTATTTTTATTTATATACATAGATATATA |
| 72 | 64544 | 64471 | - | tRNA | tRNA-Pro-UGG | | – | 64544 | 1 | tRNA-Pro UGG | – | TAGGCAAAGAAAAAAAAAAAAAGTACAATACAGGTAGAGAAAAAAGGTTTA |
| | | | | | | | – | 64559 | -15 | tRNA-Pro UGG | – | AGTAAGTAAAATAAGTAGGCAAAGAAAAAAAAAAAAAGTACAATACAGGTA |
| | | | | | | | – | 64709 | -165 | tRNA-Pro UGG | – | TTATAACATTTGTATTTTTATATATATTATATATAGTATATACTACACATA |
| 73 | 64780 | 64908 | + | gene | psaJ | photosystem I subunit IX | + | 64732 | -48 | psaJ | + | TTTGTTTATTAATCTAGTTAACATAATTATGTGTAGTATATACTATATATA |
| 74 | 65027 | 65224 | + | gene | rpl33 | ribosomal protein L33 | + | 64741 | -39 | psaJ 2/ rpl33? | + | TAATCTAGTTAACATAATTATGTGTAGTATATACTATATATAATATATATA |
| 75 | 65252 | 65479 | + | gene | rps18 | ribosomal protein S18 | + | 65033 | -219 | rpl33 inter (rps18) | + | ATTTTTTTAAAAGGTGGTATAAACCTAAAATTTCAAATAGAATTATGGCTA |
| 76 | 65909 | 65559 | _ | gene | rpl20 | 50S ribosomal protein L20 | – | not detected | | | | |
| | | | | | | | – | 66272 | | rps12 inter | – | ATCAGAAATTATACAAATAACTGATATTATAAATATATATTATATATTACA |
| 77 | 66199 | 67098 | – | gene | rps12 | | – | 66447 | | rps12 intron | – | CTTAAGCGAGAAATTTAATTAAATAAATTATTTTGCATAATATAATATTACTTA |
| | | | | | | | – | 66856 | rps12 | -47 | – | AATTCTACTTCTATTAAAAATTAGTTTTTAAACAAAAAAATTTTTATTTGT |
| 78 | 68388 | 66883 | - | gene | clpP | ATP-dependent Clp protease proteolytic subunit | – | 67134 | clpP inter | clp exon 3 | – | GTTTCATTTATTTTTTTAATAAAAAAATAGAAAATTTAATAATTATTAATA |
| | 68388 | 68318 | | exon 1 | | | – | 67805 | clpP inter 2 | clp exon 2 | – | AAAAGCGCTTGTACAGTTTTATAAGAAAAAAATAATAAAATTATCTTAATC |
| | 67805 | 67514 | | exon 2 | | | – | 68477 | clpP | -59 | – | TATTTATAAAATTGTATTCTATTTTATAGAAAAACTATATATAATAAGTAG |
| | 67134 | 66883 | | exon 3 | | | | | | | | |
| 79 | 68717 | 70243 | + | gene | psbB | photosystem II 47 kDa protein | 1 | 68573 | -144 | psbB | 1 | GACCAATCTCTCCATTGTTATCCTAAACATAGAAATGCTAAACTGTTTGTG |
| 80 | 70361 | 70468 | + | gene | psbT | photosystem II protein T | + | | | | + | |
| 81 | 70665 | 70534 | - | gene | psbN | photosystem II protein N | – | 70712 | -47 | psbN | – | AAAACTTATCAGATTAGTTGAGATTTATTAACTTTTTGTACTATTATATTA |
| | | | | | | | – | 74208 | | rpoA internal | – | TCAACAATAATAGGTTTACAAGAATCTATTCATGATATATTAATTAATTTA |
| 82 | 70763 | 70987 | + | gene | psbH | photosystem II protein H | + | 70726 | | psbH | + | TTCCAAAATAATGGTATATTTTTAATGTATATTTTTTAATATAATAGTACA |
| 83 | 71092 | 72222 | + | gene | petB | | + | 71049 | | petB | + | TAAAAAAATGAATTGCTGCTAAAATAGCAGCAATTCATTTTTTTATTTTAG |
| | 71092 | 71097 | | exon 1 | | | | | | | | |
| | 71575 | 72222 | | exon 2 | | | | | | | | |
| 84 | 72369 | 73339 | + | gene | petD | | + | 71585 | | petB exon 2 interna | + | ATCATTTACGGTTTTTCGAGGGGGAACTTTAGTAACCTATCTCAATAAAGT |
| | | | | | | | + | | | | | |
| | 72369 | 72376 | | exon 1 | | | | | | | | |
| | 72865 | 73339 | | exon 2 | | | | | | | | |
| 85 | 74480 | 73455 | + | gene | rpoA | RNA polymerase alpha subunit | – | | | | | |
| 86 | 74905 | 74513 | - | gene | rps11 | 30S ribosomal protein S11 | – | 74917 | -12 | rps11 | – | CAAAGACAAGGTTAAAAAAGTTTAAATAAAAAAACATATAAAATATATACA |
| 87 | 75066 | 74953 | - | gene | rpl36 | 50S ribosomal protein L36 | – | | | | | |
| 88 | 75337 | 75101 | - | gene | infA | translation initiation factor 1 | – | not detected | | | | |
| 89 | 75821 | 75423 | - | gene | rps8 | ribosomal protein S8 | – | not detected | | | | |
| 90 | 76271 | 75903 | - | gene | rpl14 | 50S ribosomal protein L14 | – | not detected | | | | |
| 91 | 77310 | 76367 | - | gene | rpl16 | ribosomal protein L16 | – | | | | | |
| | | | | | | | + | 77341 | | integenic | + | ACGAGTCACACACTAAGCATAGCAATTTTTTTTATGTTAAATTTATAATTA |
| 92 | 78021 | 77365 | - | gene | rps3 | 30S ribosomal protein S3 | – | 77713 | | rps3 internal | – | AGTAGAGGTCAAGGAATTGAACAATTAAAATTAAATGTACAAAATATATTA |
| | | | | | | | + | 78105 | | rpl22 | + | ATTTTTTTCAATTTTTTTTATTTAGGAATAATATTCAGTACAATAGTTATA |
| 93 | 78431 | 78072 | - | gene | rpl22 | ribosomal protein L22 | – | not detected | | | | |
| 94 | 78727 | 78449 | - | gene | rps19 | 30S ribosomal protein S19 | – | not detected | | | | |
| | | | | | | | – | 80008 | | rpl12 int | – | ATGAAATAGTTAAGTGTCAGCCACAAAAAAAATAACATATAATAAACATA |
| 95 | 80122 | 78765 | - | gene | rpl2 | 50S ribosomal protein L2 | – | 80190 | -68 | rpl23 inter (rpl2) | – | AACAGGATATACTGTTCGTTATAAACGAATGATTATAAAATTGCAATCTGG |
| 96 | 80434 | 80159 | - | gene | rpl23 | ribosomal protein L23 | – | 80473 | -39 | rpl23 | – | TAAAGTCAAAAATTCTATACAGTATCTAAAAAAATTTATAAATTATAGGAA |
| 97 | 80670 | 80597 | - | tRNA | tRNA-Ile-CAU | | – | 80670 | 1 | tRNA-Ile CAU | – | AACTATGATGATCCAATGAATATTTATTATTCATAAATAATATATAGAAAG |
| | | | | | | | – | 80711 | -41 | tRNA-Ile CAU | – | AAGAAAAATAATTGATTGAATTTAAGAACTAATATGTTATAAACTATGATG |
| | 80716 | 90620 | | repeat_region | | inverted repeat B | | | | | | |
| 98 | 81427 | 81498 | + | tRNA | tRNA-Val-GAC | | + | 81427 | 1 | tRNA-Val GAC | + | atatattttttagaagtcagattttaagttatctttttttttctgagaata |
| 99 | 81722 | 83220 | + | rRNA | 16S ribosomal RNA | | + | 81572 | -150 | 16S rna -- | + | TGTTGTTTTCTTGTTATTGAAAGAGGCTTGTGGGATTGACATAATAGGGTA |
| | | | | | | | + | 81593 | -129 | 16S | + | AGAGGCTTGTGGGATTGACATAATAGGGTAGGTATGGGTATACTAGAAATG |
| | | | | | | | + | 81687 | -35 | 16S rna - | + | ATCTATCTCTTGGGATGGAAGACGATTTGAAATCTGCTTTGTTTACGAAAA |
| | | | | | | | + | 81722 | 1 | 16S rna | + | GCTTTGTTTACGAAAAGGAAGCTATAAGTAAAAGTAATATAACTATGAAT |
| | | | | | | | + | 83225 | | 16S internal | + | GTAACAAGGTAGCCGTACTGGAAGGTGCGGCTGGATCACCTCCTTTTTAGG |
| 100 | 83441 | 83477 | + | tRNA | tRNA-Ile-GAU | exon 1 | + | 83441 | 1 | tRNA-Ile GUA | + | GTTTTTTTCGCCCTTGGTATTGCAGGTCTCTTAGGAGACCTGCACGACGGG |
| | 84270 | 84304 | | | | exon 2 | + | 84305 | -76 | tRNA-Ala UGC | + | GATGATTTACTTCACGGGCGAGGTCTCTGGTTCAAGTCCAGGATAGCCCAG |
| 101 | 84381 | 84418 | + | tRNA | tRNA-Ala-UGC | exon 1 | + | 84381 | 1 | tRNA-Ala UGC | + | ataatgatttatttttgcatgctttccttgctttctttggcctgggtatag |
| | 85186 | 85220 | | | | exon 2 | + | 84650 | | tRNA-Ala UGC intron | + | TTGGTTAGATCTAATATGGATCGTACATGGACGATAGTTGGAGTCGGCGGC |
| | | | | | | | + | 85154 | | tRNA-Ala UGC intron | + | GTTTGGGGGGGAGTTATTGTCTATCAAAGGCCTCTATGGTAAAATAAATCA |
| 102 | 85380 | 88197 | + | rRNA | | 23S ribosomal RNA | + | 85312 | -68 | 23S rna- | + | AATAAAAAAAATTTTCATCTTTATAACCAAGTTGATCTAATTTCTTATTAT |
| 103 | 88305 | 88407 | + | rRNA | | 4.5S ribosomal RNA | + | 88302 | -3 | 4.5S rna | + | AGATTTTCTGTCGTTGCAAGAAATGGAACGACAAAAGTCTTGAGAATCCAA |
| 104 | 88633 | 88751 | + | rRNA | | 5S ribosomal RNA | + | 88630 | -3 | 5S rna | + | TTACAATTACTTGGTTTTTTTTTATTATACTTTTCTTTGTTCATGGGTTGA |
| | | | | | | | + | 88633 | 1 | 5S rna | + | CAATTACTTGGTTTTTTTTTATTATACTTTTCTTTGTTCATGGGTTGATAT |
| 105 | 88965 | 89038 | + | tRNA | tRNA-Arg-ACG | | + | 88965 | 1 | tRNA-Arg ACG | + | gagaaaagattgcttttggaaaaggcttctagaacagattagtggaggcag |
| | | | | | | | + | 89190 | | intergenic | + | TACTAAAAATCATACATTACCTACTCTTTATGTATAGTACACTTAATATTA |
| 106 | 89808 | 89737 | - | tRNA | tRNA-Asn-GUU | | – | 89808 | 1 | tRNA-Asn GUU | – | taattttaggttgaaaagtttttgaatatttttagaattaaactcattgtt |
| | | | | | | | – | 89931 | -123 | tRNA-Asn GUU | – | TGTAGAGTTTAGGGGATTGCTATCGTTATCTTTTTACCCTATAATAAAAGA |
| | | | | | | | – | 90537 | | intergenic | – | TATATAGCTATATTATTTAAATAATATTTATTTATGATAACATATTTTATG |
| | | | | | | | – | 91974 | | intergenic | – | TTCAACAATTATCAAAACGATTTTTTGAATTACTAAGTTATAATCAAATTA |
| | | | | | | | – | 92494 | | intergenic | – | TCTATAAAAAAATTTCGTCGTCTATCTTCTTTTATTAGTATTATGTTTTTA |
| 107 | 92927 | 93277 | + | gene | rpl21 | 50S ribosomal protein L21 | + | 92933 | | rpl21 inter | + | TTTTATTTAAAACAAATTAATAAAATTAAAAAATTTATAACATAATGAGTA |
| 108 | 93358 | 93567 | + | gene | rpl32 | ribosomal protein L32 | | | | | | |
| 109 | 93655 | 94521 | + | gene | cysT | sulfate transport protein | | | | | | |
| 110 | 94741 | 94820 | + | tRNA | tRNA-Leu-UAG | | + | 94739 | -2 | tRNA-Leu UAG | + | TATACTCCGTATCATTTTTTTTATAAAAAAATATAGTATATAATTGAAGTA |
| 111 | 94947 | 95909 | + | gene | ccsA | cytochrome c biogenesis protein | + | 94887 | -60 | ccsA | + | AAAAATATATTTTATAATATCTTTTGTCTTTTTATTTATTTATAATGAGTA |
| | | | | | | | – | 90537 | | intergenic | – | TATATAGCTATATTATTTAAATAATATTTATTTATGATAACATATTTTATG |
| | | | | | | | – | 91974 | | intergenic | – | TTCAACAATTATCAAAACGATTTTTTGAATTACTAAGTTATAATCAAATTA |
| 112 | 95246 | 95325 | – | | tRNA-Pro pseudo | | – | 94697 | | tRNA-Pro pseudo | – | GTATCATTTTTTTTATAAAAAAATATAGTATATAATTGAAGTAAGCCGCTA |
| | | | | | | | + | 96387 | 1 | ndhD internal | + | ATAGGAGTCAAAATAATACCAATTCCTTGAATAATAATTATAATTATTTTA |
| | | | | | | | + | 97067 | | ndhD internal | + | TTTTTTATTAATTAAAAGTTGAAAATCGAAAGTAAATTCATTAGAATTATA |
| 113 | 97628 | 96129 | - | gene | ndhD | NADH dehydrogenase subunit 4 | – | 97660 | -44 | ndhD (maybe -/+) | – | AAGAATCTATATATATATATATATATATATATATATATATAGATTATATAC |
| 114 | 98022 | 97777 | - | gene | psaC | photosystem I subunit VII | – | 98066 | -44 | pscA (maybe -/+) | – | TATTTTTTAATTATATAAAAAAATAATAAAAAAAAATATATATATATATAT |
| 115 | 98551 | 98249 | - | gene | ndhE | NADH dehydrogenase subunit 4L | – | non detected | | | | |
| 116 | 99180 | 98605 | - | gene | ndhG | NADH dehydrogenase subunit 6 | – | non detected | | | | |
| 117 | 99821 | 99270 | - | gene | ndhI | NADH dehydrogenase subunit I | – | non detected | | | | |
| | | | | | | | + | 100895 | | ndhA internal | + | TGGGTTTTCTTACCTTCTACTCTTGCTTATTCAATATATATAATAAAATTA |
| 118 | 101685 | 99873 | - | gene | ndhA | NADH dehydrogenase subunit 1 | – | 102042 | -357 | ndhH inter (ndhA) | – | TCGTTATTTAGTAAGAATTGGTGAAATGAAAGAATCTGTTAAAATAATTCA |
| | 101685 | 101130 | | exon 1 | | | | | | | | |
| | 100423 | 99873 | | exon 2 | | | | | | | | |
| 119 | 102865 | 101687 | - | gene | ndhH | NADH dehydrogenase subunit 7 | – | 102635 | | ndhH internal | – | CAATATCTTCCTTACGTAACACGATGGGATTATTTAGCTACAATGTTTACA |
| 120 | 103184 | 102918 | - | gene | rps15 | ribosomal protein S15 | | | | | | |
| 121 | 104751 | 103357 | - | gene | ORF464 | | | | | | | |
| 122 | 109509 | 108112 | - | gene | chlN | photochlorophyllide reductase subunit N | – | 108471 | | chlN inter | – | TAGAAATTTCTTTAGCAAGATTTTTAATCAGATGTGGTATGATTGTTTATG |
| 123 | 110416 | 109547 | - | gene | chlL | photochlorophyllide reductase subunit L | – | 110471 | -55 | chlL | – | AATCAATTAATTCAAAAGCTCATAATAACATAAAATATGTTATCATAAATA |
| | | | | | | | + | 110491 | | | | TAAAATATGTTATCATAAATAAATATTATTTAAATAATATAGCTATATAAA |
| | | | | | | | + | 110100 | | | + | GTAGAGTTTAGGGGATTGCTATCGTTATCTTTTTACCCTATAATAAAAGAA |
| | 120314 | 110410 | | repeat_region | | Inverted Repeat A | | | | | | |
| 124 | 111222 | 111293 | + | tRNA | tRNA-Asn-GUU | | + | 111222 | 1 | tRNA-Asn GUU | + | taattttaggttgaaaagtttttgaatatttttagaattaaactcattgtt |
| 125 | 112065 | 111992 | - | tRNA | tRNA-Arg-ACG | | – | 112065 | 1 | tRNA-Arg ACG | – | GAGAAAAGATTGCTTTTGGAAAAGGCTTCTAGAACAGATTAGTGGAGGCAG |
| 126 | 112397 | 112279 | - | rRNA | | 5S ribosomal RNA | – | 112397 | 1 | 5S rna | – | CAATTACTTGGTTTTTTTTTATTATACTTTTCTTTGTTCATGGGTTGATAT |
| | | | | | | | – | 112401 | -3 | 5S rna | – | TTACAATTACTTGGTTTTTTTTTATTATACTTTTCTTTGTTCATGGGTTGA |
| 127 | 112725 | 112623 | - | rRNA | | 4.5S ribosomal RNA | – | 112727 | -2 | 4.5S rna | – | GATTTTCTGTCGTTGCAAGAAATGGAACGACAAAAGTCTTGAGAATCCAAG |
| 128 | 115650 | 112833 | - | rRNA | | 23S ribosomal RNA | – | 115718 | -68 | 23S rna | – | AATAAAAAAAATTTTCATCTTTATAACCAAGTTGATCTAATTTCTTATTAT |
| | | | | | | | – | 115876 | | tRNA-Ala UGC intron | – | GTTTGGGGGGGAGTTATTGTCTATCAAAGGCCTCTATGGTAAAATAAATCA |
| | | | | | | | – | 116380 | | tRNA-Ala UGC intron | – | TTGGTTAGATCTAATATGGATCGTACATGGACGATAGTTGGAGTCGGCGGC |
| 129 | 116649 | 116612 | - | tRNA | tRNA-Ala-UGC | exon 1 | – | 116649 | 1 | tRNA-Ala UGC | – | ATAATGATTTATTTTTGCATGCTTTCCTTGCTTTCTTTGGCCTGGGTATAG |
| | 115844 | 115810 | | | | exon 2 | | | | | | |
| 130 | 117589 | 117553 | - | tRNA | tRNA-Ile-GAU | exon 1 | – | 117589 | 1 | tRNA-Ile-GAU | – | TGGTTTTTTTCGCCCTTGGTATTGCAGGTCTCTTAGGAGACCTGCACGACG |
| | 116760 | 116726 | | | | exon 2 | | | | | | |
| 131 | 119308 | 117810 | - | rRNA | | 16S ribosomal RNA | – | 119308 | 1 | 16S rna | – | GCTTTGTTTACGAAAAGGAAGCTATAAGTAAAAGTAATATAACTATGAAT |
